# DNA origami demonstrate the unique stimulatory power of single pMHCs as T-cell antigens

**DOI:** 10.1101/2020.06.24.166850

**Authors:** Joschka Hellmeier, Rene Platzer, Alexandra S. Eklund, Thomas Schlichthärle, Andreas Karner, Viktoria Motsch, Elke Kurz, Victor Bamieh, Mario Brameshuber, Johannes Preiner, Ralf Jungmann, Hannes Stockinger, Gerhard J. Schütz, Johannes B. Huppa, Eva Sevcsik

## Abstract

T-cells detect with their T-cell antigen receptors (TCRs) the presence of rare peptide/MHC complexes (pMHCs) on the surface of antigen presenting cells (APCs). How they convert a biochemical interaction into a signaling response is poorly understood, yet indirect evidence pointed to the spatial antigen arrangement on the APC surface as a critical factor. To examine this, we engineered a biomimetic interface based on laterally mobile functionalized DNA origami platforms, which allow for nanoscale control over ligand distances without interfering with the cell-intrinsic dynamics of receptor clustering. We found that the minimum signaling unit required for efficient T-cell activation consisted of two ligated TCRs within a distance of 20 nanometers, if TCRs were stably engaged by monovalent antibody fragments. In contrast, antigenic pMHCs stimulated T-cells robustly as well-isolated entities. These results identify the minimal requirements for effective TCR-triggering and validate the exceptional stimulatory potency of transiently engaging pMHCs.

## INTRODUCTION

T-cells are activated by productive T-cell antigen receptor (TCR)-peptide/MHC (pMHC) interactions within the immunological synapse, the dynamic area of contact between a T-cell and an antigen presenting cell (APC). Despite demonstrably low TCR:pMHC affinities^1^, T-cells detect the presence of a few stimulatory antigenic pMHCs on the APC surface, where they are typically vastly outnumbered by non-stimulatory endogenous pMHCs^2,3^. Mechanisms underlying this remarkable sensitivity remain unresolved, and it is unclear how pMHC binding triggers the early intracellular signaling response. Spatial arrangement of TCRs and their pMHC ligands and assembly into organized structures has been suggested to play a decisive role in this^4,5^.

Upon ligand engagement, monomeric^6,7^ TCRs rapidly reorganize to form TCR-microclusters, which are considered hubs of the signaling response^8–10^. This led to the notion that pre-clustered ligands could facilitate T-cell activation at low antigen density by promoting TCR clustering^11–14^. Recent studies found that T-cell activation required either deliberate pre-clustering of single ligand molecules at spacings below 50 nm^15^, or their stochastic accumulation to small clusters^16^. Both studies used high-affinity ligands with long TCR dwell times, such that TCR organization mirrored ligand organization. It hence appears likely that certain spatial requirements for the organization of triggered TCRs control TCR-proximal signaling. Whether the same spatial requirements also apply for the organization of the physiological TCR ligand pMHC, however, is unclear. Indeed, there are indications to the contrary: Spatially well-separated pMHC:TCR binding events were observed to elicit T-cell activation^17^, and even in T-cell induced microclusters individual pMHCs are not in close molecular proximity^6^. Furthermore, even a single pMHC molecule on an APC surface is sufficient for activating T-cells^18^, suggesting that the high stimulatory potency of pMHC may not depend on its (pre-)clustering.

Based on these considerations we hypothesized that proximity of triggered TCRs but not necessarily of pMHCs should modulate the sensitivity of the T-cell response. To test this, we sought to engineer an APC-mimicking biointerface with dual functionality: first, it should generate defined exclusion zones around individual ligands to isolate them as they cluster during T-cell activation; second, it should permit the directed pre-organization of single ligand molecules with nanometer precision. To this end, we employed rectangular DNA origami platforms anchored to fluid planar supported lipid bilayers (SLB) and functionalized with either one of two different types of TCR ligands: high-affinity anti-TCR single chain antibody fragments (scF_V_) to serve as templates for the organization of TCRs, or low-affinity antigenic pMHCs as the natural ligand (**Fig. 1a**). In this fashion our dynamic biointerface supports experimentally controllable and defined adjustment of ligand distances on the nanoscale while permitting at the same time the microscopic reorganization of ligands and receptors in the course of T-cell activation.

**Figure 1.**
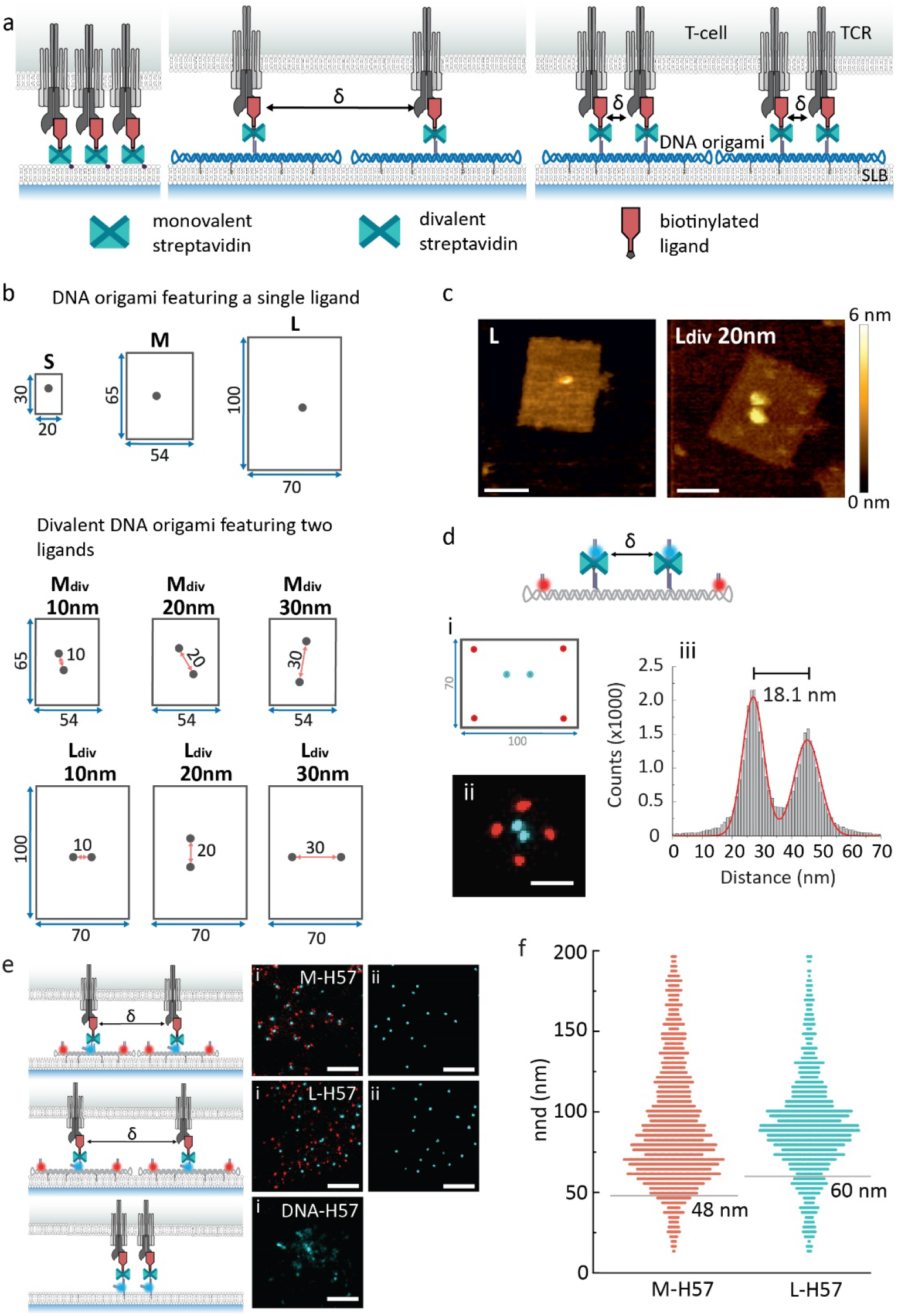
Mobile DNA origami platforms for nanoscale ligand organization. **a,** Ligands directly anchored to the SLB via mSA can rearrange freely during T-cell activation (left). DNA origami platforms functionalized with ligands at pre-defined positions and attached to the SLB via cholesterol-modified oligonucleotides set a minimum distance δ between neighboring ligands (middle). Divalent DNA origami platforms feature two ligands with a predefined δ (right). **b,** DNA origami layouts and nomenclature. DNA origami platforms of different sizes (<u>S</u>mall, <u>M</u>edium, <u>L</u>arge) were functionalized with either one (S, M, L) or two (M_div_, L_div_) dSA as indicated in the schemes. Distances are given in nm. **c,** Representative AFM images of large DNA origami platforms featuring a single ligand or two ligands spaced 20 nm apart. Scale bar, 50 nm. **d,** Mapping ligand localization on divalent DNA origami platforms via DNA-PAINT super-resolution microscopy. i, Biotinylated ligands were replaced with biotinylated DNA-PAINT docking strands. ii, Exemplary pseudo-color DNA-PAINT super-resolution image of large DNA origami platforms featuring two ligand attachment sites at 20 nm distance. Ligand (cyan) and platform (red) positions were imaged consecutively by Exchange-PAINT ^24^ using Cy3B-labeled imager strands. Scale bar, 50 nm. iii, Cross-sectional histogram of ligand positions from summed DNA-PAINT localizations of 100 individual DNA origami platforms. **e,** Representative pseudo-color DNA-PAINTsuper-resolution images of H57-decorated constructs within microclusters showing platform (red) and ligand (cyan) sites (i). ii, DNA-PAINT ligand positions after post-processing. DNA-anchored H57-scF_V_s free to move without restrictions (DNA-H57, δ~5nm), M-H57 (δ=48 nm) and L-H57 (δ=60 nm) are shown. Scale bar, 200 nm. **f,** Nearest neighbor distances (nnds) of ligand positions identified in (**e)**. Minimum ligand distances δ are indicated as grey lines. 6.8% and 13.5% of determined nnds were below δ for M and L platforms, respectively. Data are from at least 17 cells recorded in three independent experiments.

When confronting CD4^+^ effector T-cells with this novel biointerface we found that T-cell activation via monovalent anti-TCR scF_V_ required close proximity (≤ 20 nm) of ligands within units of at least two molecules, which is in line with previous studies using stably binding ligands^15,16^. Intriguingly, pMHCs did not exhibit this requirement but instead stimulated T-cells robustly as well-isolated entities. Together, our results indicate that early T-cell signaling emerges from small assemblies of triggered TCRs, which can be formed either by stable TCR-binding of closely spaced anti-TCR scF_V_ molecules or by repeated short-lived engagement of single pMHC molecules.

## RESULTS

### Mobile DNA origami platforms for nanoscale ligand organization

We designed rectangular DNA origami tiles of different sizes and ligand occupancies (**Fig. 1b, Supplementary Figs. 1-5**). At sites chosen for modification, staple strands were elongated and hybridized with biotin-conjugated oligonucleotides. To rule out double ligand occupancy at a single modification site, we used divalent streptavidin (dSA)^19,20^ for the attachment of site-specifically biotinylated TCR ligands (**Fig. 1c**). dSA distances on DNA origami platforms featuring two ligand attachment sites were verified via DNA Points Accumulation for Imaging in Nanoscale Topography (DNA-PAINT)^21^, a localization-based super-resolution microscopy method using freely diffusing dye-labeled oligonucleotides that transiently bind to their target-bound complements to achieve the necessary target “blinking” (**Fig. 1d**; for details we refer to **Supplementary Fig. 5**). Two types of ligands (labeled with AlexaFluor™ 555, AF555) were used: (i) a single-chain antibody fragment derived from the TCRβ-reactive monoclonal antibody H57-597 (H57-scF_V_)^22^ and (ii) the nominal activating ligand of the 5c.c7 TCR, i.e. the IE^k^-embedded moth cytochrome c peptide (pMHC).

Our goal was to create a nanostructured biointerface that allows for independent control of overall ligand surface density as well as nanoscale spacing while permitting the large-scale reorganization of ligated receptors to microclusters in the course of immunological synapse formation and T-cell activation. For this, ligand-decorated DNA origami were anchored via cholesterol-modified oligonucleotides^23^ to SLBs containing His-tagged adhesion (ICAM-1) and costimulatory (B7-1) molecules at a density of 100 molecules per μm^2^ each. As shown in **Fig. 1b**, we designed DNA origami platforms of three different sizes: 30 x 20 nm (<u>S</u>mall), 65 x 54 nm (<u>M</u>edium) and 100 x 70 nm (<u>L</u>arge). SLBs featuring ligands attached via either a His-tagged monovalent streptavidin (mSA) or a DNA/dSA anchor were used here as control, as they do not impose any external spatial constraints on ligand organization. The diffusion coefficients of all SLB-anchored DNA origami platforms were about half of that of mSA-anchored H57-scF_V_s, with *D_origami-scFv_* ~ 0.25 μm^2^ s^−1^ and *D_mSA-scFv_* = 0.6 μm^2^s^−1^ (**Supplementary Fig. 6, Supplementary Table 1**). Ligand occupancies of DNA origami platforms were determined via single molecule microscopy of the biointerfaces and by comparing the platform brightness to the brightness of a single fluorescent ligand molecule (**Supplementary Fig. 7**). For DNA origami platforms with a single modification site, functionalization efficiency was ~60%; platforms with two sites for modification yielded a mixed population featuring two ligands (~40%), one ligand (~45%) and no ligand (~15%) (**Supplementary Table 2**). Note that our functionalization strategy was optimized to strictly avoid the presence of two ligands at a single modification site since double occupancies at less than 1% of modification sites might bias our results. For each experiment, the surface density of ligands on SLBs was assessed by relating the average fluorescence signal per area to the brightness of a single fluorescent ligand molecule.

To verify that DNA origami platforms were effective in physically separating ligands, we used DNA-PAINT super-resolution imaging to determine nearest neighbor distances (nnds) of ligands within T-cell-induced microclusters (**Fig. 1e, Supplementary Figs. 8,9**). For this, T-cell blasts isolated from CD4^+^ 5c.c7 TCR transgenic mice were incubated for 10 minutes on biointerfaces featuring H57-scF_V_-functionalized DNA origami platforms at ligand densities of ~4 μm^−2^ and fixed afterwards. We analyzed medium (M-H57) and large (L-H57) DNA origami platforms, as well as H57-scF_V_s directly anchored to the bilayer, which lacked DNA origami-induced exclusion zones (DNA-H57). T-cells on all biointerfaces showed characteristic microcluster formation. For M-H57 and L-H57, nearest neighbor distances between individual ligands measured within microclusters corresponded well with minimum ligand distances *δ* as permitted by the DNA origami platforms (48 nm and 60 nm, respectively; see **Figure 1f**). Note that for DNA-H57, ligand positions could not be assigned as ligands were spaced below the resolution limit of DNA-PAINT at the experimental conditions applied (~15 nm); here, qPAINT analysis^25^ yielded an estimated average ligand spacing of ~8 nm.

### Physical separation of ligated TCRs in microclusters compromises T-cell activation

Upon TCR-engagement with stimulatory ligands, TCR-proximal signaling results in increased levels of intracellular calcium acting as second messenger to promote T-cell activation. We thus conducted live-cell ratiometric calcium imaging using the calcium-sensitive dye Fura-2 AM to monitor activation levels of T-cells (for details on the analysis of calcium traces please refer to the Methods section and **Supplementary Figs. 10-12**). The percentage of activated cells was determined for each biointerface at different ligand densities. Data were plotted as dose-response curves (**Fig. 2a**), and fitted with Eq. 6 (see Methods section) to determine the ligand densities at half-maximum response, hereafter referred to as “activation threshold”. All fit parameters are listed in **Supplementary Table 3**.

**Figure 2.**
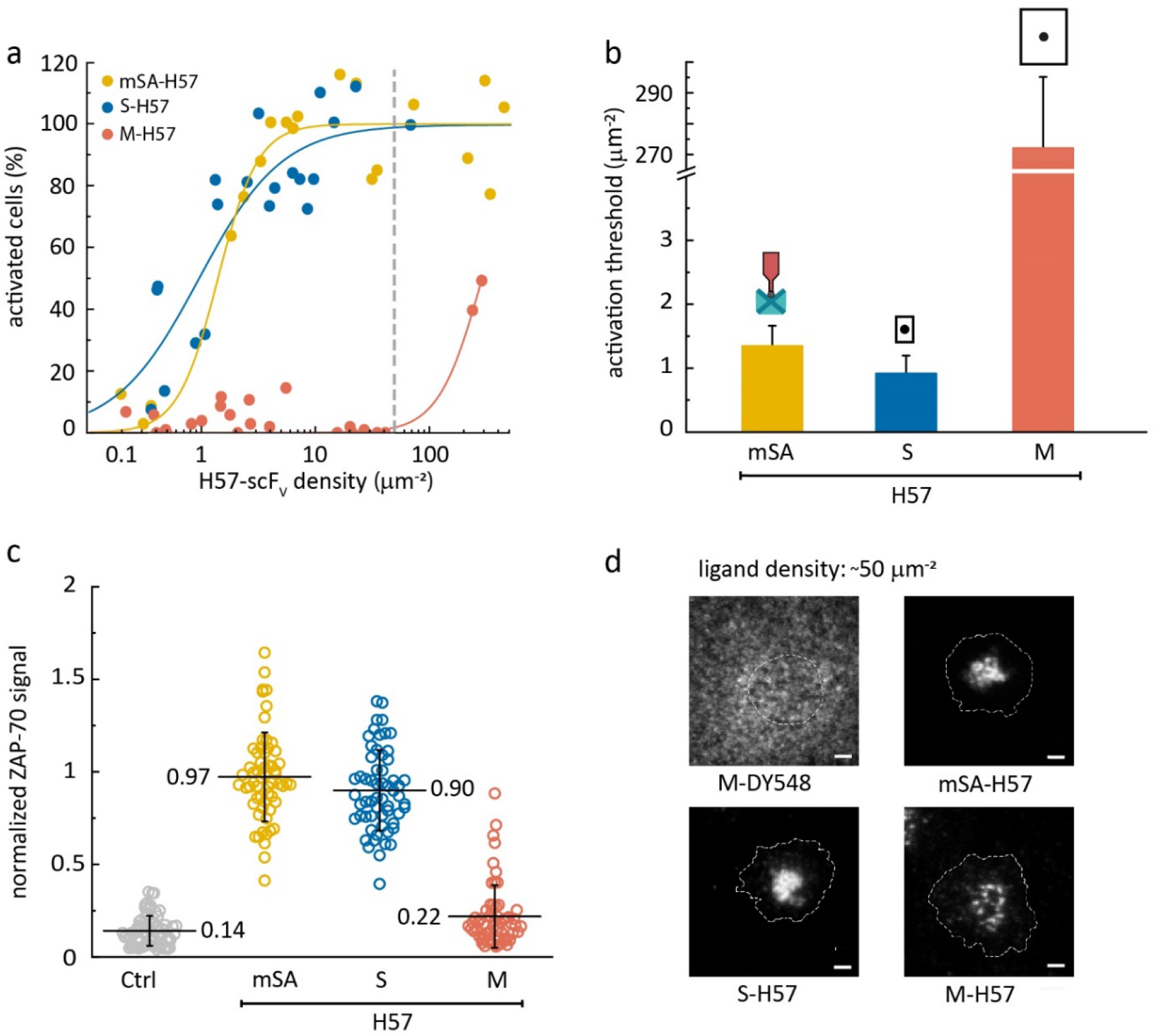
H57-scF_V_-induced T-cell activation depends on ligand spacing. **a,** Percentage of activated T-cells at different ligand surface densities. Data were normalized to a positive control (=100%) containing His-tagged ICAM-1 and B7-1 at 100 μm^−2^, and His-tagged pMHC (His-pMHC) at 150 μm^−2^. Each data point represents the population average of 241 ± 46 cells (mean ± s.e.m.) at a given ligand density. Dose-response curves were fitted with Eq. 6 (see Methods section) to extract activation thresholds **(b).** For each construct, data are from at least three independent experiments and three different mice. Error bars represent the standard error of the fits. **c**, ZAP-70-recruitment to the T-cell plasma membrane at ligand densities of ~50 μm^−2^, indicated by a vertical dashed grey line in **(a)**. T-cells were fixed 10 minutes after cell seeding, and immunostained for ZAP-70. The signal was analyzed and normalized using a positive control (as in **(a)**) as a reference. Cells on bilayers containing only ICAM-1 and B7-1 at 100 μm^−2^ are shown as a negative control. Data are shown as means ± s.d., each data point representing a single cell. n > 50 cells from at least three independent experiments and three different mice. **d,** Representative TIRF images for the different constructs featuring AF555-labeled ligands at densities of ~50 μm^−2^. The cell outline is indicated by a dashed white contour line. Images were recorded 10 minutes after seeding. Dy548-labeled DNA origami platforms without ligands at ~50 μm^−2^ are shown for comparison. In order to visualize signals for M-DY548, image brightness was increased by a factor of 10. Scale bar, 2 μm.

H57-scF_V_ anchored to SLBs via either mSA (**Fig. 2a, b**) or DNA/dSA (**Supplementary Fig. 13**) activated T-cells at thresholds amounting to ~ 1 molecule per μm^2^, similar to nominal SLB-resident pMHCs (see **Fig. 4**). Attachment of H57-scF_V_ to small DNA origami platforms (S-H57), which allow for close packing of ligands down to distances *δ* of 20 nm (**Supplementary Table 4**), did not affect recorded activation thresholds. However, for M-H57, which prevent ligand packing below *δ* = 48 nm, the activation threshold increased by more than 2 orders of magnitude. Even at surface densities of ~200 μm^−2^, at which the SLB was almost fully covered by M-H57, T-cell activation was inefficient. Of note, the presence of high densities of ligand-free DNA origami platforms on SLBs (~50 μm^−2^) neither activated T-cells on bilayers without ligand, nor did they impede T-cell activation driven by bilayer-anchored His-pMHC (**Supplementary Fig. 14**), implying that platform-enforced ligand distancing rather than the mere presence of DNA origami platforms interfered with T-cell activation. Activation trends observed at 24°C were maintained at 37°C yet with lower activation thresholds (**Supplementary Fig 15a, b**).

Calcium signaling is initiated downstream of a tyrosine kinase cascade involving the TCR and TCR-proximal phosphorylation targets. The cytoplasmic tyrosine kinase ZAP-70 binds to phosphorylated immunoreceptor tyrosine-based activation motifs (ITAMs) present on the cytoplasmic tails of the TCR-associated CD3 subunits^26^. To determine whether early T-cell signaling events such as ZAP-70 recruitment are subject to minimal ligand distance requirements, we performed immunostaining experiments on T-cells, which had been fixed 10 minutes after their encounter with the biointerface at a ligand density of ~ 50 μm^−2^. As shown in **Fig. 2c**, ZAP-70 recruitment from the cytoplasm was observed for mSA-H57 and S-H57 but not M-H57, in agreement with the calcium response recorded for H57-scF_V_ attached to the different constructs. This implies that distancing of H57-scF_V_s affects TCR-signaling in its earliest stages. Of note, TCR microclusters and the characteristic central supramolecular activation cluster (cSMAC) were induced by all constructs at activating ligand densities of ~50 μm^−2^, yet also for M-H57, which failed to promote ZAP-70 recruitment and calcium signaling (**Fig. 2d, Supplementary Fig. 16**).

### Pairs of closely spaced ligated TCRs are sufficient units for T-cell activation

To assess the geometric requirements for efficient T-cell activation, we attached two H57-scF_V_s to 65 x 54 nm DNA origami to generate mobile divalent platforms (M_div_-H57) with defined distances between individual H57-scF_V_s of 10 nm, 20 nm and 30 nm (**Fig. 1b, d**). While monovalent platforms of this size no longer supported efficient stimulation (**Fig. 2a, b**), divalent platforms with *δ* = 10 nm and *δ* = 20 nm ligand spacing were highly potent activators (**Fig. 3a, b**). Activation thresholds for M_div_-H57 with *δ* = 30 nm were one order of magnitude higher, but still significantly below those measured for monovalent M-H57.

**Figure 3.**
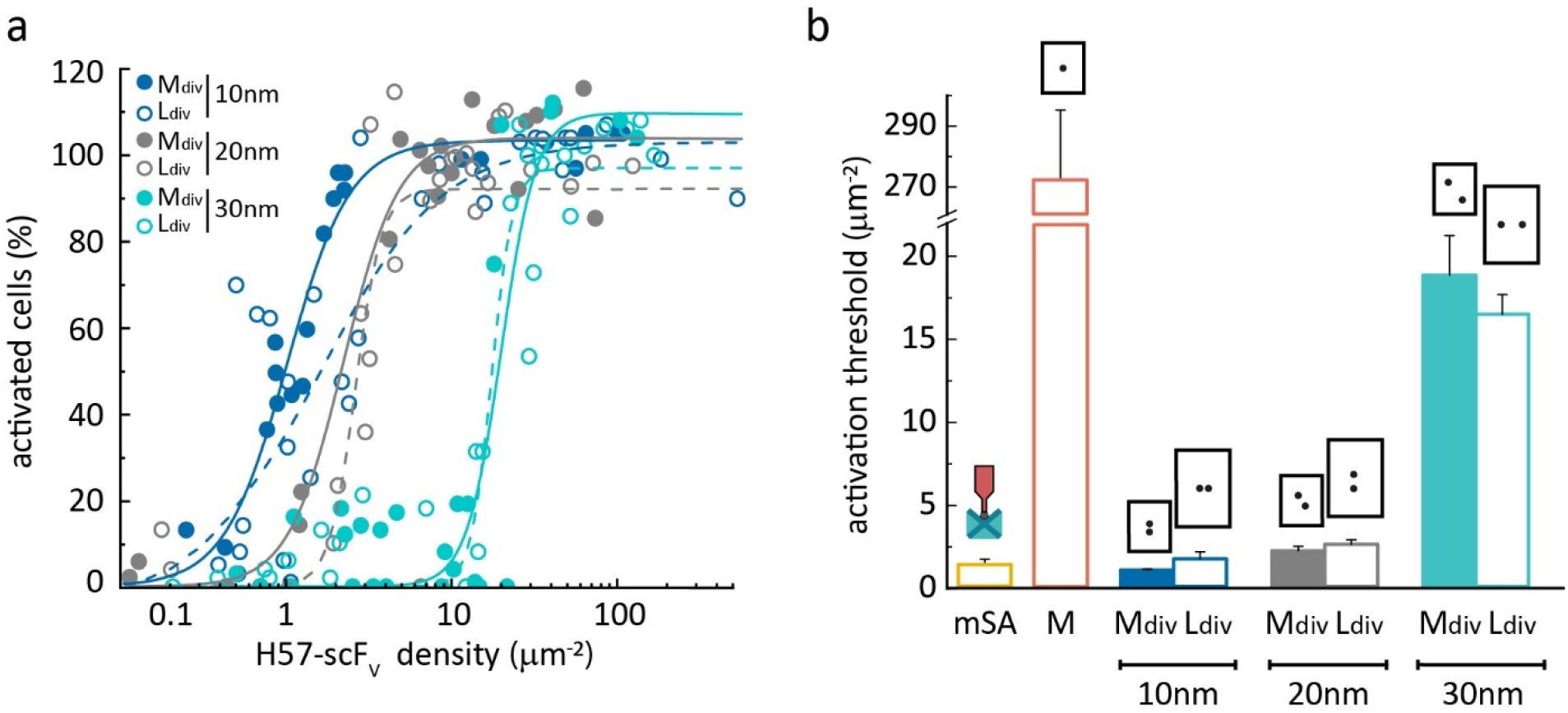
Isolated pairs of closely spaced H57-scF_V_s are sufficient units for T-cell activation. **a,** Dose-repose curves for divalent DNA origami platforms featuring two H57-scF_V_s at different distances. Each data point represents the population average of 264 ± 32 cells (mean ± s. e. m.) at a given ligand density. Fits for L_div_-H57 using Eq. 6 are shown as dashed lines. For each construct, data are from at least three independent experiments and three different mice. **b,** Activation thresholds determined from fits of data from **a.** Data for mSA-H57 are shown as reference. Error bars represent the standard error of the fits.

Although unlikely, we cannot completely rule out that two neighboring M_div_-H57 platforms contributed to signaling. In fact, the smallest possible distance between two ligands on two different adjacent M_div_-H57 platforms is ~40 nm for all layouts (**Supplementary Table 4**), which is slightly below the minimum ligand separation on monovalent M DNA origami platforms. To increase the minimal distance between two ligand pairs placed on two distinct DNA origami platforms, we increased the platform size to 100 x 70 nm (L_div_-H57). As shown in **Fig. 3**, activation thresholds were not markedly affected by this: specifically, for L_div_-H57 with 10 and 20 nm ligand spacing, for which the minimal inter-platform distance between individual ligands amounted to ~60 nm, we determined similar activation thresholds when compared to those measured for mSA-H57. These observations imply that isolated H57-scFv ligand pairs are by themselves sufficiently stimulatory to activate T-cells.

Due to incomplete functionalization, M_div_ and L_div_ DNA origami platforms contained fractions of platforms carrying either two, one or no ligands. Adding an even 10-fold molar excess of monovalent M-H57 to M_div_-H57_10 nm_, however, did not affect the activation threshold (**Supplementary Fig. 17**). Hence, when using a mixture of DNA origami platforms featuring two, one or no H57scF_V_, only those decorated with two scF_V_s contributed to the signaling response. We observed efficient ZAP-70 recruitment (**Supplementary Fig. 18**) and cSMAC formation (**Supplementary Fig. 11d-i**) for all divalent platforms at ligand densities of ~50 μm^−2^.

### Well-isolated monomeric agonist pMHCs efficiently activate T-cells

Results obtained using stably binding TCR-reactive antibodies provide information concerning spatial TCR-arrangements and - stoichiometries required for productive signaling. It may be unjustified, however, to conclude on similar spatial constraints underlying T-cell stimulation via the physiological TCR-ligand pMHC. After all, pMHCs bind TCRs transiently with lifetimes of seconds rather than tens of minutes in a docking angle that is fairly conserved among stimulatory TCR:pMHC pairs. Furthermore, pMHCs offer a second epitope for coreceptor engagement, which is known to sensitize T-cells for antigen by a factor of 10 to 50 ^2^. We hence assessed whether T-cell stimulation via nominal pMHCs followed similar spatial distancing requirements as H57-scF_V_-mediated T-cell activation.

Intriguingly, activation thresholds for nominal pMHCs did not depend on the manner of ligand presentation: pMHCs tethered to mSA (mSA-pMHC) and medium sized DNA origami platforms (M-pMHC) gave rise to similar dose-response curves at 24°C (**Fig. 4a, b)** and at 37°C **(Supplementary Fig. 15c,d)**. Even when placing pMHCs on large DNA origami platforms with *δ* = 60 nm (L-pMHC) the efficiency of T-cell triggering was unperturbed.

**Figure 4.**
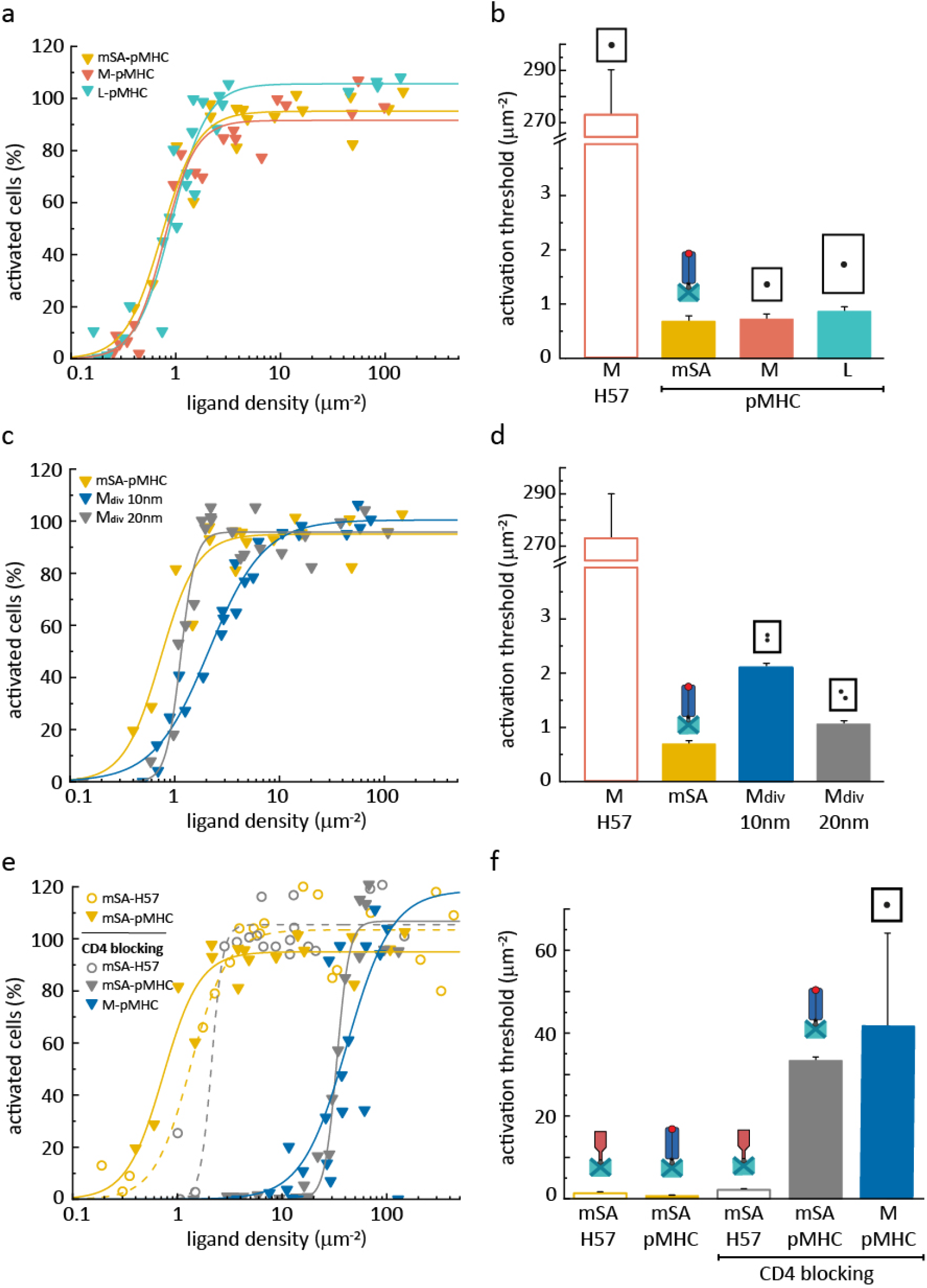
T-cell activation is independent of pMHC spacing. Dose-response curves of monovalent **(a)** and divalent **(c)** DNA origami platforms featuring pMHC measured at 24°C, and after blocking of CD4 with anti-CD4 Fab **(e)**. Each data point represents the population average of 225 ± 56 **(a)**, 189 ± 85 **(c)** and 223 ± 53 **(e)** cells (mean ± s.e.m.) at a given ligand density. For each construct, data are from at least three independent experiments and three different mice. Data for mSA-H57 and M-H57 are shown as references. **b, d, f**, Activation thresholds determined from fits of data from **a, c** and **e**. Error bars represent the standard error of the fits.

Furthermore, pre-organization of two pMHC molecules on divalent DNA origami platforms did not enhance their stimulatory potency. On the contrary, activation thresholds were slightly increased at a spacing of 10 nm, possibly due to steric hindrance between adjacent pMHCs (**Fig. 4c, d**). This effect may even be underrepresented in our study since T-cell activation triggered by DNA origami platforms featuring one instead of two pMHCs could not be accounted for. All pMHC constructs produced high levels of ZAP-70 recruitment (**Supplementary Fig. 18b**) and showed clustering on the SLB at ligand densities of ~50 μm^−2^ (**Supplementary Fig. 19**).

To test whether CD4-binding to pMHC affected the different spatial requirements for TCR triggering observed for pMHC and H57-scF_V_, we blocked CD4 with an anti-CD4 Fab prior to cell seeding. While this markedly increased activation thresholds, the observed effect was similar for pMHC anchored directly to the SLB or attached to DNA origami. In control experiments, T-cell activation thresholds for SLB-anchored H57-scF_V_ remained unchanged by CD4 blocking (**Fig. 4e, f**).

## DISCUSSION

The molecular organization of membrane-bound ligands and receptors at cell-cell interfaces is critical for cellular communication ^27–30^ yet presents inherently a formidable challenge to investigators. On the one hand, experimental approaches involving molecularly defined biochemical or structural biology techniques invariably remove molecular players from the live cell context. On the other hand, even state-of-the art imaging reaches its limits given the spatial and temporal resolution necessary to follow nanoscale reorganization processes *in situ*. Antigen recognition by T-cells illustrates this conundrum: while central to adaptive immunity and with most molecular players already identified, knowledge on its operational principles is still limited. Key to identifying molecular mechanisms are cellular interventions that allow for manipulating the organization of molecules in a live cell environment.

With this in mind, we have devised a DNA origami-based biointerface, which allows for experimentally controllable and defined adjustment of protein distances while supporting at the same time the cell-driven spatial reorganization of ligand and receptor molecules in the course of a cellular signaling process. In this study we applied our biointerface to identify ligand arrangements for productive TCR triggering, both by imposing steric constraints to cell-mediated clustering processes, and by pre-arranging clusters of ligands. While efficient T-cell activation via monovalent TCRβ-reactive scF_V_s required close proximity of ligands within units of at least two molecules, such a requirement was absent for the natural ligand, i.e. nominal pMHC (**Fig. 5a**), which stimulated T-cells effectively when present as individual, well-separated entities. This disparity is likely rooted in the fundamentally different nature by which scF_V_s and pMHCs interact with the TCR.

**Figure 5.**
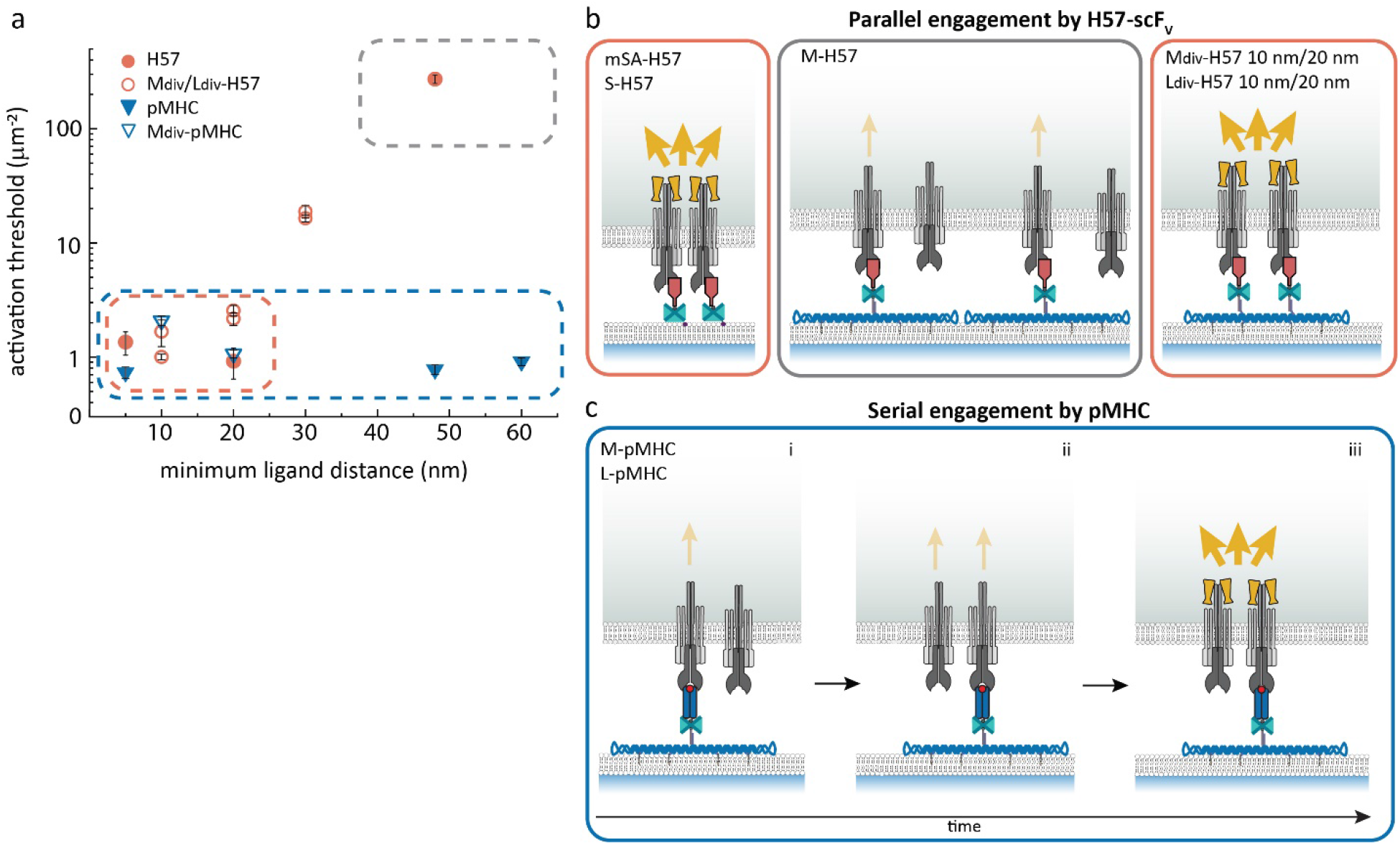
Ligand-specific spatial requirements for T-cell activation. **a,** Minimum ligand distances δ are plotted versus the activation threshold. For DNA origami platforms featuring a single ligand, δ corresponds to the smallest possible distances between two ligands on adjacent platforms assuming quasi-crystalline packing in clusters. For divalent platforms, it corresponds to the predefined distance between the two ligands on an individual platform. The lateral extension of mSA-anchored ligands was approximated with 5 nm. **b,** Parallel engagement model of antibody-induced triggering. Two or more triggered TCRs within 20 nm form signaling-competent TCR assemblies resulting in ZAP-70 recruitment (yellow dumbbells) and initiation of calcium signaling (yellow arrows). H57-scF_V_ – triggered T-cell activation requires at least two simultaneous ligand:receptor engagements within a distance of 20 nm (“parallel engagement”). This can occur via T-cell induced clustering of mSA-anchored scF_V_s or pre-organization of scF_V_s on divalent DNA origami platforms. H57-scFvs isolated on M DNA origami platforms at δ=48 nm fail to stimulate T-cells efficiently. **c,** Serial engagement model of pMHC-induced triggering. A single isolated pMHC molecule can create a signaling-competent TCR assembly by sequentially engaging multiple TCRs. **i,** A single pMHC:TCR binding event does not induce efficient downstream signaling. **ii**, Via serial, short-lived binding events, a single isolated pMHC molecule may engage several TCRs sequentially. **iii,** If two or more TCRs within a distance of 20 nm are triggered in this way, a signaling-competent TCR assembly is created.

With an interaction half-life of ~50 minutes at room temperature^22^, a H57-scF_V_ molecule can be expected to engage a single TCR and stay bound for the entire duration of the experiment. As a consequence, the specific arrangement of H57-scF_V_ serves as a template for the molecular organization of ligated TCR molecules. In this case, T-cell stimulation was only observed under conditions that allowed for close proximity of ligated TCRs in clusters (*δ* ≤ 20 nm). In agreement with Wind and colleagues^15^, enforcement of minimum ligand distances of 48 nm created a signaling-impaired state. Furthermore, isolated TCR pairs enforced by divalent DNA origami platforms were still signaling-competent: platforms featuring scF_V_s at 10 and 20 nm distance and thus closely mimicking IgG antibodies^31^ were most potent. Given the lateral dimensions of the TCR-CD3 complex with a diameter of about 10 nm^32^, we consider it likely that close apposition of two TCR-CD3 complexes but not necessarily their physical association is required for efficient initiation of intracellular signaling. In fact, a recent FRET-based study of ours failed to provide any evidence for the formation of physically linked dimeric TCR-CD3 structures upon engagement with nominal pMHCs^6^.

Let us examine the conditions necessary to create signaling-competent TCR assemblies in more detail: as concluded from our experiments on H57-scF_V_s, configurations supporting the parallel engagement of two proximal TCRs are sufficient for T-cell activation (**Fig. 5b**). In general, in case of freely diffusing ligands the probability for the formation of such parallel engagements depends on the ligand (and receptor) density as well as the ligand:receptor dwell time. Evidently, in our system this condition was met at a minimum density of approximately 1 H57-scF_V_ molecule per μm^−2^ (**Fig. 2b**), which amounts to 100 or more scF_V_ molecules per T-cell synapse. Note that the high-affinity of the H57-scF_V_:TCR interaction impeded the dissociation of any ligand:receptor pairs, once they had formed. Similar requirements for productive signaling were previously reported in a chimeric antigen receptor (CAR) system, where DNA hybridization was used to mimic receptor-ligand interactions^16^: in that system, initiation of signaling was found to depend on clustering of ligated receptors, which, in turn, required high ligand densities (≥ 1 μm^−2^) and long ligand:receptor dwell times.

In a natural APC – T-cell interface, however, neither of these conditions applies: the nominal antigen density on APCs can be as low as one to five antigenic pMHC molecules per cell^33,34^ and dwell times for the TCR:pMHC interaction are short (1.7 s and 100 ms at room temperature and 37°C, respectively)^22^. The probability of stochastic occurrence of parallel engagements is thus very low, suggesting that pMHC-triggered T-cell activation does not hinge on the same mechanism. In fact, signaling was unperturbed when we actively prohibited the formation of parallel engagements by using DNA origami platforms to isolate individual pMHC molecules. While it is possible that individual, well-isolated pMHC-engaged TCRs effectively mediate downstream signaling, the importance of TCR clustering for productive signaling^35^ argues against this. It appears more plausible that pMHC-mediated TCR triggering gives rise to similar signaling-competent TCR assemblies as H57-scF_V_ does, but without the need for either ligand pre-organization or T-cell-induced ligand clustering. In such a model, the short dwell times allow for sequential binding of the same pMHC molecule to multiple TCRs; as a consequence, a single isolated pMHC molecule may create an assembly of signaling-competent TCRs by serial engagement of two or more TCRs within a range of 20 nm (**Fig. 5c**). Indeed, sequential engagement and triggering of up to ~200 TCRs by a single pMHC molecule has been reported^36^ and proposed to enhance the T-cell’s sensitivity for antigen, particularly at low antigen densities^37^.

In conclusion, we have demonstrated that well-isolated monomeric agonist pMHC molecules efficiently stimulate T-cells. Based on our results we propose that signaling-competent TCR assemblies can be created either via parallel engagements by closely spaced high-affinity ligands or via serial, rapid and short-lived engagements by a single nominal pMHC. This notion integrates well with recent findings on TCR triggering by pMHC^6,17^ as well as artificial ligands^15,16^ and reconciles the frequently observed relevance of TCR clustering^35^ with the single molecule sensitivity of sensitized antigen detection^18^.

## MATERIALS AND METHODS

### Assembly of DNA origami platforms

DNA origami structures were assembled in a single folding reaction carried out in a test tube (AB0620, ThermoFisher Scientific) with 10 μl of folding mixture containing 10 nM M13mp18 scaffold DNA (New England Biolabs), 100 nM unmodified oligonucleotides (Integrated DNA technologies, Eurofins), 500 nM fluorescently labeled (IBA Lifesciences) or biotinylated oligonucleotides (Biomers) and folding buffer (5 mM Tris (AM9855G, ThermoFisher Scientific), 50 mM NaCl (AM9759, ThermoFisher Scientific), 1 mM EDTA (AM9260G, ThermoFisher Scientific), 12.5 mM MgCl_2_) (AM9530G, ThermoFisher Scientific)). Oligonucleotide sequences are shown in **Supplementary Tables 5-8**. At sites chosen for ligand and cholesterol anchor attachment, staple strands were elongated at their 3’-end with 21 and 25 bases, respectively. DNA origami were annealed using a thermal protocol (90°C, 15 min; 90°C – 4°C, 1°C/min; 4°C, 6h) and purified using 100kDa Amicon^®^Ultra centrifugal filters (UFC510096, Merck). DNA origami were stored up to 4 weeks at −20°C. For functionalization, DNA origami were incubated with a 10x molar excess of dSA for 30 min at 24°C and excessive dSA was removed using 100kDa Amicon^®^Ultra centrifugal filters (UFC210024, Merck). As a last step, AF555-conjugated and site-specifically biotinylated TCR ligands (H57-scF_V_, pMHC) were added at a 10x molar excess for 60 min at 24°C. Functionalized DNA origami platforms were used for experiments at the same day.

### Agarose gel electrophoresis

DNA origami were mixed with DNA loading buffer (B7025S, New England Biolabs) and subjected to agarose gel electrophoresis (1% agarose (A9539, Sigma Aldrich), 1x Tris Acetate-EDTA (TAE) (15558042, ThermoFisher Scientific), 10 mM MgCl_2_) to validate correct folding. Agarose gels were run at 24°C for 75 min at 100V, stained with 1xSYBR™ gold nucleic acid stain (S11494, ThermoFisher Scientific)) and visualized with a Gel Doc™ XR+ (Bio Rad).

### High speed AFM imaging

High speed AFM (HS-AFM)^38–41^ (RIBM, Japan) was conducted in tapping mode at 24°C in AFM imaging buffer (40 mM Tris, 2 mM EDTA, 12.5 mM MgCl_2_), with free amplitudes of 1.5 - 2.5 nm and amplitude set points larger than 90%. Silicon nitride cantilevers with electron-beam deposited tips (USC-F1.2-k0.15, Nanoworld AG), nominal spring constants of 0.15 Nm^−1^, resonance frequencies around 500 kHz, and a quality factor of approximately 2 in liquids were used. For sample preparation, 2 μl of 500 μM MgCl_2_ solution was preincubated on a freshly cleaved mica surface for 5 min followed by a washing step with deionized water. 2 μl of purified DNA origami solution (1:10 diluted in AFM imaging buffer) were applied to the mica surface for 5 min. Finally, the sample was washed with AFM imaging buffer.

### Preparation of functionalized planar SLBs

Vesicles containing 98% 1-palmitoyl-2-oleoyl-sn-glycero-3-phosphocholine (POPC) and 2% 1,2-dioleoyl-sn-glycero-3-[N(5-amino-1-carboxypentyl)iminodiaceticacid]succinyl[nickel salt] (Ni-DOGS NTA) (Avanti Polar Lipids) were prepared at a total lipid concentration of 0.5mg ml^−1^ as described ^22^ in 10x Dulbecco’s phosphate-buffered saline (PBS) (D1408-500ml, Sigma Aldrich). Glass coverslips (#1.5, 24×60 mm, Menzel) were plasma cleaned for 10 min and attached with the use of dental imprint silicon putty (Picodent twinsil 22, Picodent) to Lab-Tek™ 8-well chambers (ThermoFisher Scientific), from which the glass bottom had been removed^42^. Coverslips were incubated with a fivefold diluted vesicle solution for 10 min, before they were extensively rinsed with PBS (D1408-500ML, Sigma Aldrich). For functionalization, SLBs were first incubated for 60 min with cholesterol-oligonucleotides (Integrated DNA technologies) complementary to the elongated staple strands at the bottom side of the DNA origami and then rinsed with PBS. DNA origami were incubated on SLBs in PBS + 1% BSA (A9418-10G, Sigma Aldrich) for 60 min. Finally, His10-tag ICAM-1 (50440-M08H, Sino Biological) (270 ng mL^−1^) and His10-tag B7-1 (50446-M08H, Sino Biological) (130 ng mL^−1^) were incubated for 75 min at 24°C and then rinsed off with PBS. PBS was replaced with HBSS for imaging (H8264-500ML, Sigma Aldrich).

### Total internal reflection fluorescence (TIRF) microscopy

TIRF microscopy experiments were performed on a home-built system based on a Zeiss Axiovert 200 microscope equipped with a 100x, NA=1.46 Plan-Apochromat objective (Zeiss). TIR illumination was achieved by shifting the excitation beam parallel to the optical axis with a mirror mounted on a motorized table. The setup was equipped with a 488 nm optically pumped semiconductor laser (Sapphire, Coherent), a 532 nm diode-pumped solid state (DPSS) laser (Spectra physics Millennia 6s) and a 640 nm diode laser (iBeam smart 640, Toptica). Laser lines were overlaid with an OBIS Galaxy beam combiner (Coherent). Acousto-optic modulators (Isomet) were used to modulate intensity (1-3kW/cm^2^) and timings using an in-house developed package implemented in LABVIEW (National Instruments). A dichroic mirror (Di01-R405/488/532/635-25×36, Semrock) was used to separate excitation and emission light. Emitted signals were split into two color channels using an Optosplit II image splitter (Oxford Instruments) with a dichroic mirror (DD640-FDi01-25×36, Semrock) and emission filters for each color channel (ET 570/60, ET 675/50, Chroma) and imaged on the same back-illuminated Andor IXON Ultra EMCCD camera.

### DNA origami platform characterization and determination of ligand surface densities

Ligand occupancies of SLB-anchored DNA origami were determined via brightness analysis. In a first step, the dark fraction of DNA origami, bearing no ligand, was determined for each construct separately via two-color colocalization experiments. For this, DNA origami were modified with a single or two biotinylated oligonucleotides labeled with Abberior STAR 635P (Biotin-DNA-AS635P) and pre-stained with YOYO™-1 iodide (YOYO) at a concentration of 1μg ml^−1^ for 45 min at 24°C. Excessive YOYO was removed using 100 kDa Amicon^®^ Ultra centrifugal filters and DNA origami-bearing SLBs were produced as described above. Positions of diffraction-limited spots were determined for both color channels and corrected for chromatic aberrations as described^6^. Detected signal positions were counted as colocalized if signals were within a distance of 240 nm. The fraction of DNA origami carrying at least one biotin modification, *f*_*coloc*_1_, was determined by relating the number of signals in the red color channel (Biotin-DNA-AS635P) that colocalized with a signal in the green color channel (YOYO), N_coloc_1_, to the number of detected green signals, N_total_1_ (Eq. 1). In a second step, the efficiency of functionalization of existing biotin groups with a ligand, *f*_*coloc*_2_, was determined using DNA origami labeled with Biotin-DNA-AS635P and TCR ligands (H57-scF_V_, pMHC (both labeled with AF555)). For this, the number of green signals (H57-scF_V_, pMHC) that colocalized with a red signal (Biotin-DNA-AS635P), N_coloc_2_, was divided by the number of red signals (Biotin-DNA-AS635P, N_total_2_).

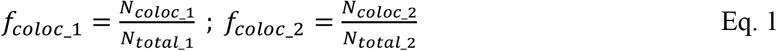

The fraction of DNA origami carrying at least one ligand (*f_bright_*) is then given by

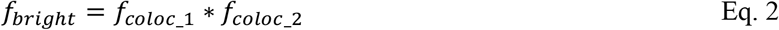

For divalent DNA origami modified at two sites, ligand occupancy was evaluated by using a MATLAB (Mathworks)-based maximum-likelihood estimator to determine position, integrated brightness *B*, full width at half-maximum (FWHM) and local background of individual signals in the images^43,44^. Briefly, DNA origami were anchored to SLBs, and the integrated brightness *B* was determined for all recorded positions. Images were taken at multiple different locations. The brightness values *B* of a monomer reference (bilayer-anchored 2xHis_6_-tag pMHC-555^6^) were used to calculate the probability density function (pdf) of monomers, *ρ*_1_(*B*). Because of the independent photon emission process, the corresponding pdfs of *N* colocalized emitters can be calculated by a series of convolution integrals.

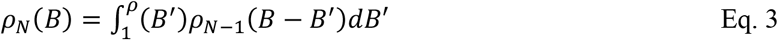

A weighted linear combination of these pdfs was used to calculate the brightness distribution of a mixed population of monomers and oligomers.

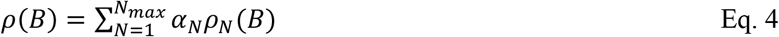

Brightness values for each DNA origami construct were pooled and used to calculate *ρ*_1_(*B*). For H57-scF_V_, brightness values were corrected according to the protein to dye ratio (0.93 – 1.0) of the used H57-scF_V_ preparation (see section ‘Protein expression and purification’). A least-square fit with Eq. 4 was employed to determine the weights of the individual pdfs, *α_N_*, with 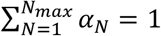. For all fits, no higher contributions than dimers (*α*_2_) were observed. A minimum of ~ 900 brightness values was applied to calculate *ρ*_1_(*B*) and *ρ*_2_(*B*). To account for DNA origami carrying no ligand, *α_N_corrected_*, was determined by multiplying evaluated monomer *α*_1_ and dimer *α*_2_ contributions with the fraction of DNA origami carrying at least one ligand.

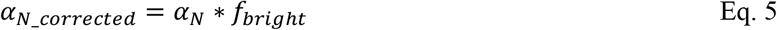

For mobility analysis of DNA origami, ~ 10 image sequences were recorded at different locations on the SLB at an illumination time of 3 ms and a time lag of 10 ms. Images were analyzed using in-house algorithms implemented in MATLAB^45^. Mean-square displacements (MSDs) were averaged over all trajectories, plotted as a function of time lag and the diffusion coefficient (D) was determined by fitting the function *MSD* = 4*Dt* + 4*σ_xy_*, where *σ_xy_* denotes the localization precision; diffusion coefficients were determined from the first two data points of the MSD-t-plot.

Average surface densities of AF555-labeled ligand on SLBs were determined by dividing mean intensities per μm^2^ by the single-molecule brightness of bilayer-anchored AF555-labeled His-pMHC. All experiments were carried out at 24°C, unless stated otherwise.

For experiments with SLBs featuring both M_div_-H57_10 nm_ and M-H57, platforms were functionalized separately as described above. M-H57 platforms were additionally modified with biotin-AS635P to be able to distinguish the two DNA origami platforms on the SLB. To obtain the three different molar ratios of M_div_-H57_10 nm_ : M-H57 (1:0, 1:2, 1:10), appropriate amounts of M-H57 were added to a SLB containing M_div_-H57_10 nm_ after 15 min and both constructs were then incubated for 60 min and then rinsed with PBS. For imaging PBS was replaced for HBSS.

### Sample preparation for DNA-PAINT imaging of divalent DNA origami

On divalent DNA origami platforms (M_div_, L_div_), biotinylated TCR ligands were replaced with biotinylated DNA-PAINT docking strands (Biotin-TEG-V-2T-P3*’) to image ligand positions. Additionally, four staple strands at the corners were extended with P1’ DNA-PAINT docking strands for barcoding. Four buffers were used for sample preparation:

- buffer A (10 mM Tris-hydrochloric acid (HCl), 100 mM NaCl, 0.05% Tween 20, pH 8.0)
- buffer B (5 mM Tris-HCl, 10 mM MgCl_2_, 1 mM EDTA, 0.05% Tween 20, pH 8.0)
- buffer C (1x PBS, 500 mM NaCl, pH 7.2)
- imaging buffer

The imaging buffer consisted of buffer C supplemented with 1x PCA (40x PCA: 154 mg PCA, 10 ml water and sodium hydroxide (NaOH) were mixed and adjusted to pH 9.0), 1x PCD (100x PCD: 9.3 mg PCD in 13.3 ml of buffer (100 mM Tris-HCl pH 8.0, 50 mM potassium chloride (KCl), 1mM EDTA, 50% Glycerol), and 1x Trolox (100x Trolox: 100 mg Trolox, 430μl 100% Methanol, 345 μl

1M NaOH in 3.2ml UP-H2O). Stock solutions were stored at −20°C.

For sample preparation, a μ-slide VI 0.5 glass bottom (80607, Ibidi) was used with a channel volume of 40 μl. Solutions were always added on one side of the channel and removed from the opposite side. 100 μl of biotin-labeled BSA dissolved in buffer A at 1 mg ml^−1^ (A8549-10MG, Sigma-Aldrich) was flushed through the channel five times and incubated for 5 min. After washing with 1 ml buffer A, 100 μl of neutravidin (0.5 mg ml^−1^ in buffer A, (31000, ThermoFisher Scientific)) was flushed through the channel five times and incubated for 5 min. After washing the channel with 3 ml buffer A a solution of 100 μl of biotin-modified DNA strands (Z-TEG-Biotin) in buffer A at 1 μM was flushed through five times and incubated for 15 min. The channel was then washed with 2 ml of buffer B before flushing in 5 times 100 μl of the DNA origami in buffer B at a concentration of 500 pM. DNA origami were incubated for 20 min and washed with 1 ml buffer B. 100 μl of 90 nm standard gold nanoparticles (G-90-100, cytodiagnostics) diluted 1:5 in buffer C were added and incubated for 3 min before washing with 2 ml of buffer C. Finally, 500 μl of the imaging solution containing 8 nM of the imager strand P1-Cy3B was flushed in to image barcode positions. After washing the channel with 3 ml buffer C, 500 μl of 10 nM imager strand P3-Cy3B was flushed in to image ligand positions. Imaging parameters for DNA-PAINT on divalent DNA origami platforms are shown in **Supplementary Table 9**.

### Sample preparation for DNA-PAINT imaging of DNA origami platforms within microclusters

For DNA-PAINT imaging, biotinylated oligos were extended with DNA-PAINT docking sites at their 3’ end (P3*’-2T-Z’-4T-TEG-Biotin) and DNA origami platforms were functionalized as described above by attaching biotinylated H57-scF_V_ labeled with AS635P. A DNA-anchored H57-scF_V_ free to move without restrictions (DNA-H57) was assembled by incubating P3*’-2T-Z’-4T-TEG-Biotin with a 10x molar excess of dSA at 24°C for 15 min, followed by addition of H57-scF_V_ to the second available binding pocket of the dSA. TCR ligands were added at a 10x molar excess over dSA at 24°C for 60 min. Finally, the construct was anchored to complementary cholesterol oligonucleotides on SLBs. DNA origami without ligand were labeled by hybridizing an Alexa Fluor™647 (AF647)-labeled oligo to an elongated staple strand next to the ligand attachment site. Prior to T-cell seeding, ligand surface densities on SLBs were determined as described before. 4.0 *10^6^ cells per well were seeded onto SLBs and allowed to settle for 10 min at 24°C. Fixation buffer (PBS, 8% formaldehyde (28908, ThermoFisher Scientific), 0.2% glutaraldehyde (G7776-10ML, Sigma Aldrich), 100 mM sodium orthovandate (Na_3_VO_4_) (S6508-10G, Sigma Aldrich), 1M sodium fluoride (NaF) (S7920-100G, Sigma Aldrich)17) was added 1:1 and incubated for 20 min at 24°C, before samples were rinsed with PBS. Permeabilization buffer (PBS, 100 mM Na_3_VO_4_, 1 M NaF, 0.2% Triton-X (85111, ThermoFisher Scientific) was added and after 1 min samples were washed with PBS before blocking with passivation buffer (PBS, 3% BSA) for 30 min. MgCl_2_ was added additionally to each buffer to a final concentration of 10 mM. Samples were extensively washed with passivation buffer and 0.05% sodium-azide was added to each well. Samples were kept up to 72 h at 4°C prior further usage.

Before adding gold fiducials, samples were washed with 100 μl of buffer C. Subsequently, 100 μl of 90 nm standard gold nanoparticles, diluted 1:2 in buffer C, were added and incubated for 3 min before washing with 2 ml of buffer C. Prior image acquisition, all fluorophores were deactivated by a high intensity bleach pulse. Imaging parameters for DNA-PAINT cell experiments are detailed in **Supplementary Table 10.** 500 μl of imaging buffer containing imager strand P1-Cy3B at 8 nM was flushed in to image barcode positions. After washing the channel with 4 ml buffer C, 500 μl of 10 nM imager strand P3-Cy3B in imaging buffer were added to image ligand positions.

### DNA-PAINT super-resolution microscopy setup

#### Super-resolution microscope 1

DNA-PAINT imaging was partly carried out using an inverted Nikon Eclipse Ti microscope (Nikon Instruments) and the Perfect Focus System, by applying an objective-type total internal reflection fluorescence (TIRF) configuration with an oil-immersion objective (Apo SR TIRF 100x, NA 1.49, oil). Two lasers were used for excitation: 640 nm (150 mW, Toptica iBeam smart) or 561 nm (200 mW, Coherent Sapphire). They were coupled into a single-mode fiber, which was connected to the microscope body via a commercial TIRF Illuminator (Nikon Instruments). The laser beam was passed through cleanup filters (ZET642/20 or ZET561/10, Chroma Technology) and coupled into the microscope objective, using a beam splitter (ZT647rdc or ZT561rdc, Chroma Technology). Fluorescence light was spectrally filtered with two emission filters (ET705/72m or ET600/50m, Chroma Technology) and imaged with a sCMOS camera (Andor Zyla 4.2) without further magnification, resulting in an effective pixel size of 130 nm after 2 x 2 binning. Camera readout sensitivity was set to 16-bit, readout bandwidth to 540 MHz.

#### Super-resolution microscope 2

DNA-PAINT imaging was partly carried out using an inverted Nikon Eclipse Ti 2 microscope (Nikon Instruments) with the Perfect Focus System, by applying an objective-type TIRF configuration with an oil-immersion objective (Apo SR TIRF 100x, NA 1.49, oil). A 561 nm laser (MPB Communication, 2W, DPSS system) was used for excitation and was coupled into a single-mode fiber. The beam was coupled into the microscope body using a commercial TIRF Illuminator (Nikon Instruments). The laser beam was passed through cleanup filters (ZET561/10, Chroma Technology) and coupled into the microscope objective using a beam splitter (ZT561rdc, Chroma Technology). Fluorescence light was spectrally filtered with two emission filters (ET600/50m, Chroma Technology) and imaged with a sCMOS camera (Andor Zyla 4.2) without further magnification, resulting in an effective pixel size of 130 nm after 2 x 2 binning. Camera readout sensitivity was set to 16-bit, readout bandwidth to 540 MHz.

### Data analysis for DNA-PAINT

#### Determination of TCR ligand positions of divalent DNA origami

Raw imaging data (Tiff image stacks) was subjected to spot-finding and subsequent super-resolution reconstruction using the «Picasso» software package^46^. Drift correction was performed with a redundant cross-correlation and gold nanoparticles as fiducials. For the DNA origami averaging and distance calculation, 49-100 individual structures were picked and averaged using the Picasso «Average» tool. The averaged ligand positions were plotted and fitted using the Origin software. The average distance between two potential ligand positions was then calculated using the distance between the maxima of the fit. Localization precision was determined by NeNA analysis^47^.

#### Determination of nnds within microclusters

Raw imaging data (Tiff image stacks) was subjected to spot-finding and subsequent super-resolution reconstruction using the «Picasso» software package. Drift correction was performed with a redundant cross-correlation and gold nanoparticles as fiducials. Channel alignment from exchange experiments was performed using gold nanoparticles as well. Microclusters were identified either based on the accumulation of localizations of individual P3-Cy3B (for DNA-H57) or P1-Cy3B (for M-H57 and L-H57) imager strand binding events. Individual ligand positions characterized by clouds of localizations were identified within the microclusters. Center positions of localization clouds were determined using a modified Ripley’s K function where the number of neighboring localizations within 10 nm was taken into account and the center position, defined as the localization with the highest number of neighbors, was taken as the determined ligand position. True repetitively visited ligand sites were then identified as exhibiting ≥ 15 localizations within 10 nm distance of the center position. Furthermore, it was required that the mean frame number of all localizations recorded for an individual site was between 20% and 80% of the total number of frames (Supplementary Figure 9). Undesired imager sticking was filtered out by removing all sites harboring more than 80% of the localizations within 5% of frames of the total acquisition. The resulting detected ligand positions were then used to measure nearest neighbor distances (nnds).

### Calcium imaging experiments and analysis

10^6^ T-cells were incubated in T-cell media supplemented with 5 μg ml^−1^ Fura-2 AM (11524766, ThermoFisher Scientific) for 20 min at 24°C. Excessive Fura-2 AM was removed by washing 3x with HBSS + 2% FBS. T-cells were diluted with HBSS + 2% FBS to get a final concentration of 5* 10^3^ cells μl^−1^. 10^5^ cells were transferred to the Lab-Tek™ chamber and image acquisition was started immediately after T-cells landed on the functionalized SLBs. Fura-2 AM was excited using a monochromatic light source (Polychrome V, TILL Photonics), coupled to a Zeiss Axiovert 200M equipped with a 10x objective (Olympus), 1.6x tube lens and an Andor iXon Ultra EMCCD camera. A longpass filter (T400lp, Chroma) and an emission filter were used (510/80ET, Chroma). Imaging was performed with excitation at 340 nm and 380 nm, with illumination times of 50 ms and 10 ms, respectively. The total recording time was 10 min at 1 Hz. Precise temperature control was enabled by an in-house-built incubator equipped with a heating unit. Unless stated otherwise, experiments were carried out at 24°C.

ImageJ was used to generate ratio and sum images of 340nm/380nm. T-cells were segmented and tracked via the sum image of both channels using an in-house Matlab algorithm based on Gao et al.^48^. Cellular positions and tracks were stored and used for intensity extraction based on the ratio image. Intensity traces were normalized to the starting value at time point zero. Traces were categorized in “activating” and non-activating” based on an arbitrary activation threshold ratio of 0.4. The activation threshold was chosen based on comparison of individual traces of a positive control (ICAM-1 100 μm^−2^, B7-1 100 μm^−2^, His-pMHC 150 μm^−2^) and a negative control (ICAM-1 100 μm^−2^, B7-1 100 μm^−2^) (*n* > 40). The percentage of activated cells was evaluated for different ligand surface densities and normalized to the positive control. Data were plotted as % activated cells *A* as a function of ligand surface densities *L* to generate dose-response curves and fitted with Eq. 6 to extract the activation threshold *T_A_*, the maximum response *A_max_* and the Hill coefficient *n*.

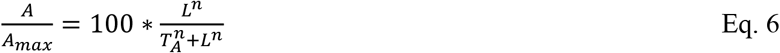

All fit parameters are summarized in **Supplementary Table 3**.

### T-cell fixation and immunostaining

Prior to T-cell seeding, ligand surface densities on SLBs were determined as described above. 2.5 * 10^6^ cells/well were seeded onto SLBs and allowed to settle for 10 min at 24°C. Fixation buffer (PBS, 8% formaldehyde (28908, ThermoFisher Scientific), 0.2% glutaraldehyde (G7776-10ML, Sigma Aldrich), 100 mM Na_3_O_4_V (S6508-10G, Sigma Aldrich), 1 M NaF (S7920-100G, Sigma Aldrich)) was added 1:1 and incubated for 20 min at 24°C, before samples were rinsed with PBS. Permeabilization buffer (PBS, 100 mM Na_3_O_4_V, 1 M NaF, 0.2% Triton-X (85111, ThermoFisher Scientific)) was added and after 1 min samples were washed with PBS before blocking with passivation buffer (PBS, 3% BSA) for 30 min. ZAP-70 antibody labeled with AF647 (clone 1E7.2, #51-6695-82, ThermoFisher Scientific) was added at a final concentration of 1.25 μg ml^−1^ and incubated overnight at 4°C. For experiments with DNA origami MgCl_2_ was added additionally to each buffer at a final concentration of 10 mM to ensure nanoplatforms integrity during fixation and immunostaining. Samples were washed extensively with passivation buffer. Imaging was performed in HBSS.

### Determination of TCR surface densities

Average TCR surface densities were calculated from T-cells in contact with ICAM-1-functionalized SLBs and labeled to saturation with H57-scFv site-specifically labeled with AF555^6^. T-cell brightness per μm^2^ was then divided by the single molecule signal of a single bilayer-anchored His-pMHC-AF555.

### Tissue culture

Primary CD4^+^ T-cells isolated from lymph node or spleen of 5c.c7 *αβ* TCR transgenic mice were pulsed with 1 μM moth cytochrome c peptide (MCC) 88-103 peptide (C18-reverse phase HPLC-purified; sequence: ANERADL<u>IAYLKQATK</u>, T-cell epitope underlined, Elim Biopharmaceuticals Inc, USA) and 50 U ml^−1^ IL-2 (eBioscience) for 7 days to arrive at an antigen-experienced T-cell culture^49^. T-cells were maintained at 37°C and 5% CO2 in RPMI 1640 media (Life technologies) supplemented with 10% FCS (MERCK), 100 μg ml^−1^ penicillin (Life technologies), 100 μg ml^−1^ streptomycin (Life technologies), 2 mM L-glutamine (Life technologies), 0.1 mM non-essential amino acids (Lonza), 1 mM sodium pyruvate (Life technologies) and 50 μM β-mercaptoethanol (Life technologies). Dead cells were removed on day 6 after T-cell isolation by means of density-dependent gradient centrifugation (Histopaque 1119, Sigma). Antigen-experienced T-cells were used for experiments on day 7 – 9.

### Protein expression and purification

We extended a single chain antibody fragment of the variable domain (scF_V_) reactive against the murine TCR*β* chain (mAb clone: H57-597) C-terminally with an AVI tag for site-specific biotinylation followed by a 3C protease cleavable 12x histidine tag. The H57-scF_V_ was expressed in E.coli as inclusion bodies and refolded *in vitro* as described^22^. After refolding, the H57-scF_V_ was concentrated with a stirred 10 kDa ultrafiltration cell (PBGC04310, MERCK and purified by means of gel filtration (Superdex-200 10/300 GE Healthcare Life Sciences) using an ÄKTA pure chromatography system (GE Healthcare Life Sciences). Eluted fractions containing monomeric H57-scF_V_ were concentrated with 10 kDa Amicon^®^ Ultra-4 centrifugal filters (MERCK), treated with a 3C protease (71493, MERCK) to remove the 12x histidine tag and subjected to site-specific biotinylation with a BirA biotin ligase (Acro Biosystems Inc.). 3C protease (containing a 6x histidine tag), BirA biotin ligase (containing a GST-tag) and unprocessed H57-scF_V_ (still containing a 12x histidine tag) were removed by a batch purification with Ni-NTA agarose (ThermoFischer Scientific) and glutathione agarose (ThermoFisher Scientific). The supernatant containing biotinylated H57-scF_V_ was further purified via gel filtration using Superdex 75 (10/300 GL, GE Healthcare Life Sciences). Finally, biotinylated H57-scF_V_s were randomly conjugated on surface-exposed lysines with Alexa Fluor 555 (AF555) carboxylic acid, succinimidyl ester (ThermoFisher Scientific) or Abberior Star 635P (AS635P) carboxylic acid, succinimidyl ester (Abberior) according to the manufacturer’s instructions. To remove excess dye, the AF555- or AS635P-conjugated and biotinylated H57-scF_V_s were purified by gel filtration using Superdex 75 (10/300 GL, GE Healthcare Life Sciences). Fractions containing monomeric, fluorescently-labeled and biotinylated H57-scF_V_ were again concentrated with 10kDa Amicon^®^Ultra-4 centrifugal filters (UFC801024, MERCK) and stored in PBS supplemented with 50% glycerol at −20°C. The protein to dye ratio ranged between 0.93 and 1.0 for the AF555-labeled H57-scF_V_ and was 2.0 for the AS635P-labeled H57-scF_V_ as determined by spectrophotometry (280 to 555 nm ratio).

2xHis_6_-tag pMHC-AF555 was produced as described in^6^. Biotinylated pMHC was produced as follows: The I-E^k^ protein subunits I-E^k^*α*-biotin and I-E^k^β_0_ were expressed as inclusion bodies in E. coli. To arrive at I-E^k^/MCC(ANP)-biotin complexes, I-E^k^*α*-biotin and I-E^k^β_0_ subunits were refolded in vitro in the presence of MCC(ANP) sequence: ANERADLIAYL[ANP]QATK (Elim Biopharmaceuticals Inc) as described^42,50^. Refolded I-E^k^/MCC(ANP)-biotin complexes were purified via 14.4.4S mAb-based affinity chromatography (Cyanogen bromide-activated Agarose, MERCK) followed by gel filtration (Superdex-200 10/300 GE Healthcare Life Sciences). Eluted protein fractions were site-specifically biotinylated using the BirA (Acro Biosystems Inc.) and subjected to a batch purification step with glutathione agarose (ThermoFisher Scientific) to remove the BirA ligase and gel filtration (Superdex-200 10/300 GE Healthcare Life Sciences). Fractions containing monomeric I-E^k^/MCC(ANP)-biotin complexes were concentrated with 10kDa Amicon^®^Ultra-4 centrifugal filters (UFC801024, MERCK), snap frozen in liquid nitrogen and stored in 1x PBS at −80 °C. The MCC peptide derivative MCC-C (sequence: ANERADLIAYLKQATKGGSC) was purchased from Elim Biopharmaceuticals, purified via reversed phase chromatography (Pursuit XRs C18 column, Agilent) and conjugated to AF555 C2-maleimide (ThermoFisher Scientific) according to the manufacturer’s instructions. Purity of the MCC-C peptide and efficient fluorophore coupling to AF555 C2-maleimide was verified via MALDI-TOF (Bruker). For fluorescence labeling of pMHC, we exchanged the placeholder peptide MCC(ANP) for site-specifically AF555-conjugated MCC peptides (MCC-AF555) under acidic conditions (1x PBS, 100mM citric acid, pH5.1) for 72 hours at room temperature. Following peptide exchange, AF555-conjugated I-E^k^/MCC complexes (either C-terminally extended with 2xHis_6_ or biotin) were subjected to gel filtration (Superdex-200 10/300 GE Healthcare Life Sciences) to remove an excess of unreacted MCC-AF555. Fractions containing monomeric AF555-conjugated I-E^k^/MCC complexes were concentrated with 10kDa Amicon^®^Ultra-4 centrifugal filters and stored in 1x PBS supplemented with 50% glycerol at −20 °C. Quantitative I-E^k^/MCC[ANP] peptide replacement (>99%) was verified via spectrophotometry (280 to 555 nm ratio).

Trans-divalent streptavidin (dSA) was prepared with some adaptions as described^20^. The pET21a (+) vectors encoding “alive” (i.e. biotin binding) and “dead” (i.e. biotin non-binding) streptavidin subunits were kindly provided by Alice Ting (Stanford University, USA). We substituted the 6x histidine tag of the “alive” subunit with a cleavable 6x glutamate tag to allow for purification via cation exchange chromatography preceded by a recognition site of the 3C protease for optional removal of the tag^7^. Both, “alive” and “dead” streptavidin subunits were expressed in *E. coli* (BL-21) for 4 h at 37°C and refolded from inclusion bodies as described^19^. After refolding, the streptavidin tetramer mixture was concentrated in a stirred 10kDa ultrafiltration cell (PBGC04310, MERCK). Further concentration and buffer exchange to 20 mM Tris-HCl pH 8.0 were carried out with 10kDa Amicon^®^Ultra-4 centrifugal filters (MERCK). The mixture of tetramers was then purified by anion exchange chromatography (MonoQ 5/50 GE Healthcare Life Sciences) using a column gradient from 0.1 to 0.4 M NaCl. dSA was eluted with 0.25 M NaCl, concentrated again (10kDa Amicon^®^Ultra-4 centrifugal filters) and further purified via gel filtration (Superdex-200 10/300, GE Healthcare Life Sciences). After removal of the poly-E tag with the 3C protease (MERCK), the protein was again subjected to gel filtration (Superdex-200 10/300 GE Healthcare Life Sciences). Monomeric fractions of dSA were then concentrated (10kDa Amicon^®^Ultra-4 centrifugal filters) and stored in PBS supplemented with 50% glycerol at −20°C.

Monovalent streptavidin (mSA) was produced as described above with some adaptations. The sequence of the “dead” subunit was C-terminally extended with a 3xHis_6_-tag for attachment to lipid bilayers containing 18:1 Ni-DOGS NTA. After expression in E. coli and refolding from inclusion bodies *in vitro*, the mixture of tetramers was purified by anion exchange chromatography (MonoQ 5/50 GE Healthcare Life Sciences) using a column gradient from 0.1 to 0.4 M NaCl. mSA was eluted with 0.22 M NaCl, concentrated again (10kDa Amicon^®^Ultra-4 centrifugal filters) and further purified via gel filtration (Superdex-200 10/300, GE Healthcare Life Sciences). After removal of the poly-E tag located on the “alive” subunit with 3C protease (MERCK) followed by a gel filtration step (Superdex-200 10/300 GE Healthcare Life Sciences), the protein was stored in PBS supplemented with 50% glycerol at −20°C.

### Animal model and ethical compliance statement

5c.c7 αβ TCR-transgenic mice bred onto the B10.A background were a kind gift from Björn Lillemeier (Salk Institute, USA). Animal husbandry, breeding and sacrifice for T-cell isolation was evaluated by the ethics committees of the Medical University of Vienna and approved by the Federal Ministry of Science, Research and Economy, BMWFW (BMWFW-66.009/0378-WF/V/3b/2016). They were performed in accordance to Austrian law (Federal Ministry for Science and Research, Vienna, Austria), the guidelines of the ethics committees of the Medical University of Vienna and the guidelines of the Federation of Laboratory Animal Science Associations (FELASA), which match those of Animal Research: Reporting *in vivo* Experiments (ARRIVE). Further, animal husbandry, breeding and sacrifice for T-cell isolation was conducted under Project License (I4BD9B9A8L) which was evaluated by the Animal Welfare and Ethical Review Body of the University of Oxford and approved by the Secretary of State of the UK Home Department. They were performed in accordance to Animals (Scientific Procedures) Act 1986, the guidelines of the ethics committees of the Medical Science of University of Oxford and the guidelines of the Federation of Laboratory Animal Science Associations (FELASA), which match those of Animal Research: Reporting in vivo Experiments (ARRIVE). Both male and female mice at 8-12 weeks old were randomly selected and sacrificed for isolation of T-cells from lymph nodes and spleen.

## Code availability

Software used to analyze single molecule brightness and diffusion, as well as for DNA-PAINT data filtering and position detection are available from the corresponding author on request.

## Data availability

The datasets generated during and/or analyzed during the current study are available from the corresponding author on request.

## ACKNOWLEDGEMENTS

This work was supported by the Austrian Science Fund (FWF project V538-B26, ES and JH; the PhD program Cell Communication in Health and Disease W1205, RP, JBH and HS), the TU Wien doctoral college BioInterface (JH), the European Fund for Regional Development (EFRE, IWB2020, AK and JP), the Federal State of Upper Austria (JP), the Vienna Science and Technology Fund (WWTF, LS13-030, GJS and JBH), the Boehringer Ingelheim Fonds (RP), the German Research Foundation through the Emmy Noether Program (DFG JU 2957/1-1, RJ), the SFB1032 (project A11, RJ), the European Research Council through an ERC Starting Grant (MolMap; grant agreement number 680241, RJ), the Allen Distinguished Investigator Program through The Paul G. Allen Frontiers Group (tRJ), the Danish National Research Foundation (Centre for Cellular Signal Patterns, DNRF135, RJ), the Human Frontier Science Program through a Young Investigator Grant (HFSP RGY0065), the Max Planck Foundation (RJ) and the Max Planck Society (to RJ), the International Max Planck Research School for Molecular and Cellular Life Sciences (IMPRS-LS)(ASE), the German Research Foundation through the QBM graduate school (TS) and the Wellcome Trust (Principal Research Fellowship 100262 Z/12/Z, EK). We thank V. Mühlgrabner for help with tissue culture.

## AUTHOR CONTRIBUTIONS

ES and JH conceived and designed the research; JH conducted most experiments and performed the analysis. MB wrote Matlab codes and assisted with analysis; VM and VB performed experiments to determine DNA origami occupancy; RP and JBH provided proteins and T-cells, EK provided T-cells, JBH, HS and GJS advised on experimental design and discussed results; AK and JP performed and analyzed HS-AFM measurements; ASE, TS and RJ performed and analyzed DNA-PAINT measurements. ES wrote the manuscript. All authors commented on the manuscript.

