## Supplementary Information for "DNA origami demonstrate the unique stimulatory power of single pMHCs as T-cell antigens"

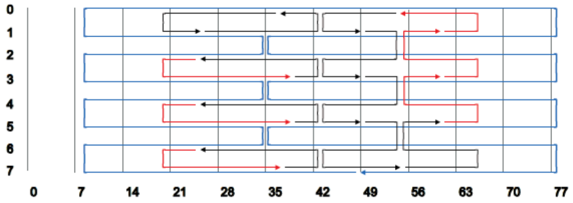
0
1
2
3
4
5
6
7
0 7 14 21 28 35 42 49 56 63 70 77

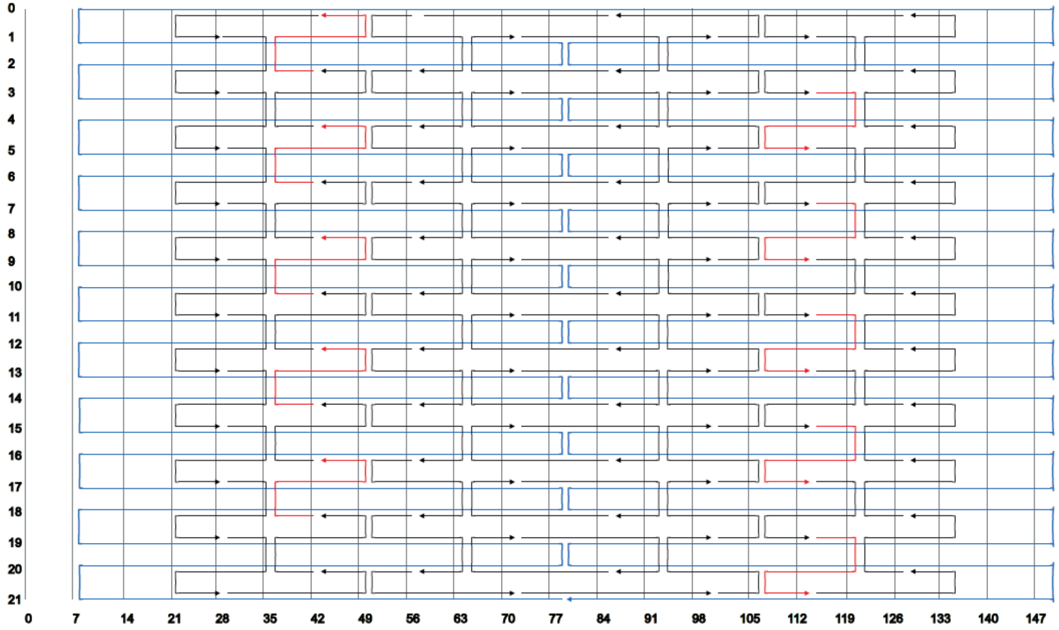
0 7 14 21 28 35 42 49 56 63 70 77 84 91 98 105 112 119 126 133 140 147

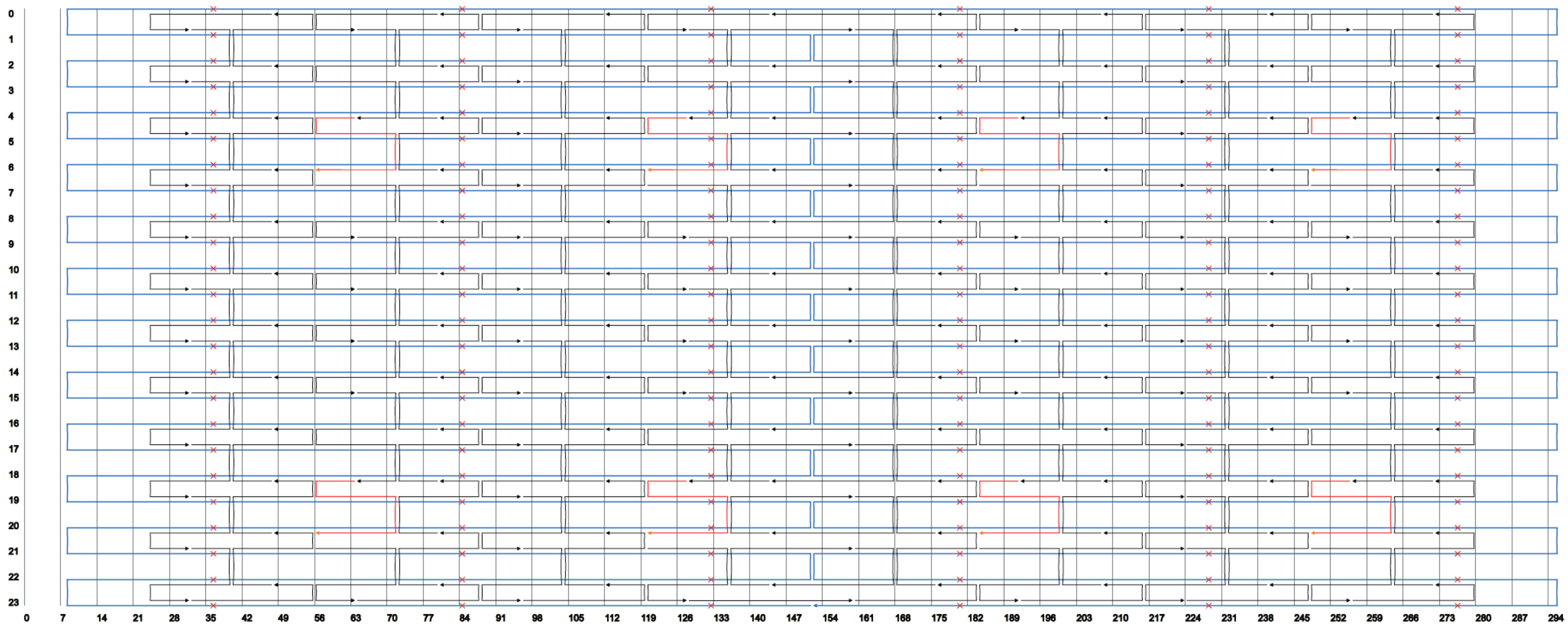

**Supplementary Fig. 1: DNA origami scaffold routing.** DNA origami (30x20 nm, 65x54 nm, 100x70 nm[1]) were designed based on the M13mp18 scaffold[2] using caDNAno[3]. At sites chosen for ligand attachment, staple strands were elongated at their 3'-end with 21 bases. For attachment to the SLB via complementary cholesterol-oligonucleotides staple strands were elongated at predefined distinct positions at their 5'-end with 25 bases (red arrows).

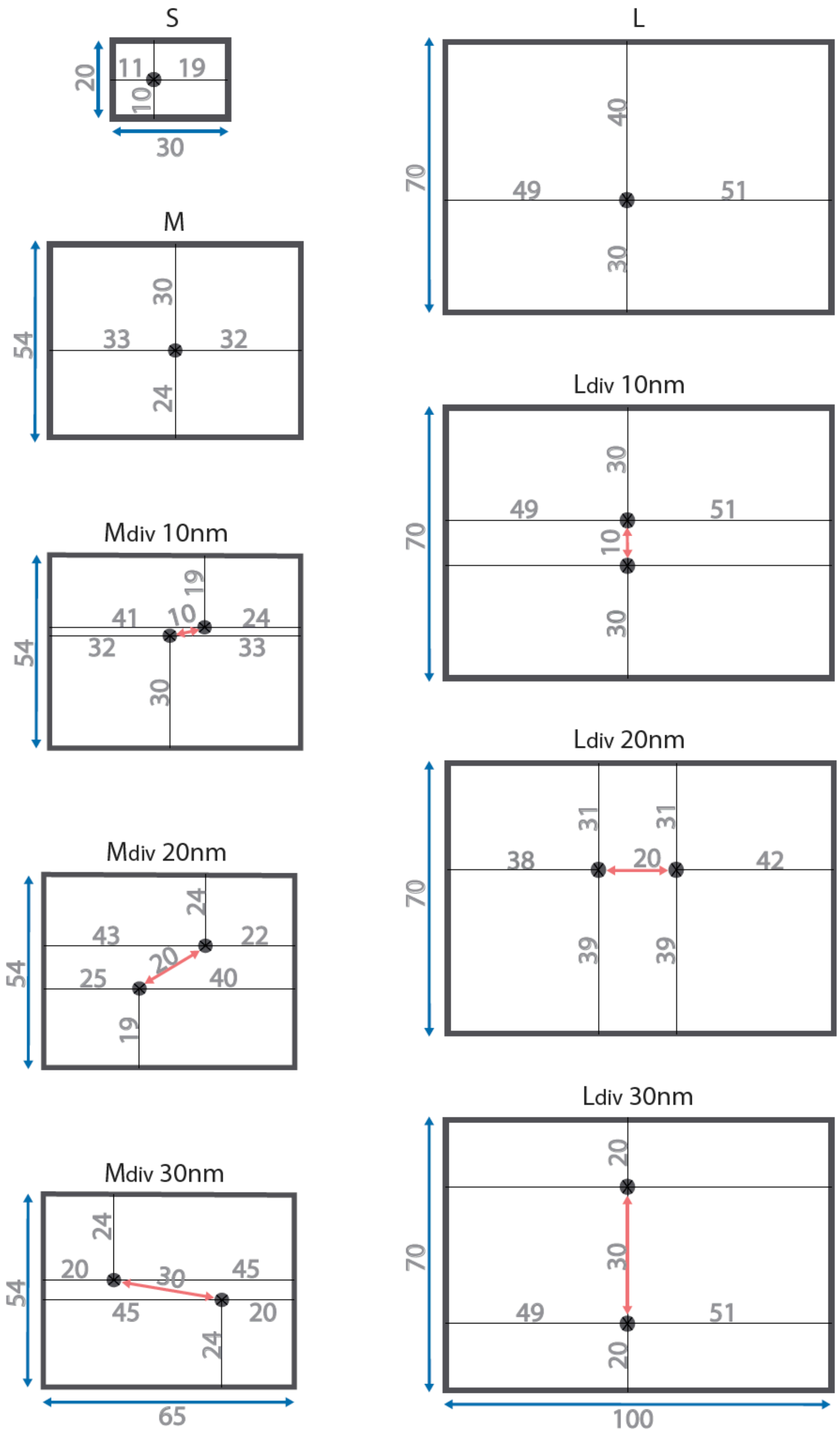

**Supplementary Fig. 2: Schematics of DNA origami used in this study.** Black dots indicate modification sites on the top side of DNA origami platforms. The distances are approximated with 0.34 nm per base pair along the helical axis and with 2 nm per helix perpendicular to the helical axis[2]. Interhelical gaps were assigned 1 nm, depending on the spacing of crossovers between helices. Distances are given in nm.

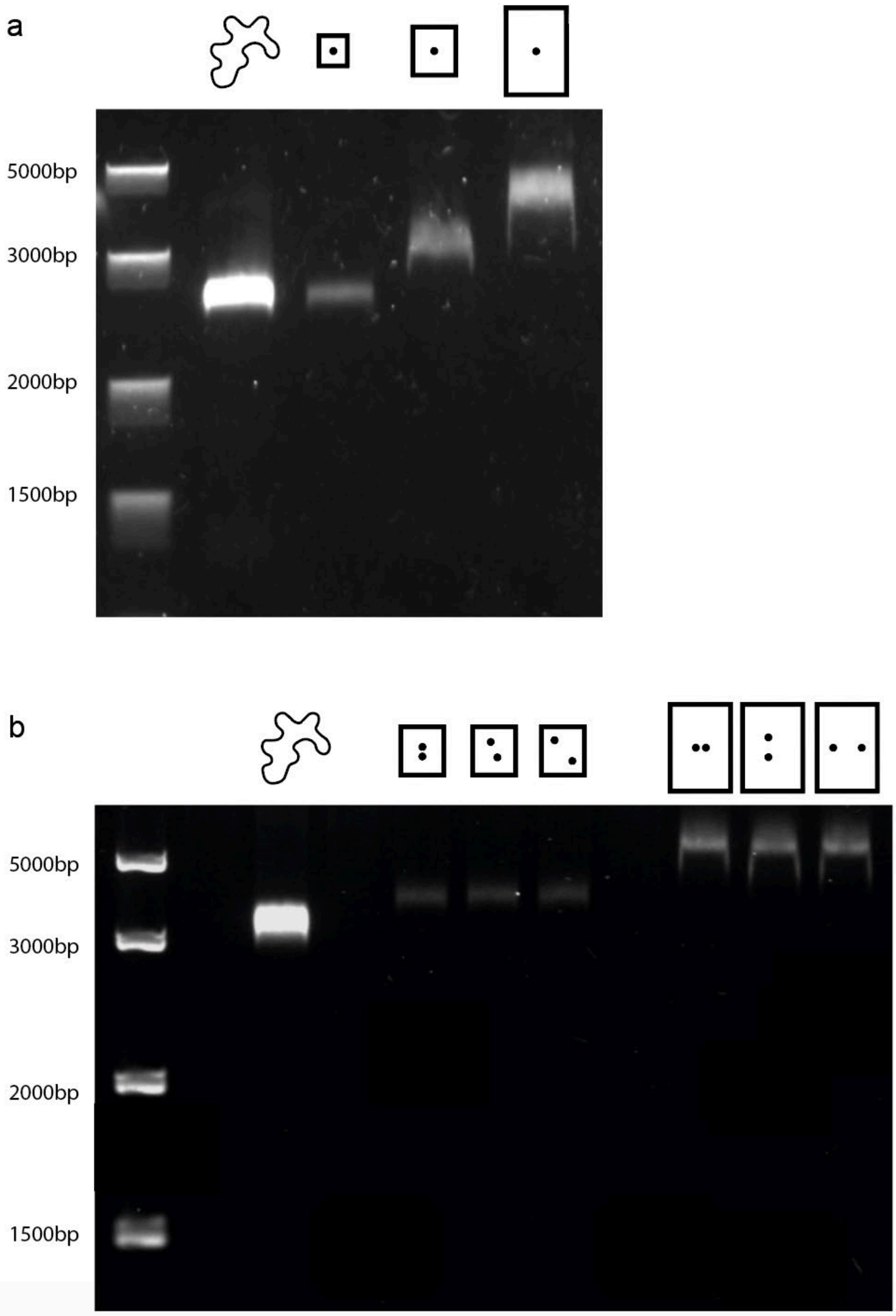

**Supplementary Fig. 3: Agarose gel electrophoresis of DNA origami platforms.** A 1% agarose gel (1xTAE, 10 mM $MgCl_2$) pre-stained with Sybr$^{TM}$ gold nucleic acid gel stain was run at 100V for 75 min. A FastRuler middle range DNA ladder was used as a reference. Schematic sketches above the individual bands indicate the different DNA origami layouts functionalized with dSA. **a,** from left to right: M13mp18 scaffold, S, M, L. **b**, M13mp18 scaffold, $M_{div}$ 10nm, $M_{div}$ 20nm, $M_{div}$ 30nm, $L_{div}$ 10nm, $L_{div}$ 20nm, $L_{div}$ 30nm.

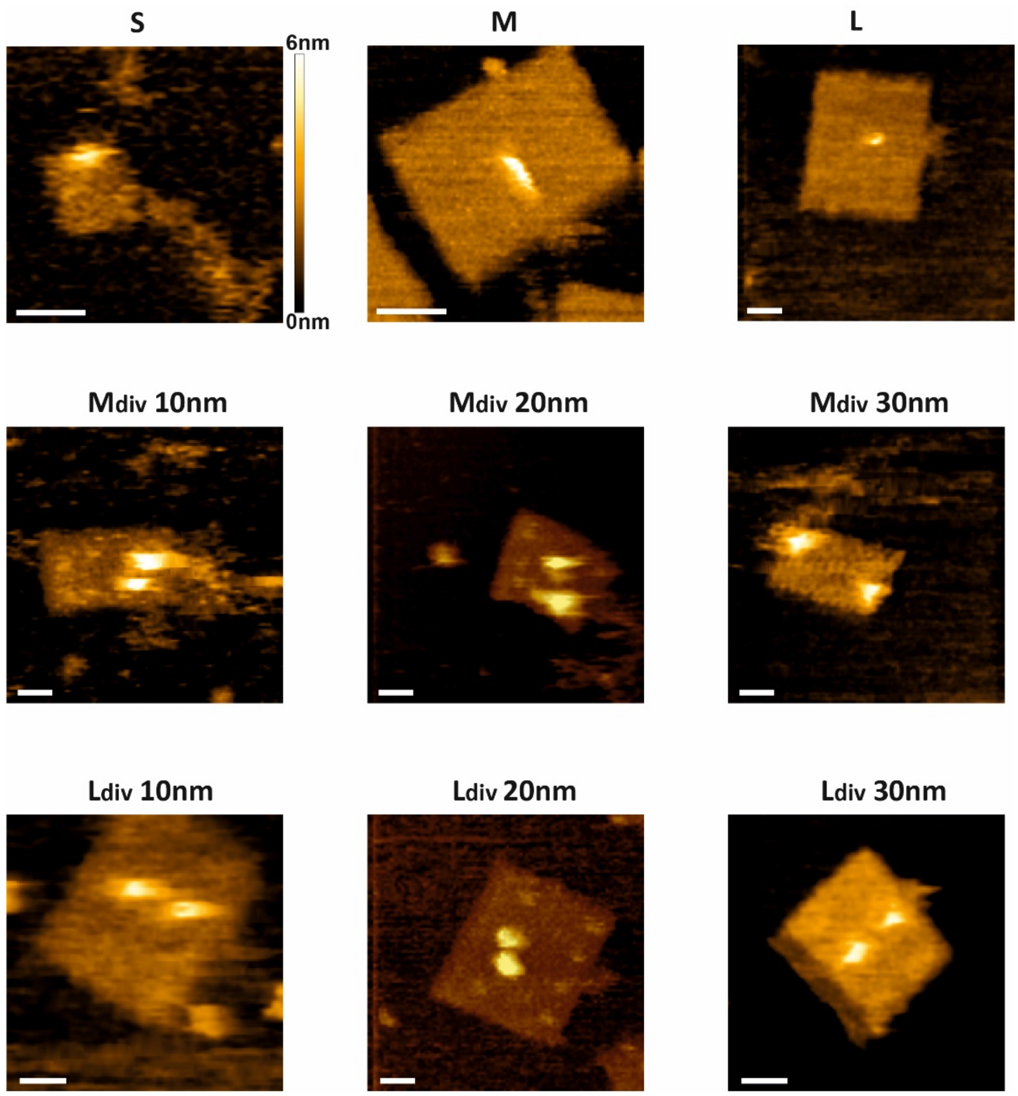

**Supplementary Fig. 4: AFM images of functionalized DNA origami platforms.** The different DNA origami platforms used in this study were functionalized with dSA as indicated in Supplementary Fig.2 and imaged using highspeed-AFM (HS-AFM). Scale bar, 25 nm.

**Note**

Unfolded scaffold, which is unstructured and highly flexible, is imaged as a blur with HS-AFM. While almost all of the M13mp18 Phage DNA is used to fold the large "L" DNA origami structure, half ("M" DNA origami) or 90% ("S" DNA origami) remain unfolded. Note that the divalent streptavidin used for attachment of biotinylated TCR-ligands was placed on top of a ~5nm linker. During AFM imaging, the AFM tip pushes the streptavidin molecule towards the DNA origami platform surface, sometimes even ripping it off from the structure, thereby distorting the distances.

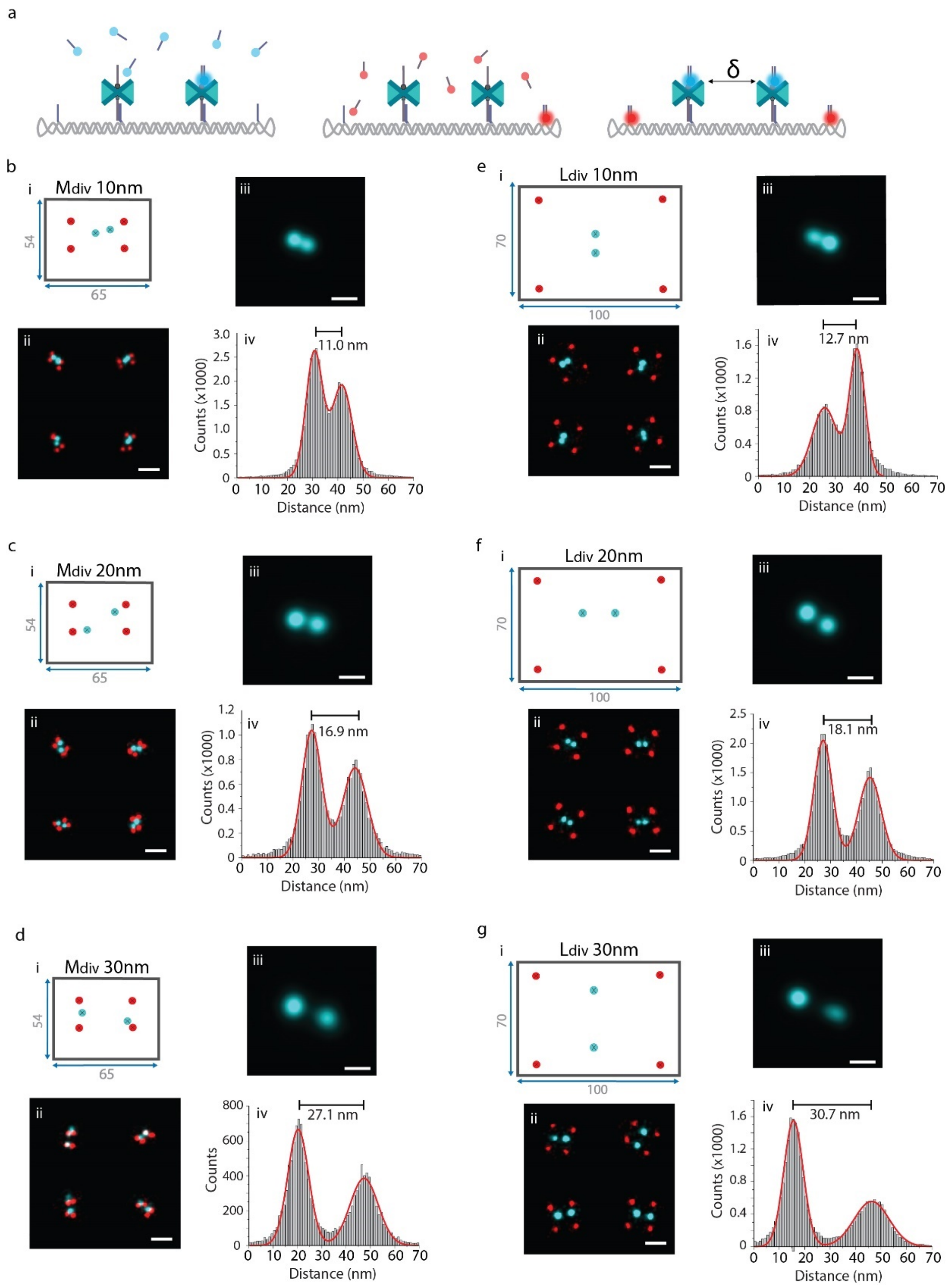

**Supplementary Fig. 5: Mapping ligand positions on divalent DNA origami platforms via DNA-PAINT super-resolution microscopy. a,** Principle of DNA-PAINT. Freely diffusing dye-labeled oligonucleotides ("imager" strands) transiently bind to their target-bound complements ("docking" strands) to achieve the necessary target "blinking" for stochastic super-resolution microscopy. For this, biotinylated TCR ligands were replaced with biotinylated DNA-PAINT docking strands (cyan) on divalent DNA origami platforms. Additionally, four staple strands at the corners were extended with DNA-PAINT docking sites as platform markers (red). Ligand (cyan) and platform (red) positions were imaged consecutively by Exchange-PAINT[4] using Cy3B imager strands. **i,** DNA origami layouts showing ligand and platform positions for the following constructs: **b,** $M_{div}$ 10nm, **c,** $M_{div}$ 20nm, **d,** $M_{div}$ 30nm, **e,** $L_{div}$ 10nm, **f,** $L_{div}$ 20nm, **g,** $L_{div}$ 30nm. **ii,** 4 exemplary pseudo-color DNA-PAINT super-resolution images of DNA origami platforms. Scale bar, 50 nm. **iii,** DNA-PAINT sum images of ligand positions, $n \geq 49$. Scale bar, 20 nm. **iv,** Intensity profiles across ligand positions indicated in (iii).

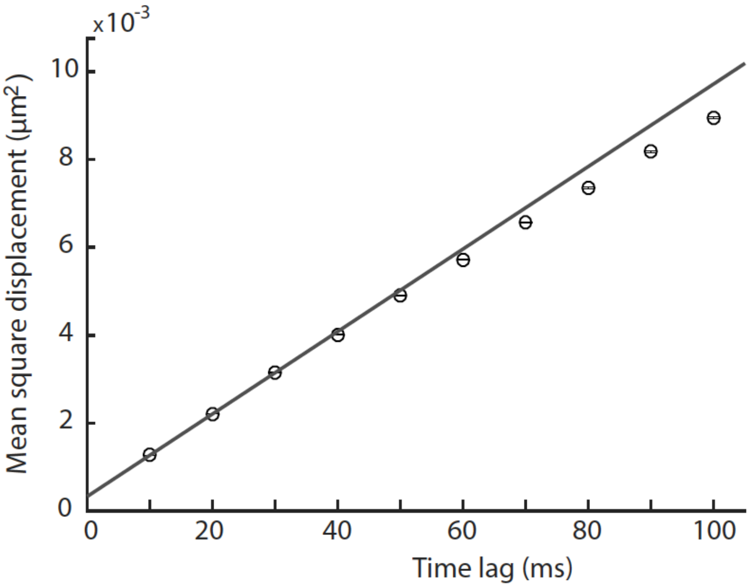

**Supplementary Fig. 6: Determination of diffusion behavior of DNA origami on SLBs.** Diffusion constants were evaluated by performing single-molecule tracking experiments. Mean square displacements were determined and plotted as a function of time lags. Assuming pure Brownian motion, the diffusion coefficient D was derived by fitting the first two data points with a linear fit. An exemplary plot for M-H57 is shown.

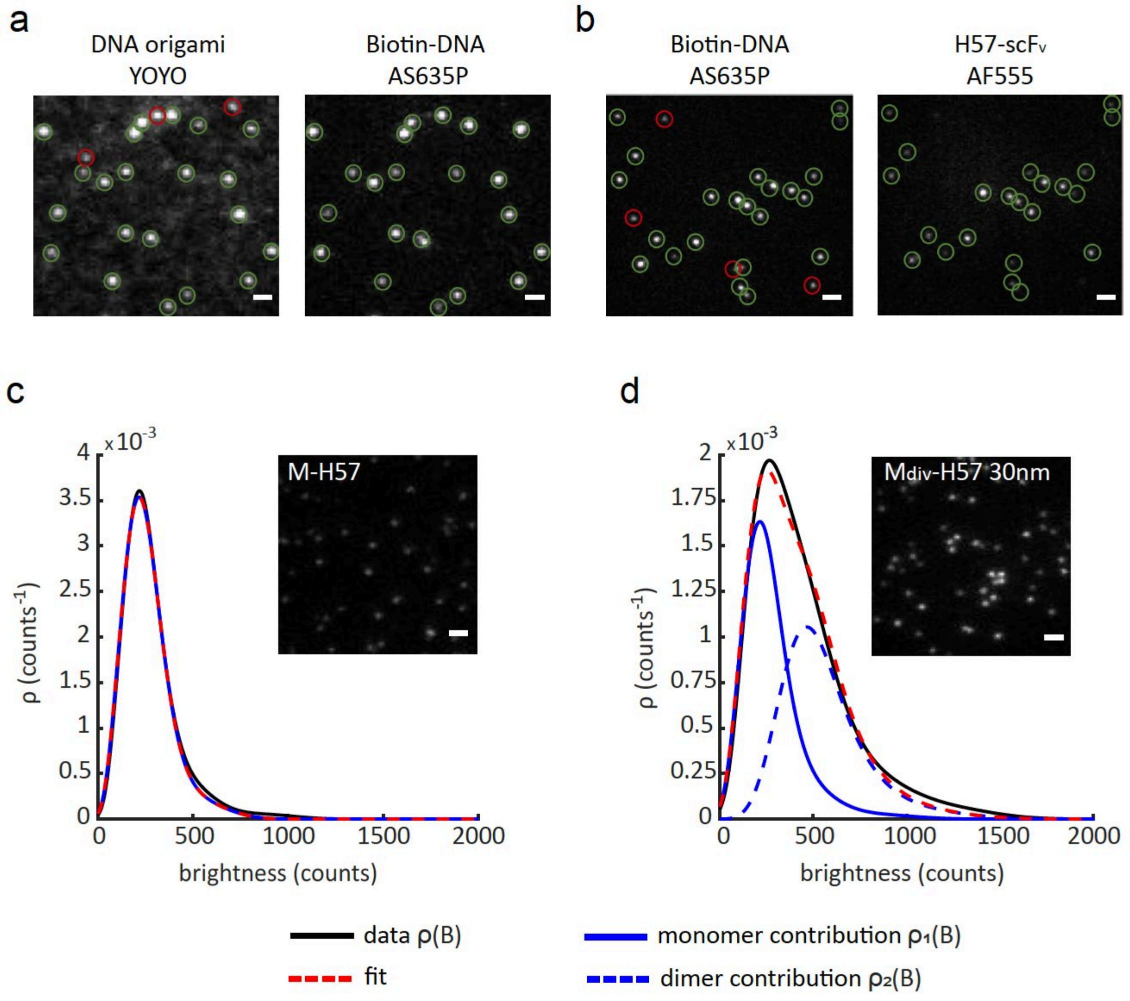

**Supplementary Fig. 7: Determination of DNA origami ligand occupancy. a,** To determine the fraction of attachment sites on DNA origami available for modification, elongated staple strands were hybridized with fluorescently labeled biotinylated oligos (Biotin-DNA-AS635P) and two-color colocalization with DNA origami, pre-stained with YOYO, was assessed. Exemplary TIRF images of $M_{div}$ 30nm DNA origami on a SLB are shown. Green open circles indicate signals detected in both color channels; red open circles indicate signals detected only in one channel. **b**, We quantified the colocalization of hybridized biotin-DNA-AS635P and AF555-labeled H57-scF$_{VS}$ to determine the functionalization efficiency of existing biotin modifications with ligand (here shown for $M_{div}$-H57$_{30nm}$). **c, d,** Exemplary TIRF images and corresponding brightness distributions $\rho_n$ of DNA origami on SLBs decorated with AF555-labeled H57-scF$_V$. Data for medium-sized DNA origami bearing 1 (M-H57, 800 signals) **(c)** or 2 ($M_{div}$-H57$_{30nm}$, 708 signals) **(d)** possible ligand attachment sites are shown. The detected signals were fitted and deconvolved into monomer and dimer contributions (see methods section)[5]. Scale bar, 2 µm.

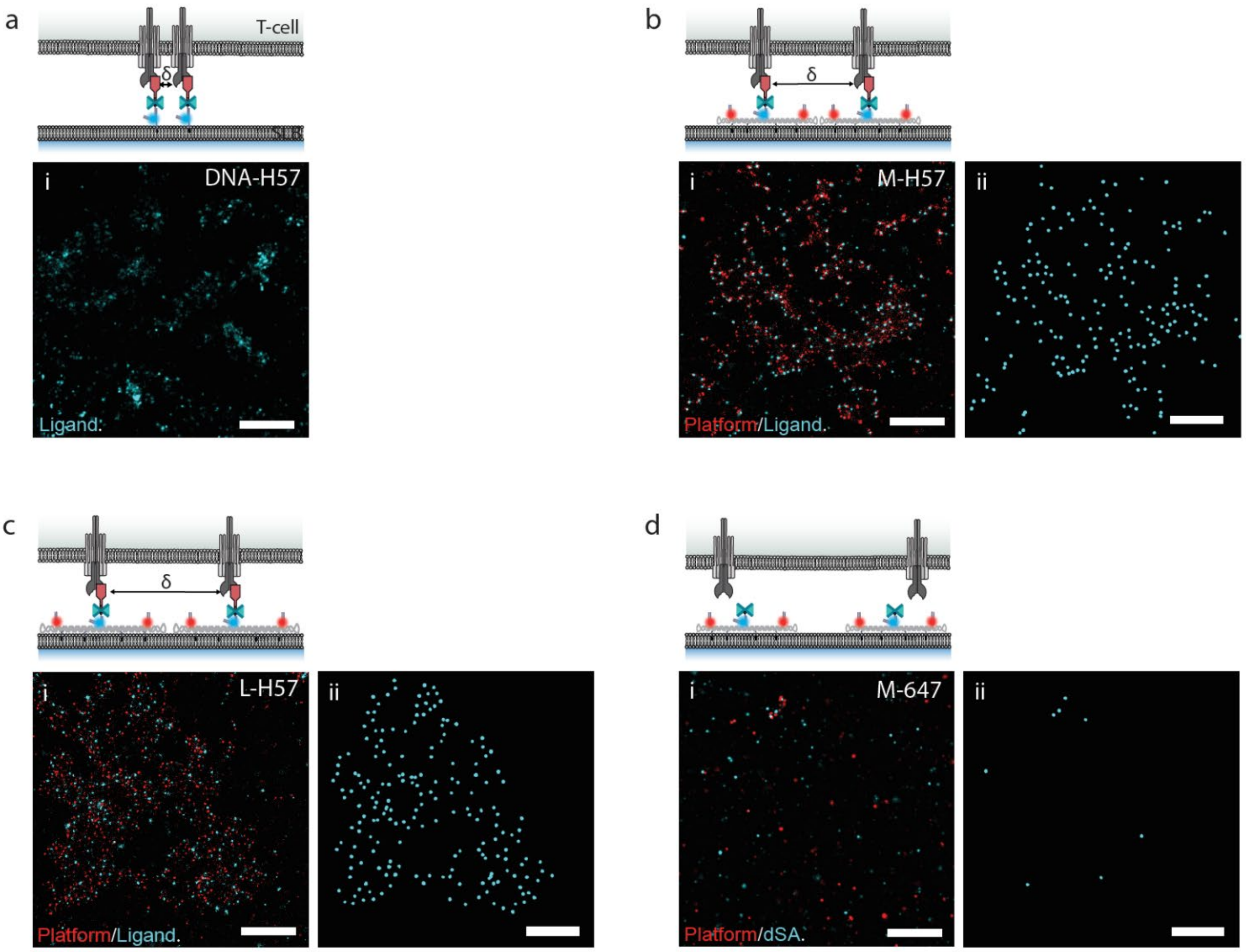

**Supplementary Fig. 8: DNA-PAINT super-resolution imaging of DNA origami platforms within microclusters.** DNA origami platforms impose minimum distances δ on ligands in microclusters. For DNA-PAINT, biotinylated oligos were extended with DNA-PAINT docking sites at their 3' end for imaging ligand positions (cyan). Additionally, four staple strands at the platform corners were extended with DNA-PAINT docking sites to serve as platform markers (red). **a,** DNA-anchored H57-scF$_{VS}$ free to move without restrictions (DNA-H57, δ~5 nm), **b**, H57-scF$_{VS}$ on medium-sized DNA origami platforms (M-H57, δ=48 nm), **c**, H57-scF$_{VS}$ on large DNA origami platforms (L-H57, δ=60 nm), **d**, medium-sized DNA origami platforms without ligands labeled with AF647 (M-647).

**i**, Exemplary pseudo-color DNA-PAINT super-resolution images of H57-decorated constructs underneath fixed T-cells. Ligand (cyan) and marker (red) positions were imaged consecutively by Exchange-PAINT using Cy3b imager strands. Characteristic microcluster formation could be observed for all constructs except for DNA origami platforms without ligands. **ii**, Detected ligand positions after filtering. Ligand positions from **i** were classified as «true» or «false» based on their blinking statistics (see Methods section and Supplementary Fig. 9). Note that individual positions of the freely arrangeable DNA-H57 could not be resolved via DNA-PAINT at the experimental conditions applied. Scale bars, 500 nm.

**Note on DNA super-resolution imaging of S-H57 platforms**

The smallest DNA origami platform (S-H57, 30x20 nm) could not be successfully imaged via DNA-PAINT due to undesired binding events to the structures. This is likely caused by transient hybridization of the DNA imager strand to the unfolded scaffold (note that for the S DNA origami platform only 10% of the scaffold was used for the platform design).

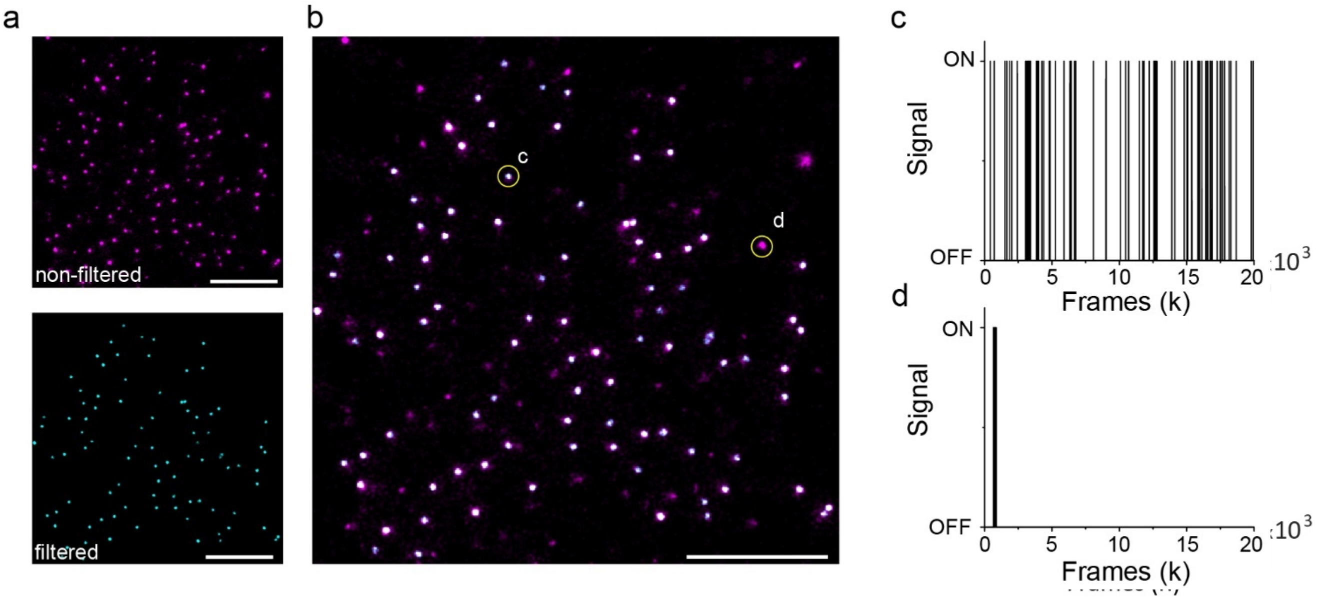

**Supplementary Fig. 9: Filtering of DNA-PAINT data**. **a**, An exemplary DNA-PAINT super-resolution image is shown with non-filtered (magenta) and filtered data (cyan). **b,** Overlay of images from **(a)**. Signals arising from specific binding (after filtering) classified as «true» localizations are shown in white. **c**, Specific binding (indicated in **b)** is characterized by repetitive ($n \geq 15$) single-molecule detection events within a 10 nm radius due to transient binding of the imager strand to a complementary docking site. **d**, Non-specific binding (indicated in **(b)** is characterized by a single binding event of longer duration. These non-specific signals were filtered out and discarded for further analysis. Scale bar, 500 nm

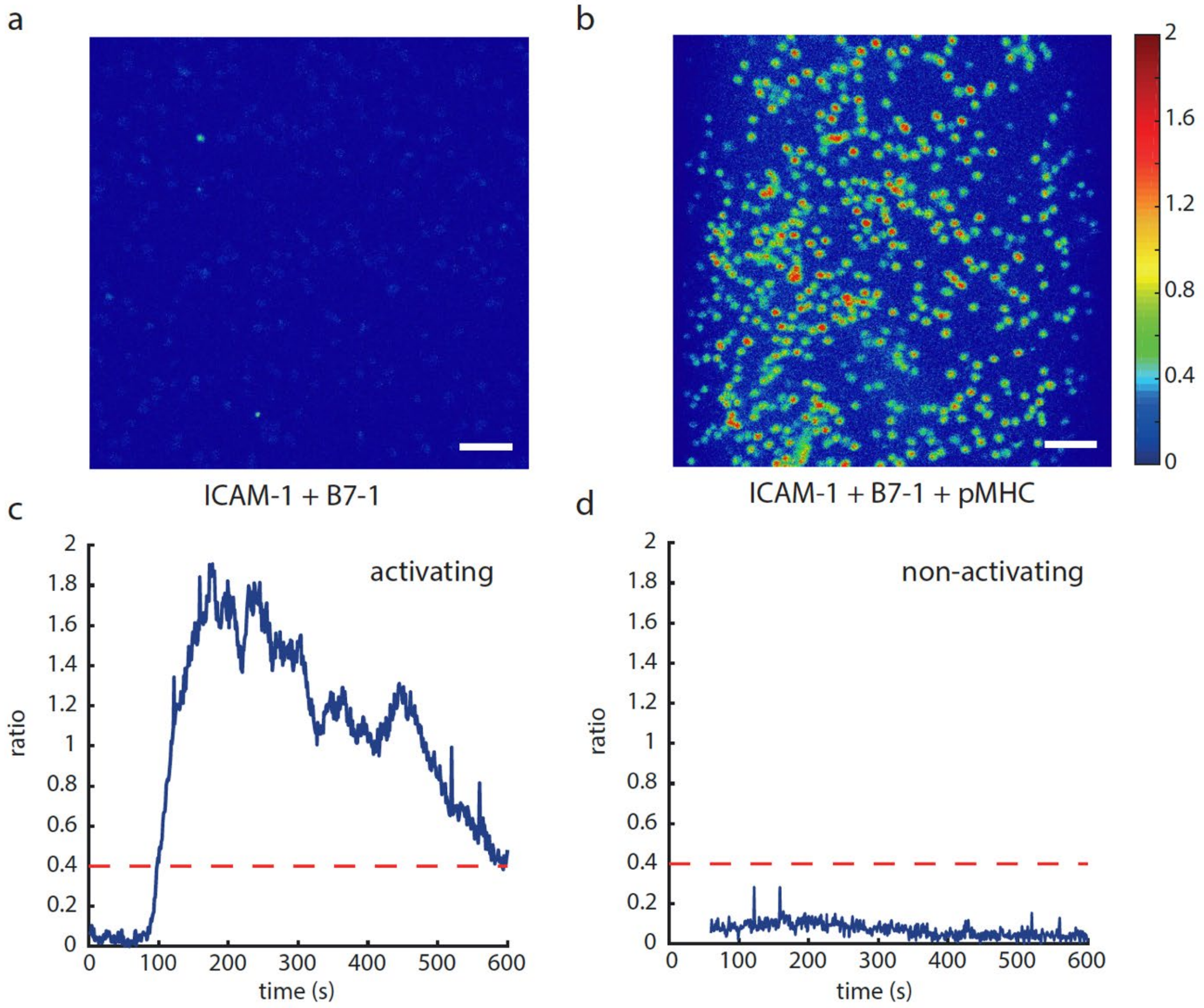

**Supplementary Fig. 10: Calcium imaging experiments to assess the T-cell activation state.** T-cells were loaded with the ratiometric $Ca^{2+}$-sensitive dye Fura-2 AM, seeded onto SLBs and fluorescence emission was recorded at excitation wavelengths 340 nm and 380 nm over 10 min. Activation was tracked via a change of the intensity ratio (340/380nm). Exemplary ratio images recorded at activating (ICAM-1 100 $\mu m^{-2}$, B7-1 100 $\mu m^{-2}$, pMHC 150 $\mu m^{-2}$) **(a)** and non-activating (ICAM-1 100 $\mu m^{-2}$, B7-1 100 $\mu m^{-2}$) **(b)** conditions are shown 5 min after cell seeding. **c,d,** For each cell, the intensity ratio 340/380nm was determined and plotted over time. Exemplary calcium traces for a T-cell under activating **(c)** and non-activating **(d)** conditions are shown. The threshold ratio for counting a cell as “activated” was set to 0.4 for all experiments, indicated by a red dashed line. Scale bar, 4 µm.

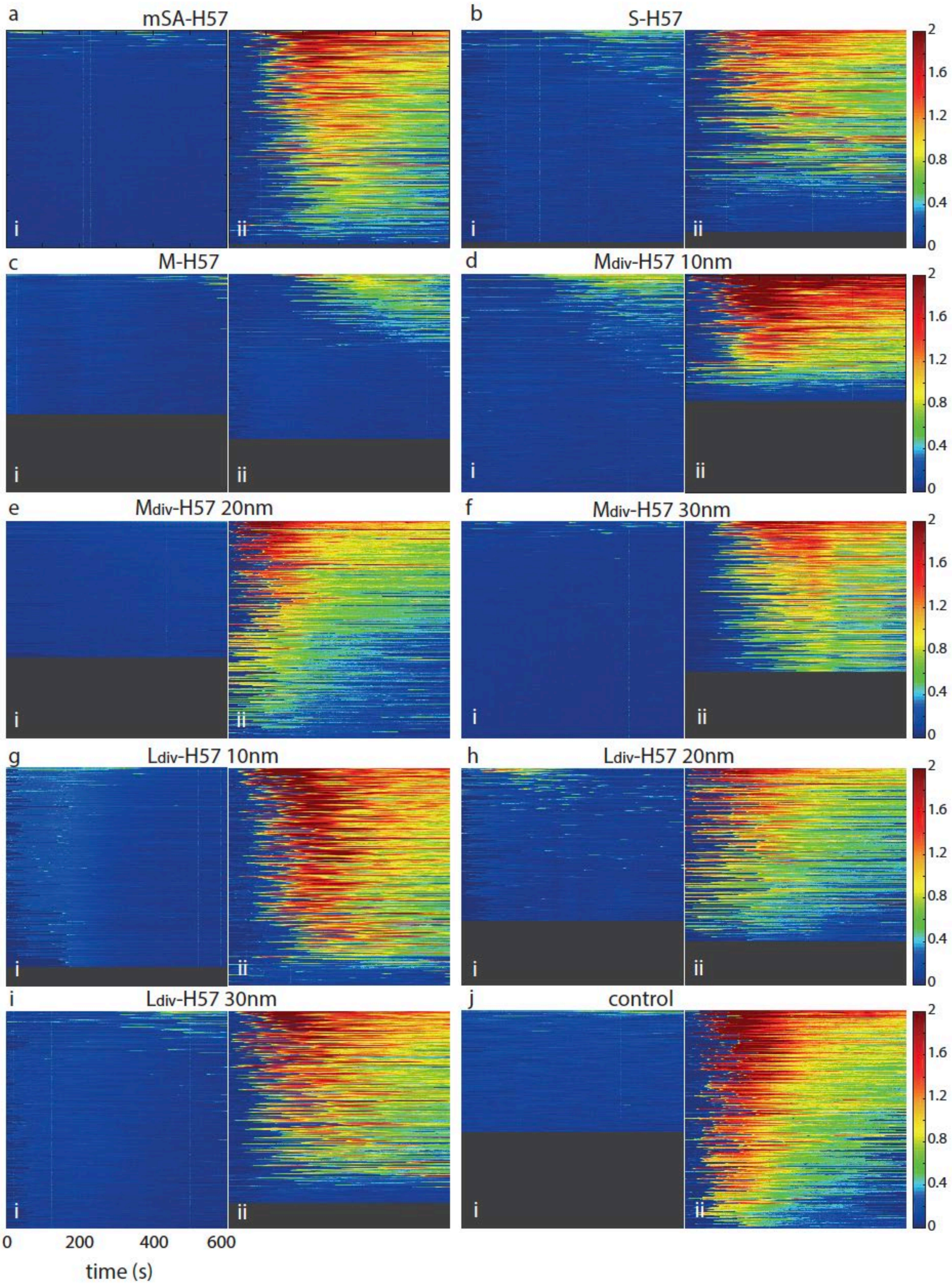

**Supplementary Fig. 11: Population calcium flux on H57-scF$_V$ biointerfaces.** Pseudo-colored calcium flux (Fura-2 AM ratio) traces of >100 cells per region for: **a**, mSA-H57, **b**, S-H57, **c**, M-H57, **d**, M$_{div}$-H57$_{10nm}$, **e**, M$_{div}$-H57$_{20nm}$, **f**, M$_{div}$-H57$_{30nm}$, **g**, L$_{div}$-H57$_{10nm}$, **h**, L$_{div}$-H57$_{20nm}$, **i,** L$_{div}$-H57$_{30nm}$ at non-activating (**i**) and activating conditions (**ii**). For M-H57, traces at the highest measured ligand density of 248 µm$^{-2}$ are shown. **j**, A negative control containing ICAM-1 and B7-1 at 100 µm$^{-2}$ and a positive control containing ICAM-1 and B7-1 at 100 µm$^{-2}$, and His-pMHC at 150 µm$^{-2}$ are shown as a reference. Each horizontal line represents one cell. Cell traces are ranked by their integrated calcium flux in descending order. Calcium was imaged at 1 Hz for 10 min at 24°C.

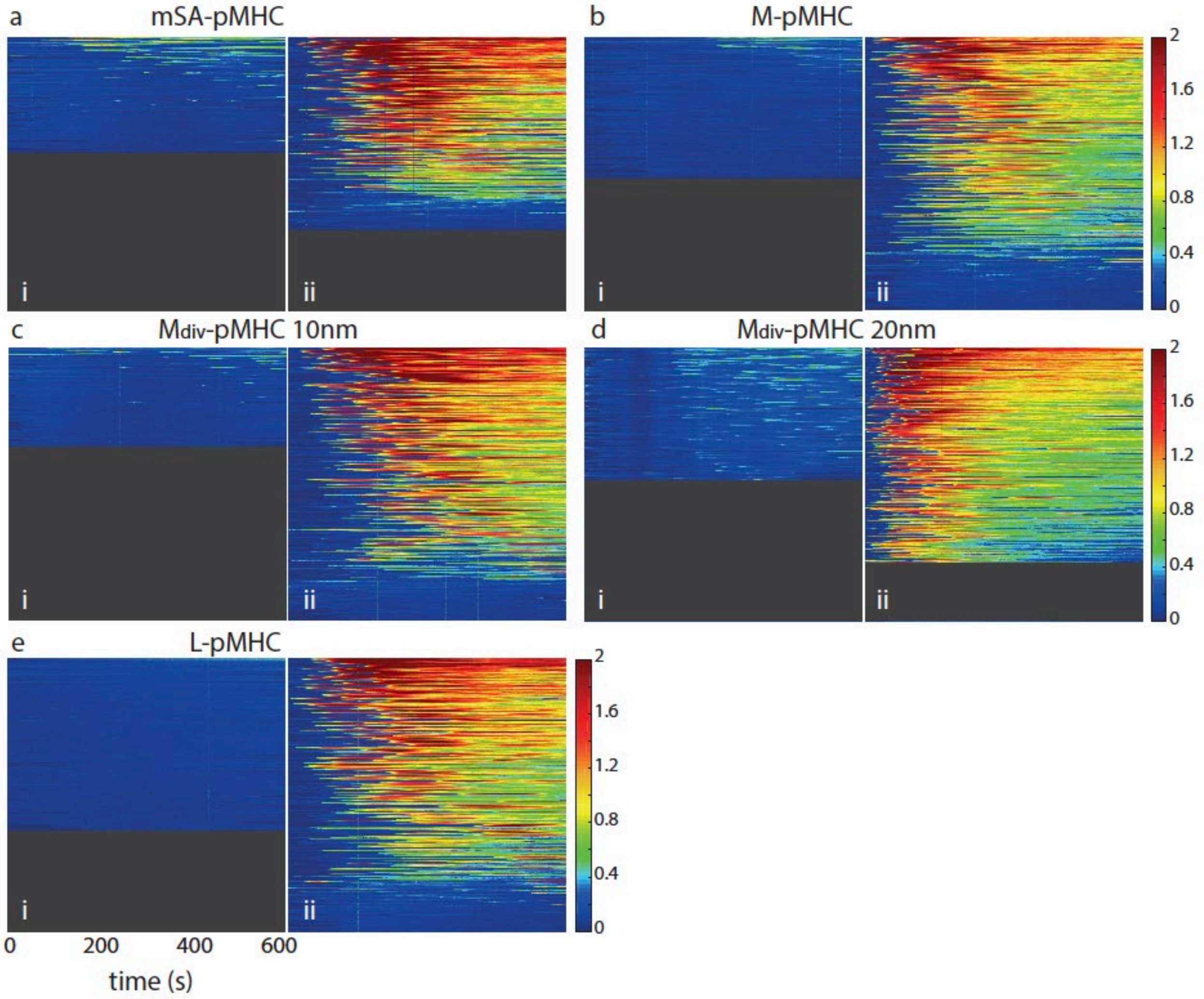

**Supplementary Fig. 12: Population calcium flux on pMHC biointerfaces.** Pseudo-colored calcium flux (Fura-2 AM ratio) traces of >100 cells per region for: **a**, mSA-pMHC, **b**, M-pMHC, **c**, $M_{div}$-$pMHC_{10nm}$, **d**, $M_{div}$-$pMHC_{20nm}$, **e**, L-pMHC) at non-activating (**i**) and activating conditions (**ii**) are shown. Each horizontal line represents one cell. Cell traces are ranked by their integrated calcium flux in descending order. Calcium was imaged at 1 Hz for 10 min at 24°C.

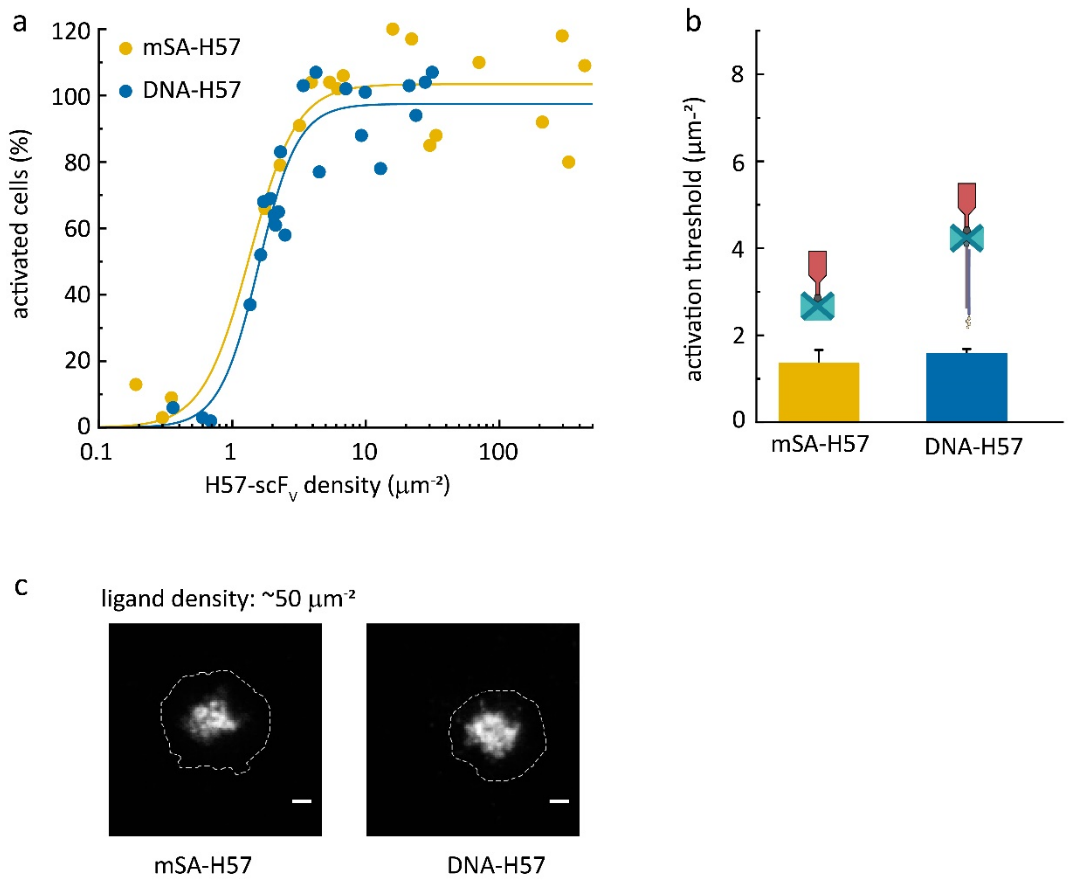

**Supplementary Fig. 13: Comparison of signaling potency of mSA-H57 and DNA-H57. a**, Percentage of activated T-cells at different ligand surface densities are shown. Data were normalized (=100%) to a positive control containing ICAM-1 and B7-1 at 100 µm$^{-2}$, and His-pMHC at 150 µm$^{-2}$. Each data point represents the population average of 294 ± 106 cells (mean ± s.e.m.) at a given ligand density. **b**, Dose-response curves were fitted with Eq. 6 to extract activation thresholds. For each construct, data are from at least three independent experiments and three different mice. Error bars represent the standard error of the fits. **c**, Representative TIRF images for the different constructs (labeled with AF555) at ligand densities of ~50 µm$^{-2}$. The cell outline is indicated by a dashed white contour line. Images were recorded 10 min after seeding. Scale bar, 2µm.

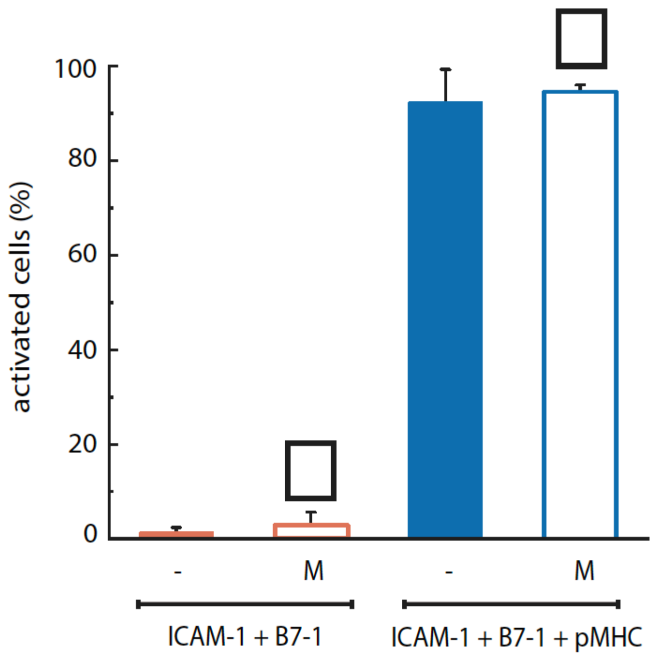

**Supplementary Fig. 14: Ligand-free DNA origami platforms do not influence T-cell activation behavior.** The presence of ligand-free medium-sized DNA origami platforms (M) at a density of ~50 $\mu m^{-2}$ did not alter the intracellular calcium response at either non-activating (ICAM-1 100 $\mu m^{-2}$, B7-1 100 $\mu m^{-2}$) or activating (ICAM-1 100 $\mu m^{-2}$, B7-1 100 $\mu m^{-2}$, His-pMHC 150 $\mu m^{-2}$) conditions. Data are the mean of at least two independent experiments (± s.e.m.).

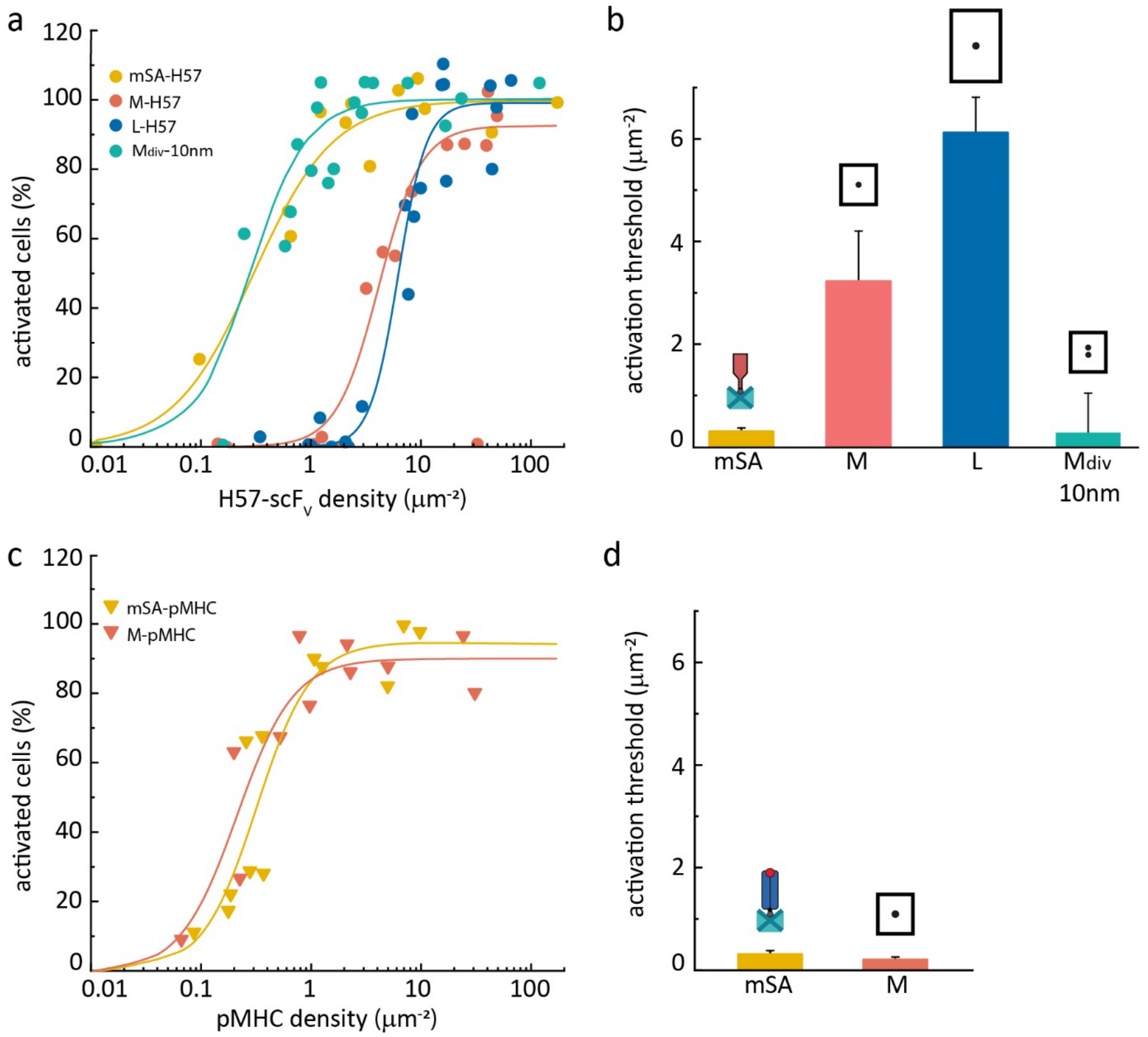

**Supplementary Fig. 15: T-cell activation on biointerfaces at 37°C.** The percentage of activated cells at different surface densities of H57-scFv **(a)** or pMHC **(c)** was measured at 37°C. Data were normalized to a positive control containing His-tagged ICAM-1 and B7-1 at 100 µm$^{-2}$, and His-pMHC at 150 µm$^{-2}$ (=100%). Each data point represents the population average of 187 ± 46 **(a)** and 170 ± 57 **(c)** cells (mean ± s.e.m.) at a given ligand density. For each construct, data are from at least two independent experiments and two different mice. **b, d,** Activation thresholds determined from fits of data from **(a)** and **(c)**. $t_{1/2}$ for the H57:TCR and pMHC:TCR interaction at 37°C are 6 min and 100 ms, respectively[6]. Error bars represent the standard error of the fits.

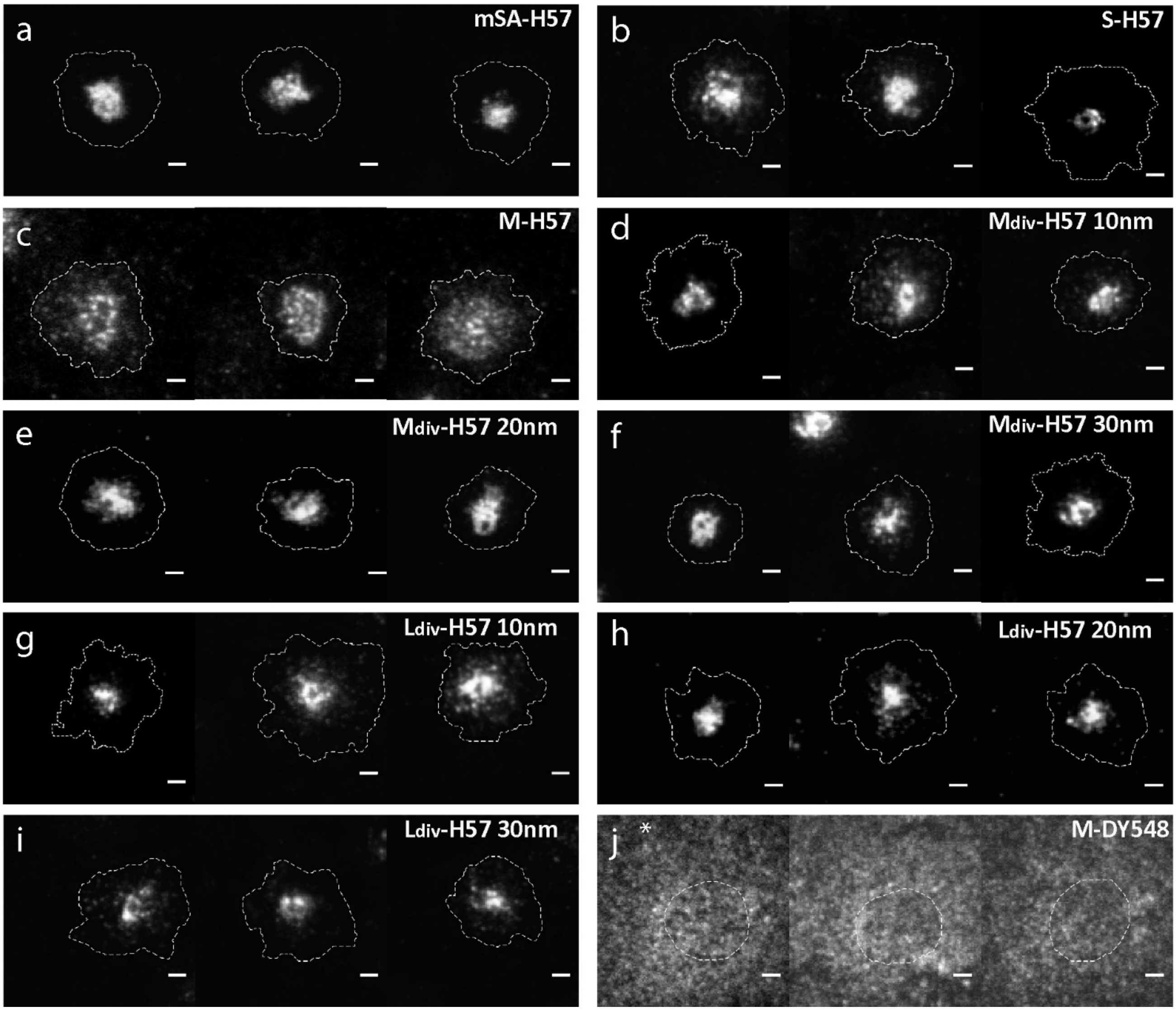

**Supplementary Fig. 16: H57-scF$_V$ organization at the T-cell-SLB interface.** Representative TIRF images showing the organization of AF555-labeled H57-scF$_V$ at densities of ~50 µm$^{-2}$ for the different constructs. Images were recorded 10 min after cell seeding. All constructs bearing H57-scF$_V$ (**a,** mSA-H57, **b,** S-H57, **c,** M-H57, **d,** M$_{div}$-H57$_{10nm}$, **e,** M$_{div}$-H57$_{20nm}$, **f,** M$_{div}$-H57$_{30nm}$, **g,** L$_{div}$-H57$_{10nm}$, **h,** L$_{div}$-H57$_{20nm}$, **i,** L$_{div}$-H57$_{30nm}$) showed characteristic microcluster or cSMAC formation. For ligand-free DNA origami platforms visualized by labeling with a Dy548-oligo (**j,** M-DY548) no microcluster formation could be observed. Scale bar, 2µm. *In order to visualize signals for M-DY548, image brightness was increased by a factor of 10.

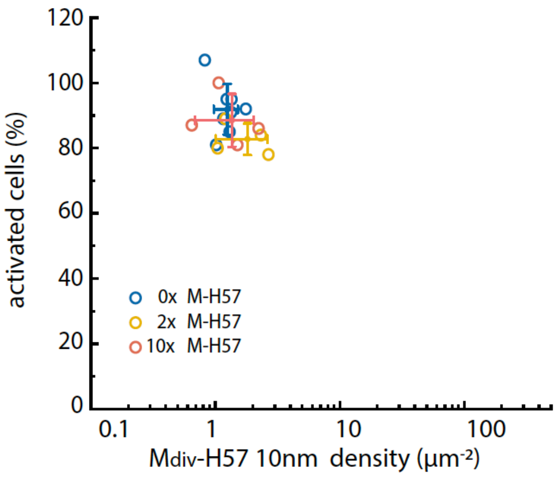

**Supplementary Fig. 17: On biointerfaces featuring $M_{div}$-H57$_{10nm}$, only DNA origami platforms functionalized with two scF$_V$s contribute to signaling.** A 2- or 10-fold molar excess of M-H57 was added at surface densities of $M_{div}$-H57$_{10nm}$ between ~0.5 and 2 µm$^{-2}$ and the percentage of activated cells was determined. Each data point represents the population average of 171 ± 28 cells (mean ± s.e.m.). For each condition, the mean ± s.d. is shown.

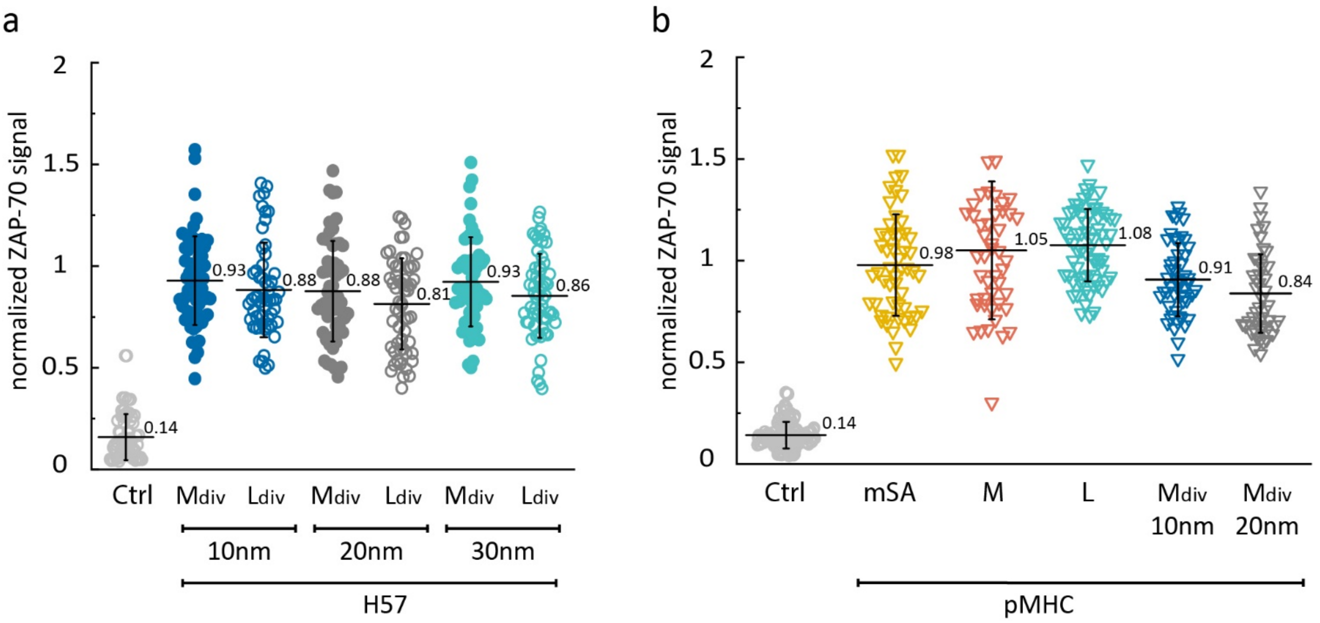

**Supplementary Fig. 18: ZAP-70 recruitment to the TCR.** Immunostaining of ZAP-70 recruited to the T-cell plasma membrane on biointerfaces featuring divalent H57-scF$_v$ DNA origami platforms **(a)** or pMHC constructs **(b)**. Ligand densities were ~50 μm$^{-2}$. T-cells were fixed 10 min after cell seeding and immunostained for ZAP-70. The signal was analyzed and normalized using a positive control containing His-tagged ICAM-1 and B7-1 at 100 μm$^{-2}$, and His-pMHC at 150 μm$^{-2}$ as a reference. Cells on bilayers containing only ICAM-1 and B7-1 at 100 μm$^{-2}$ are shown as a negative control (Ctrl). Data are shown as means ± s.d., each data point representing a single cell, n > 50 cells from at least two independent experiments and two different mice.

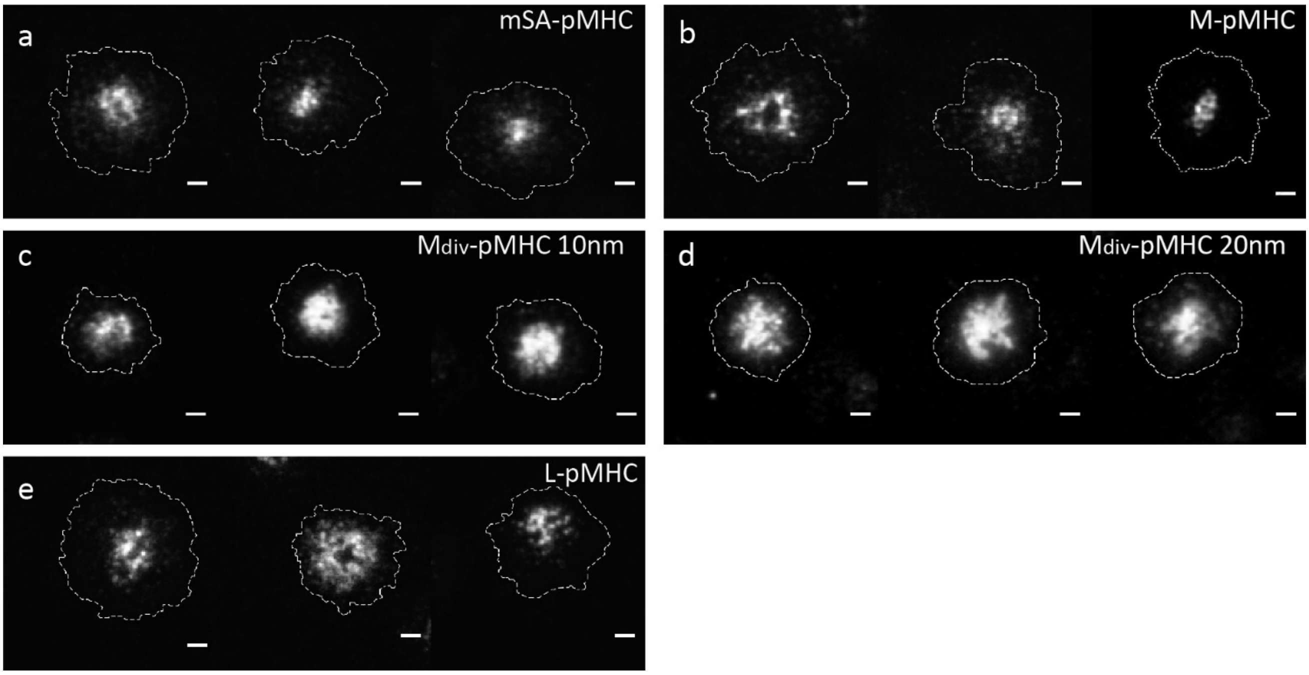

**Supplementary Fig. 19: pMHC organization at the T-cell-SLB interface.** Representative TIRF images showing the organization of pMHC (labeled with AF555) at densities of ~50 $\mu m^{-2}$ for the different constructs. Images were recorded 10 min after cell seeding. All constructs bearing pMHC (**a,** mSA-pMHC, **b,** M-pMHC, **c,** $M_{div}$-$pMHC_{10nm}$, **d,** $M_{div}$-$pMHC_{20nm}$, **e,** L-pMHC) showed characteristic microcluster or cSMAC formation. Scale bar, 2µm.

**Supplementary Table 1: Diffusion coefficients of H57-scF$_V$ constructs on SLBs.** Single-molecule trajectories of TCR ligand constructs were recorded on SLBs, pooled and diffusion coefficients were determined by mean square displacement analysis. DNA origami without modification were stained with YOYO. Diffusion coefficients are given as mean ± s.e.m.. Data are from at least two independent experiments.

| layout | ligand | D ($\mu m^2/s$) | trajectories (n) |
|---|---|---|---|
| **S** | - | 0.257 ± 0.046 | 19,615 |
| **M** | - | 0.273 ± 0.063 | 32,582 |
| **L** | - | 0.236 ± 0.040 | 46,441 |
| **mSA** | H57-scF$_V$ | 0.548 ± 0.021 | 10,070 |
| **DNA** | H57-scF$_V$ | 0.454 ± 0.009 | 2,218 |
| **mSA** | pMHC | 0.562 ± 0.018 | 5,582 |
| **S** | H57-scF$_V$ | 0.284 ± 0.057 | 20,474 |
| **M** | H57-scF$_V$ | 0.239 ± 0.010 | 10,047 |
| **L** | H57-scF$_V$ | 0.322 ± 0.028 | 6,561 |
| **M** | pMHC | 0.288 ± 0.011 | 8,482 |
| **L** | pMHC | 0.264 ± 0.010 | 2,420 |
| **$M_{div}$ 10nm** | H57-scF$_V$ | 0.238 ± 0.009 | 17,187 |
| **$M_{div}$ 20nm** | H57-scF$_V$ | 0.305 ± 0.008 | 6,090 |
| **$M_{div}$ 30nm** | H57-scF$_V$ | 0.233 ± 0.002 | 11,418 |
| **$M_{div}$ 10nm** | pMHC | 0.261 ± 0.023 | 9,959 |
| **$M_{div}$ 20nm** | pMHC | 0.256 ± 0.011 | 10,770 |
| **$L_{div}$ 10nm** | H57-scF$_V$ | 0.287 ± 0.017 | 22,094 |
| **$L_{div}$ 20nm** | H57-scF$_V$ | 0.261 ± 0.033 | 5,000 |
| **$L_{div}$ 30nm** | H57-scF$_V$ | 0.249 ± 0.045 | 9,746 |

**Supplementary Table 2: Ligand occupancies of DNA origami platforms.** Efficiency of functionalization with TCR ligands was determined via sequential two-color colocalization experiments as described in the Methods section. Data are the mean of at least two independent experiments (± s.e.m.).

| layout | ligand | no ligand (%) | single occupancy (%) | double occupancy (%) |
|---|---|---|---|---|
| **S** | H57-scF$_V$ | 58.7 ± 1.1 | 41.3 ± 1.1 | n.a. |
| **M** | H57-scF$_V$ | 40.0 ± 0.7 | 60.0 ± 0.7 | n.a. |
| **L** | H57-scF$_V$ | 39.0 ± 1.0 | 61.0 ± 1.0 | n.a. |
| **M** | pMHC | 39.8 ± 1.2 | 60.2 ± 1.2 | n.a. |
| **L** | pMHC | 39.5 ± 1.4 | 60.5 ± 1.4 | n.a. |
| **$M_{div}$ 10nm** | H57-scF$_V$ | 23.7 ± 1.3 | 42.7 ± 2.4 | 33.6 ± 2.4 |
| **$M_{div}$ 20nm** | H57-scF$_V$ | 19.4 ± 1.1 | 46.5 ± 0.9 | 34.1 ± 0.9 |
| **$M_{div}$ 30nm** | H57-scF$_V$ | 16.3 ± 1.8 | 46.5 ± 3.5 | 37.2 ± 3.5 |
| **$M_{div}$ 10nm** | pMHC | 28.6 ± 1.5 | 38.4 ± 4.7 | 33.0 ± 4.7 |
| **$M_{div}$ 20nm** | pMHC | 33.7 ± 1.0 | 28.8 ± 0.6 | 37.5 ± 0.6 |
| **$L_{div}$ 10nm** | H57-scF$_V$ | 13.9 ± 1.4 | 48.3 ± 2.2 | 37.8 ± 2.2 |
| **$L_{div}$ 20nm** | H57-scF$_V$ | 28.7 ± 2.9 | 37.9 ± 2.3 | 33.4 ± 2.3 |
| **$L_{div}$ 30nm** | H57-scF$_V$ | 12.7 ± 2.6 | 44.4 ± 2.6 | 42.9 ± 2.6 |

**Supplementary Table 3: Fitting parameters of fits to dose-response curves.** Dose-response curves recorded for the different constructs featuring one (mSA, DNA, S, M, L) or two ($M_{div}$, $L_{div}$) TCR ligands (H57-scFvs, pMHCs) were fitted with Eq. 6 to extract the activation threshold $T_A$, the maximum response $A_{max}$ and the Hill coefficient n. Standard errors of the fits are shown. *For fitting of data recorded for M-H57 at 24°C, $A_{max}$ was fixed at 100%.

| layout | ligand | modification | T (°C) | n | $A_{max}$ (%) | $T_A$ ($\mu m^{-2}$) |
|---|---|---|---|---|---|---|
| **mSA** | H57-scFv | - | 24 | 2.47 ± 1.11 | 103.45 ± 3.71 | 1.35 ± 0.31 |
| **DNA** | H57-scFv | - | 24 | 2.96 ± 0.70 | 97.50 ± 3.64 | 1.57 ± 0.11 |
| **S** | H57-scFv | - | 24 | 1.19 ± 0.42 | 103.18 ± 10.36 | 0.92 ± 0.28 |
| **M** | H57-scFv | - | 24 | 2.31 ± 1.36 | 100.00 ± 0.00* | 272.17 ± 23.02 |
| **$M_{div}$ 10nm** | H57-scFv | - | 24 | 2.33 ± 0.32 | 102.38 ± 2.95 | 1.0 ± 0.06 |
| **$M_{div}$ 20nm** | H57-scFv | - | 24 | 2.65 ± 0.49 | 102.99 ± 2.83 | 2.16 ± 0.27 |
| **$M_{div}$ 30nm** | H57-scFv | - | 24 | 4.19 ± 1.87 | 108.68 ± 12.50 | 18.82 ± 2.40 |
| **$L_{div}$ 10nm** | H57-scFv | - | 24 | 1.23 ± 0.37 | 101.99 ± 7.55 | 1.67 ± 0.43 |
| **$L_{div}$ 20nm** | H57-scFv | - | 24 | 5.55 ± 3.00 | 91.24 ± 5.82 | 2.55 ± 0.27 |
| **$L_{div}$ 30nm** | H57-scFv | - | 24 | 6.96± 2.39 | 96.04 ± 4.32 | 16.46 ± 1.20 |
| **mSA** | pMHC | - | 24 | 2.48 ± 0.61 | 95.07 ± 3.05 | 0.73 ± 0.08 |
| **M** | pMHC | - | 24 | 2.90 ± 0.53 | 91.55 ± 3.10 | 0.78 ± 0.07 |
| **L** | pMHC | - | 24 | 2.57 ± 0.53 | 105.62 ± 4.98 | 0.91 ± 0.07 |
| **$M_{div}$ 10nm** | pMHC | - | 24 | 1.65 ± 0.16 | 100.46 ± 2.65 | 2.11 ± 0.15 |
| **$M_{div}$ 20nm** | pMHC | - | 24 | 6.18 ± 1.51 | 95.89 ± 2.39 | 1.14 ± 0.04 |
| **mSA** | H57-scFv | - | 37 | 1.20 ± 0.28 | 98.46 ± 4.12 | 0.30 ± 0.07 |
| **M** | H57-scFv | - | 37 | 2.26 ± 0.45 | 91.27 ± 3.77 | 4.07 ± 0.38 |
| **L** | H57-scFv | - | 37 | 3.39 ± 1.25 | 97.75 ± 5.05 | 6.12 ± 0.69 |
| **$M_{div}$ 10nm** | H57-scFv | - | 37 | 1.61 ± 0.43 | 101.63± 5.31 | 0.36 ± 0.07 |
| **mSA** | pMHC | - | 37 | 1.76 ± 0.75 | 94.93 ± 8.88 | 0.32 ± 0.07 |
| **M** | pMHC | - | 37 | 1.69 ± 0.72 | 90.16 ± 6.37 | 0.21 ± 0.05 |
| **mSA** | H57-scFv | CD4-blocked | 24 | 8.03 ± 3.98 | 105.46 ± 2.94 | 2.13 ± 0.33 |
| **mSA** | pMHC | CD4-blocked | 24 | 7.28 ± 1.37 | 106.76 ± 3.34 | 33.38 ± 0.88 |
| **M** | pMHC | CD4-blocked | 24 | 2.02 ± 1.20 | 118.61 ± 50.43 | 41.70 ± 22.43 |

**Supplementary Table 4: Minimum distances of ligands as permitted by DNA origami platforms.** DNA origami platforms act as spacers that limit the approach of ligands to a minimum distance **δ**, representing the smallest possible distance between two individual ligands. Note that for divalent DNA origami featuring two ligands, **δ** is a fixed value. Here, **d** is the smallest possible distance between ligands on adjacent DNA origami platforms. Quasi-crystalline packing of DNA origami was assumed for all cases.

| layout | δ (nm) | d (nm) |
|---|---|---|
| **S** | 20 | n.a. |
| **M** | 48 | n.a. |
| **L** | 60 | n.a. |
| **$M_{div}$ 10nm** | 10 | 38 |
| **$M_{div}$ 20nm** | 20 | 38 |
| **$M_{div}$ 30nm** | 30 | 40 |
| **$L_{div}$ 10nm** | 10 | 60 |
| **$L_{div}$ 20nm** | 20 | 62 |
| **$L_{div}$ 30nm** | 30 | 40 |

### Supplementary Table 5: List of staple strands

| DNA origami layout [nm] | Name | Sequence |
|---|---|---|
| 30x20 | 7[56]-BLK | GAAAAACCAAGGGCGATCGCGAGATAGGGTTG |
| 30x20 | 7[80]-BLK | CAGCCAGCTTTCCGGCCCATTCGCCATTCAGGCTGGCGAA |
| 30x20 | 3[58]-BLK | GCATGCTCCACACAACAGAGACGGGCAACA |
| 30x20 | 1[40]-BLK | TCTTTTCACCAGTTACGAGCCGGATGCCAGCT |
| 30x20 | 3[72]-BLK | CGACTCTAGACGTTGTAAAACGACTTCGCTAT |
| 30x20 | 0[55]-BLK | GCATTAATGAATCGGCCAACGCGCGTGGTTTT |
| 30x20 | 0[79]-BLK | GTCGGGAAACCTGTCGAGCATAAA |
| 30x20 | 5[58]-BLK | GGCCTCGGCCAGTGCCACCCCAGCAGGCGA |
| 30x20 | 5[72]-BLK | TACGCCAGCTGCGCAACTGTTGGGGTCTATCATCGCACTC |
| 65x54 | 10[63]-BLK | CAGCTTTCGGGCGCATCGTAACCGATCGGCCT |
| 65x54 | 11[112]-BLK | AGCGCGAACAATCATAAGGGAACCCGGTGTAC |
| 65x54 | 12[95]-BLK | ACACTCATAGGGGACGACGACAGTTGCATCTG |
| 65x54 | 13[80]-BLK | CGCCAGCTAAAGACAGCATCGGAAGTCACCCT |
| 65x54 | 14[63]-BLK | GCGATTAAGTACCGAGCTCGAATTAATTGTTA |
| 65x54 | 15[112]-BLK | TTTCTTAACCGCTTTTGCGGGATCCGAGGGTA |
| 65x54 | 16[95]-BLK | GTATCGGTGCTGTTTCCTGTGTGACGTAATCA |
| 65x54 | 19[112]-BLK | TAGCAAGCCGATCTAAAGTTTTGTGTATGGGA |
| 65x54 | 2[63]-BLK | TATTTTCATTAAGCAATAAAGCCTACATTATG |
| 65x54 | 5[80]-BLK | CGGAGACATTCATCAGTTGAGATTATTACAGG |
| 65x54 | 6[63]-BLK | TCAACCGTATCGATGAACGGTAATGGTTGATA |
| 65x54 | 7[112]-BLK | TAATTTCACTAACGGAACAACATTTAGGAATA |
| 65x54 | 8[95]-BLK | AGAACGAGCAATCATATGTACCCCCGTAAAAC |
| 65x54 | 9[80]-BLK | CAAAAATAAAAGAGGACAGATGAAGAACTGAC |
| 65x54 | 1[32]-BLK | TGATTCCCAAAAGGTGGCATCAATAATCATAC |
| 65x54 | 17[32]-BLK | CATTAATTGGGCGCCAGGGTGGTTATTGCCCT |
| 65x54 | 20[143]-BLK | AGGAGGTTTACCGTAACACTGAGTCATTCCAC |
| 65x54 | 4[143]-BLK | TACCAGACCGGAATCGTCATAAATGTTCAGAA |
| 65x54 | 0[143]-BLK | ACAGGTCAGGATTAGAGAGTACCTGAAGCCCG |
| 65x54 | 0[47]-BLK | GCTCAACATGTTTTAAATATGCAAATAACAGT |
| 65x54 | 0[63]-BLK | TGAATATAAGATTTAGTTTGACCATTTAGCTA |
| 65x54 | 0[95]-BLK | TCATTTTTGCGGATGGCTTAGAGCTTAATTGC |
| 65x54 | 1[112]-BLK | ATTAAGAGTTAATTGCTCCTTTTGATAAGAGG |
| 65x54 | 1[128]-BLK | AAAGACTTTGACCATAAATCAAAAATAGCGTC |
| 65x54 | 1[32]-BLK | TGATTCCCAAAAGGTGGCATCAATAATCATAC |
| 65x54 | 1[80]-BLK | ATTTCGCAAGCAAAGCGGATTGCAGACTATTA |
| 65x54 | 10[143]-BLK | TAGCCGGACCTTCATCAAGAGTAATCAACGTA |
| 65x54 | 10[95]-BLK | CAACTTTGATTCGCGTCTGGCCTTAACGCCAT |
| 65x54 | 11[32]-BLK | GGGATAGGTTTCCGGCACCGCTTCCATTCAGG |
| 65x54 | 11[80]-BLK | CCAGTTTGCTTTGACCCCCAGCGAACACTAAA |
| 65x54 | 12[143]-BLK | CTACGAAGATTTGTATCATCGCCTATGTTACT |
| 65x54 | 12[47]-BLK | CAGCCAGCTCACGTTGGTGTAGATATCAACAT |
| 65x54 | 12[63]-BLK | CAGGAAGATCGGTGCGGGCCTCTTGCTGCAAG |
| 65x54 | 13[112]-BLK | GCAACGGCAAAGAGGCAAAAGAATTTATACCA |
| 65x54 | 13[128]-BLK | CTTTGAGGTGCAGGGAGTTAAAGGACAGCTTG |
| 65x54 | 13[32]-BLK | CTGCGCAAGTTTTCCCAGTCACGAATGCCTGC |
| 65x54 | 14[143]-BLK | CTGAGGCTACTAAAGACTTTTTCATAATGCCA |
| 65x54 | 14[95]-BLK | CAGCAGCGGGCGAAAGGGGGATGTCGCTATTA |
| 65x54 | 15[32]-BLK | AGGTCGACACGAGCCGGAAGCATACTAACTCA |
| 65x54 | 15[80]-BLK | TGGTCATATTATCAGCTTGCTTTCCTTTAATT |

| | | |
|---|---|---|
| 65x54 | 16[143]-BLK | TCACGTTGAGTTGCGCCGACAATGTTCGGTCG |
| 65x54 | 16[47]-BLK | CACAACATTCTAGAGGATCCCCGGGTTGGGTA |
| 65x54 | 16[63]-BLK | TCCGCTCACCGCTTTCCAGTCGGGGCCAACGC |
| 65x54 | 17[112]-BLK | TTTTGCTAAGGCTCCAAAAGGAGCGAGGTGAA |
| 65x54 | 17[128]-BLK | TCAACAGTCTCATAGTTAGCGTAACCAATAGG |
| 65x54 | 17[32]-BLK | CATTAATTGGGCGCCAGGGTGGTTATTGCCCT |
| 65x54 | 17[80]-BLK | CGTGCCAGAGTAAATGAATTTTCTCGTCTTTC |
| 65x54 | 18[143]-BLK | AGACAGCCTTCAGCGGAGTGAGAATAATTTTT |
| 65x54 | 18[63]-BLK | GCGGGGAGGCAGCAAGCGGTCCACTGATGGTG |
| 65x54 | 18[95]-BLK | CAGACGTTCTGCATTAATGAATCGAAACCTGT |
| 65x54 | 19[32]-BLK | TCACCGCCATAAATCAAAAGAATAGGAACAAG |
| 65x54 | 19[80]-BLK | GCCCCAGCAGCCACCACCCTCATTGAACCGCC |
| 65x54 | 2[143]-BLK | AACGAGAACAAATATCGCGTTTTAAAACTCCA |
| 65x54 | 2[95]-BLK | TAGTCAGAAATGGTCAATAACCTGTTAGATAC |
| 65x54 | 20[143]-BLK | AGGAGGTTTACCGTAACACTGAGTCATTCCAC |
| 65x54 | 20[47]-BLK | AATCCCTTTGGCCCTGAGAGAGTTAGGCGGTT |
| 65x54 | 20[63]-BLK | GTTCCGAATCCAACGTCAAAGGGCGAAAAACC |
| 65x54 | 20[95]-BLK | ACCCTCAGAGGCGAAAATCCTGTTGCTGGTTT |
| 65x54 | 21[112]-BLK | TTTGCTCAGAACCGCCACCCTCATTCAGGGA |
| 65x54 | 21[128]-BLK | GCGGATAAGTGCCGTCGAGAGGGTCCGTACTC |
| 65x54 | 21[32]-BLK | AGTCCACTATTAAAGAACGTGGACATCGGCAA |
| 65x54 | 21[80]-BLK | GTCTATCAAGAGAAGGATTAGGATTAGCGGGG |
| 65x54 | 3[112]-BLK | TAGACTGGATCAGGTCTTTACCCTTCAAAAAG |
| 65x54 | 3[32]-BLK | AGGCAAGGAGCCTTTATTTCAACGTTTAAATG |
| 65x54 | 3[80]-BLK | AAAGCTAACAGAGGGGGTAATAGTGCAAAAGA |
| 65x54 | 4[143]-BLK | TACCAGACCGGAATCGTCATAAATGTTCAGAA |
| 65x54 | 4[47]-BLK | GCGGGAGACAAAGAATTAGCAAAATTTGGGGC |
| 65x54 | 4[63]-BLK | ACCCTGTAAAGATTCAAAAGGGTGTATGATAT |
| 65x54 | 4[95]-BLK | AGTTTTGCATCGGTTGTACCAAAACAGAGCAT |
| 65x54 | 5[112]-BLK | CCACATTCATAGCGAGAGGCTTTTAAAATGTT |
| 65x54 | 5[128]-BLK | CAGATACAACGTTAATAAAACGAAACTTTAAT |
| 65x54 | 5[32]-BLK | CAATGCCTATGCCGGAGAGGGTAGTCATTGCC |
| 65x54 | 6[143-BLK] | AAAAATCTTAACGCCAAAAGGAATCCCTCGTT |
| 65x54 | 6[95]-BLK | TAGAAAGAGTCAAATCACCATCAAAGAAAGGC |
| 65x54 | 7[32]-BLK | TGAGAGTCAGATTGTATAAGCAAAAATTCGCA |
| 65x54 | 7[80]-BLK | TAGCATGTTAGTAAATTGGGCTTGGAAACACC |
| 65x54 | 8[143-BLK] | ACAAAGCTATTACCTTATGCGATTTTGGGAAG |
| 65x54 | 8[47]-BLK | AAACAGGATGGAGCAAACAAGAGATCTAGCTG |
| 65x54 | 8[63]-BLK | ATCAGAAATTTTTTAACCAATAGGCCTGTAGC |
| 65x54 | 9[112-BLK] | AGACCAGGAGGCTTGCCCTGACGAAGATGGTT |
| 65x54 | 9[128-BLK] | CTGGCTGAACGAGGCGCAGACGGTACAAAGTA |
| 65x54 | 9[32]-BLK | TTAAATTTAGCGAGTAACAACCCGACCGTAAT |
| 100x70 | 0[111]-BLK | TAAATGAATTTTCTGTATGGGATTAATTTCTT |
| 100x70 | 0[143]-BLK | TCTAAAGTTTTGTCGTCTTTCCAGCCGACAA |
| 100x70 | 0[175]-BLK | TCCACAGACAGCCCTCATAGTTAGCGTAACGA |
| 100x70 | 0[207]-BLK | TCACCAGTACAAACTACAACGCCTAGTACCAG |
| 100x70 | 0[239]-BLK | AGGAACCCATGTACCGTAACACTTGATATAA |
| 100x70 | 0[271]-BLK | CCACCCTCATTTTCAGGGATAGCAACCGTACT |
| 100x70 | 0[47]-BLK | AGAAAGGAACAACTAAAGGAATTCAAAAAAA |
| 100x70 | 0[79]-BLK | ACAACTTTCAACAGTTTCAGCGGATGTATCGG |

| | | |
|---|---|---|
| 100x70 | 1[128]-BLK | TGACAACTCGCTGAGGCTTGCATTATACCAAGCGCGATGATAAA |
| 100x70 | 1[160]-BLK | TTAGGATTGGCTGAGACTCCTCAATAACCGAT |
| 100x70 | 1[192]-BLK | GCGGATAACCTATTATTCTGAAACAGACGATTGGCCTTGAAGAGCCAC |
| 100x70 | 1[224]-BLK | GTATAGCAAACAGTTAATGCCCAATCCTCA |
| 100x70 | 1[256]-BLK | CAGGAGGTGGGGTCAGTGCCTTGAGTCTCTGAATTTACCGGGAACCAG |
| 100x70 | 1[32]-BLK | AGGCTCCAGAGGCTTTGAGGACACGGGTAA |
| 100x70 | 1[64]-BLK | TTTATCAGGACAGCATCGGAACGACACCAACCTAAAACGAGGTCAATC |
| 100x70 | 1[96]-BLK | AAACAGCTTTTTGCGGGATCGTCAACACTAAA |
| 100x70 | 10[111]-BLK | TTGCTCCTTTCAAATATCGCGTTTGAGGGGGT |
| 100x70 | 10[143]-BLK | CCAACAGGAGCGAACCAGACCGGAGCCTTTAC |
| 100x70 | 10[175]-BLK | TTAACGTCTAACATAAAAACAGGTAACGGA |
| 100x70 | 10[207]-BLK | ATCCCAATGAGAATTAACTGAACAGTTACCAG |
| 100x70 | 10[239]-BLK | GCCAGTTAGAGGGTAATTGAGCGCTTTAAGAA |
| 100x70 | 10[271]-BLK | ACGCTAACACCCACAAGAATTGAAAATAGC |
| 100x70 | 10[47]-BLK | CTGTAGCTTGACTATTATAGTCAGTTCATTGA |
| 100x70 | 10[79]-BLK | GATGGCTTATCAAAAAGATTAAGAGCGTCC |
| 100x70 | 11[128]-BLK | TTTGGGGATAGTAGTAGCATTAAAAGGCCG |
| 100x70 | 11[160]-BLK | CCAATAGCTCATCGTAGGAATCATGGCATCAA |
| 100x70 | 11[192]-BLK | TATCCGGTCTCATCGAGAACAAGCGACAAAAG |
| 100x70 | 11[224]-BLK | GCGAACCTCCAAGAACGGGTATGACAATAA |
| 100x70 | 11[256]-BLK | GCCTTAAACCAATCAATAATCGGCACGCGCCT |
| 100x70 | 11[32]-BLK | AACAGTTTTGTACCAAAAACATTTTATTTC |
| 100x70 | 11[64]-BLK | GATTTAGTCAATAAAGCCTCAGAGAACCCTCA |
| 100x70 | 11[96]-BLK | AATGGTCAACAGGCAAGGCAAAGAGTAATGTG |
| 100x70 | 12[111]-BLK | TAAATCATATAACCTGTTTAGCTAACCTTTAA |
| 100x70 | 12[143]-BLK | TTCTACTACGCGAGCTGAAAAGGTTACCGCGC |
| 100x70 | 12[175]-BLK | TTTTATTTAAGCAAATCAGATATTTTTTGT |
| 100x70 | 12[207]-BLK | GTACCGCAATTCTAAGAACGCGAGTATTATTT |
| 100x70 | 12[239]-BLK | CTTATCATTCCCGACTTGCGGGAGCCTAATTT |
| 100x70 | 12[271]-BLK | TGTAGAAATCAAGATTAGTTGCTCTTACCA |
| 100x70 | 12[47]-BLK | TAAATCGGGATTCCCAATTCTGCGATATAATG |
| 100x70 | 12[79]-BLK | AAATTAAGTTGACCATTAGATACTTTTGCG |
| 100x70 | 13[128]-BLK | GAGACAGCTAGCTGATAAATTAATTTTTGT |
| 100x70 | 13[192]-BLK | GTAAAGTAATCGCCATATTTAACAAAACTTTT |
| 100x70 | 13[224]-BLK | ACAACATGCCAACGCTCAACAGTCTTCTGA |
| 100x70 | 13[256]-BLK | GTTTATCAATATGCGTTATACAAACCGACCGT |
| 100x70 | 13[32]-BLK | AACGCAAAATCGATGAACGGTACCGGTTGA |
| 100x70 | 13[64]-BLK | TATATTTTGTCATTGCCTGAGAGTGGAAGATT |
| 100x70 | 13[96]-BLK | TAGGTAAACTATTTTTGAGAGATCAAACGTTA |
| 100x70 | 14[111]-BLK | GAGGGTAGGATTCAAAAGGGTGAGACATCCAA |
| 100x70 | 14[143]-BLK | CAACCGTTTCAAATCACCATCAATTCGAGCCA |
| 100x70 | 14[175]-BLK | CATGTAATAGAATATAAAGTACCAAGCCGT |
| 100x70 | 14[207]-BLK | AATTGAGAATTCTGTCCAGACGACTAAACCAA |
| 100x70 | 14[239]-BLK | AGTATAAAGTTCAGCTAATGCAGATGTCTTTC |
| 100x70 | 14[271]-BLK | TTAGTATCACAATAGATAAGTCCACGAGCA |
| 100x70 | 14[47]-BLK | AACAAGAGGGATAAAAATTTTTAGCATAAAGC |
| 100x70 | 14[79]-BLK | GCTATCAGAAATGCAATGCCTGAATTAGCA |
| 100x70 | 15[128]-BLK | TAAATCAAAATAATTCGCGTCTCGGAAACCAGGCAAAGGGAAGG |
| 100x70 | 15[160]-BLK | ATCGCAAGTATGTAAATGCTGATGATAGGAAC |
| 100x70 | 15[192]-BLK | TCAAATATAACCTCCGGCTTAGGTAACAATTTCATTTGAAGGCGAATT |

| | | |
|---|---|---|
| 100x70 | 15[224]-BLK | CCTAAATCAAAATCATAGGTCTAAACAGTA |
| 100x70 | 15[256]-BLK | GTGATAAAAAGACGCTGAGAAGAGATAACCTTGCTTCTGTTCGGGAGA |
| 100x70 | 15[32]-BLK | TAATCAGCGGATTGACCGTAATCGTAACCG |
| 100x70 | 15[64]-BLK | GTATAAGCCAACCCGTCGGATTCTGACGACAGTATCGGCCGCAAGGCG |
| 100x70 | 15[96]-BLK | ATATTTTGGCTTTCATCAACATTATCCAGCCA |
| 100x70 | 16[111]-BLK | TGTAGCCATTAAAATTCGCATTAAATGCCGGA |
| 100x70 | 16[143]-BLK | GCCATCAAGCTCATTTTTTAACCACAAATCCA |
| 100x70 | 16[175]-BLK | TATAACTAACAAAGAACGCGAGAACGCCAA |
| 100x70 | 16[207]-BLK | ACCTTTTTATTTTAGTTAATTTCATAGGGCTT |
| 100x70 | 16[239]-BLK | GAATTTATTTAATGGTTTGAAATATTCTTACC |
| 100x70 | 16[271]-BLK | CTTAGATTTAAGGCGTTAAATAAAGCCTGT |
| 100x70 | 16[47]-BLK | ACAAACGGAAAAGCCCCAAAAACACTGGAGCA |
| 100x70 | 16[79]-BLK | GCGAGTAAAAATATTTAAATTGTTACAAAG |
| 100x70 | 17[224]-BLK | CATAAATCTTTGAATACCAAGTGTTAGAAC |
| 100x70 | 17[32]-BLK | TGCATCTTTCCCAGTCACGACGGCCTGCAG |
| 100x70 | 17[96]-BLK | GCTTTCCGATTACGCCAGCTGGCGGCTGTTTC |
| 100x70 | 18[111]-BLK | TCTTCGCTGCACCGCTTCTGGTGCGGCCTTCC |
| 100x70 | 18[143]-BLK | CAACTGTTGCGCCATTCGCCATTCAAACATCA |
| 100x70 | 18[175]-BLK | CTGAGCAAAAATTAATTACATTTTGGGTTA |
| 100x70 | 18[207]-BLK | CGCGCAGATTACCTTTTTTAATGGGAGAGACT |
| 100x70 | 18[239]-BLK | CCTGATTGCAATATATGTGAGTGATCAATAGT |
| 100x70 | 18[271]-BLK | CTTTTACAAAATCGTCGCTATTAGCGATAG |
| 100x70 | 18[47]-BLK | CCAGGGTTGCCAGTTTGAGGGGACCCGTGGGA |
| 100x70 | 18[79]-BLK | GATGTGCTTCAGGAAGATCGCACAATGTGA |
| 100x70 | 19[160]-BLK | GCAATTCACATATTCCTGATTATCAAAGTGTA |
| 100x70 | 19[224]-BLK | CTACCATAGTTTGAGTAACATTTAAAATAT |
| 100x70 | 19[32]-BLK | GTCGACTTCGGCCAACGCGCGGGGTTTTTC |
| 100x70 | 19[96]-BLK | CTGTGTGATTGCGTTGCGCTCACTAGAGTTGC |
| 100x70 | 2[111]-BLK | AAGGCCGCTGATACCGATAGTTGCGACGTTAG |
| 100x70 | 2[143]-BLK | ATATTCGGAACCATCGCCCACGCAGAGAAGGA |
| 100x70 | 2[175]-BLK | TATTAAGAAGCGGGGTTTTGCTCGTAGCAT |
| 100x70 | 2[207]-BLK | TTTCGGAAGTGCCGTCGAGAGGGTGAGTTTCG |
| 100x70 | 2[239]-BLK | GCCCGTATCCGGAATAGGTGTATCAGCCCAAT |
| 100x70 | 2[271]-BLK | GTTTTAACTTAGTACCGCCACCCAGAGCCA |
| 100x70 | 2[47]-BLK | ACGGCTACAAAAGGAGCCTTTAATGTGAGAAT |
| 100x70 | 2[79]-BLK | CAGCGAAACTTGCTTTCGAGGTGTTGCTAA |
| 100x70 | 20[111]-BLK | CACATTAAAATTGTTATCCGCTCATGCGGGCC |
| 100x70 | 20[143]-BLK | AAGCCTGGTACGAGCCGGAAGCATAGATGATG |
| 100x70 | 20[175]-BLK | ATTATCATTCAATATAATCCTGACAATTAC |
| 100x70 | 20[207]-BLK | GCGGAACATCTGAATAATGGAAGGTACAAAAT |
| 100x70 | 20[239]-BLK | ATTTTAAAATCAAAATTATTTGCACGGATTCG |
| 100x70 | 20[271]-BLK | CTCGTATTAGAAATTGCGTAGATACAGTAC |
| 100x70 | 20[47]-BLK | TTAATGAACTAGAGGATCCCCGGGGGGTAACG |
| 100x70 | 20[79]-BLK | TTCCAGTCGTAATCATGGTCATAAAAGGGG |
| 100x70 | 21[120]-BLK | CCCAGCAGGCGAAAAATCCCTTATAAATCAAGCCGGCG |
| 100x70 | 21[160]-BLK | TCAATATCGAACCTCAAATATCAATTCCGAAA |
| 100x70 | 21[184]-BLK | TCAACAGTTGAAAGGAGCAAATGAAAAATCTAGAGATAGA |
| 100x70 | 21[224]-BLK | CTTTAGGGCCTGCAACAGTGCCAATACGTG |
| 100x70 | 21[248]-BLK | AGATTAGAGCCGTCAAAAAACAGAGGTGAGGCCTATTAGT |
| 100x70 | 21[32]-BLK | TTTTCACTCAAAGGGCGAAAAACCATCACC |

| | | |
|---|---|---|
| 100x70 | 21[56]-BLK | AGCTGATTGCCCTTCAGAGTCCACTATTAAAGGGTGCCGT |
| 100x70 | 21[96]-BLK | AGCAAGCGTAGGGTTGAGTGTTGTAGGGAGCC |
| 100x70 | 22[111]-BLK | GCCCGAGAGTCCACGCTGGTTTGCAGCTAACT |
| 100x70 | 22[143]-BLK | TCGGCAAATCCTGTTTGATGGTGGACCCTCAA |
| 100x70 | 22[175]-BLK | ACCTTGCTTGGTCAGTTGGCAAAGAGCGGA |
| 100x70 | 22[207]-BLK | AGCCAGCAATTGAGGAAGGTTATCATCATTTT |
| 100x70 | 22[239]-BLK | TTAACACCAGCACTAACAACTAATCGTTATTA |
| 100x70 | 22[271]-BLK | CAGAAGATTAGATAATACATTTGTCGACAA |
| 100x70 | 22[47]-BLK | CTCCAACGCAGTGAGACGGGCAACCAGCTGCA |
| 100x70 | 22[79]-BLK | TGGAACAACCGCCTGGCCCTGAGGCCCGCT |
| 100x70 | 23[128]-BLK | AACGTGGCGAGAAAGGAAGGGAAACCAGTAA |
| 100x70 | 23[160]-BLK | TAAAAGGGACATTCTGGCCAACAAAGCATC |
| 100x70 | 23[192]-BLK | ACCCTTCTGACCTGAAAGCGTAAGACGCTGAG |
| 100x70 | 23[224]-BLK | GCACAGACAATATTTTTGAATGGGGTCAGTA |
| 100x70 | 23[256]-BLK | CTTTAATGCGCGAACTGATAGCCCCACCAG |
| 100x70 | 23[32]-BLK | CAAATCAAGTTTTTTGGGGTCGAAACGTGGA |
| 100x70 | 23[64]-BLK | AAAGCACTAAATCGGAACCCTAATCCAGTT |
| 100x70 | 23[96]-BLK | CCCGATTTAGAGCTTGACGGGGAAAAAGAATA |
| 100x70 | 3[160]-BLK | TTGACAGGCCACCACCAGAGCCGCGATTTGTA |
| 100x70 | 3[224]-BLK | TTAAAGCCAGAGCCGCCACCCTCGACAGAA |
| 100x70 | 3[32]-BLK | AATACGTTTGAAAGAGGACAGACTGACCTT |
| 100x70 | 3[96]-BLK | ACACTCATCCATGTTACTTAGCCGAAAGCTGC |
| 100x70 | 4[111]-BLK | GACCTGCTCTTTGACCCCCAGCGAGGGAGTTA |
| 100x70 | 4[143]-BLK | TCATCGCCAACAAAGTACAACGGACGCCAGCA |
| 100x70 | 4[175]-BLK | CACCAGAAAGGTTGAGGCAGGTCATGAAAG |
| 100x70 | 4[207]-BLK | CCACCCTCTATTCACAAACAAATACCTGCCTA |
| 100x70 | 4[239]-BLK | GCCTCCCTCAGAATGGAAAGCGCAGTAACAGT |
| 100x70 | 4[271]-BLK | AAATCACCTTCCAGTAAGCGTCAGTAATAA |
| 100x70 | 4[47]-BLK | GACCAACTAATGCCACTACGAAGGGGGTAGCA |
| 100x70 | 4[79]-BLK | GCGCAGACAAGAGGCAAAAGAATCCCTCAG |
| 100x70 | 5[224]-BLK | TCAAGTTTCATTAAAGGTGAATATAAAAGA |
| 100x70 | 5[32]-BLK | CATCAAGTAAAACGAACTAACGAGTTGAGA |
| 100x70 | 5[96]-BLK | TCATTCAGATGCGATTTTAAGAACAGGCATAG |
| 100x70 | 6[111]-BLK | ATTACCTTTGAATAAGGCTTGCCCAAATCCGC |
| 100x70 | 6[143]-BLK | GATGGTTTGAACGAGTAGTAAATTTACCATTA |
| 100x70 | 6[175]-BLK | CAGCAAAAGGAAACGTCACCAATGAGCCGC |
| 100x70 | 6[207]-BLK | TCACCGACGCACCGTAATCAGTAGCAGAACCG |
| 100x70 | 6[239]-BLK | GAAATTATTGCCTTTAGCGTCAGACCGGAACC |
| 100x70 | 6[271]-BLK | ACCGATTGTCGGCATTTTCGGTCATAATCA |
| 100x70 | 6[47]-BLK | TACGTTAAAGTAATCTTGACAAGAACCGAACT |
| 100x70 | 6[79]-BLK | TTATACCACCAAATCAACGTAACGAACGAG |
| 100x70 | 7[120]-BLK | CGTTTACCAGACGACAAAGAAGTTTTGCCATAATTCGA |
| 100x70 | 7[160]-BLK | TTATTACGAAGAACTGGCATGATTGCGAGAGG |
| 100x70 | 7[184]-BLK | CGTAGAAAATACATACCGAGGAAACGCAATAAGAAGCGCA |
| 100x70 | 7[224]-BLK | AACGCAAAGATAGCCGAACAAACCCTGAAC |
| 100x70 | 7[248]-BLK | GTTTATTTTGTCACAATCTTACCGAAGCCCTTTAATATCA |
| 100x70 | 7[32]-BLK | TTTAGGACAAATGCTTTAAACAATCAGGTC |
| 100x70 | 7[56]-BLK | ATGCAGATACATAACGGGAATCGTCATAAATAAAGCAAAG |
| 100x70 | 7[96]-BLK | TAAGAGCAAATGTTTAGACTGGATAGGAAGCC |
| 100x70 | 8[111]-BLK | AATAGTAAACACTATCATAACCCTCATTGTGA |

| | | |
|---|---|---|
| 100x70 | 8[143]-BLK | CTTTTGCAGATAAAAACCAAAATAAAGACTCC |
| 100x70 | 8[175]-BLK | ATACCCAACAGTATGTTAGCAAATTAGAGC |
| 100x70 | 8[207]-BLK | AAGGAAACATAAAGGTGGCAACATTATCACCG |
| 100x70 | 8[239]-BLK | AAGTAAGCAGACACCACGGAATAATATTGACG |
| 100x70 | 8[271]-BLK | AATAGCTATCAATAGAAAATTCAACATTCA |
| 100x70 | 8[47]-BLK | ATCCCCCTATACCACATTCAACTAGAAAAATC |
| 100x70 | 8[79]-BLK | AATACTGCCCAAAAGGAATTACGTGGCTCA |
| 100x70 | 9[128]-BLK | GCTTCAATCAGGATTAGAGAGTTATTTTCA |
| 100x70 | 9[192]-BLK | TTAGACGGCCAAATAAGAAACGATAGAAGGCT |
| 100x70 | 9[224]-BLK | AAAGTCACAAAATAAACAGCCAGCGTTTTA |
| 100x70 | 9[256]-BLK | GAGAGATAGAGCGTCTTTCCAGAGGTTTTGAA |
| 100x70 | 9[32]-BLK | TTTACCCCAACATGTTTTAAATTTCCATAT |
| 100x70 | 9[64]-BLK | CGGATTGCAGAGCTTAATTGCTGAAACGAGTA |
| 100x70 | 9[96]-BLK | CGAAAGACTTTGATAAGAGGTCATATTTCGCA |

### Supplementary Table 6: List of elongated staple strands

| DNA origami layout [nm] | Name | Sequence | Docking Sequence |
|---|---|---|---|
| 30x20 | Z'-4T-6[39] | AGAGTCCTAGCATATTTAGCCTTTTAGTGTTGTACGTGGACTCCAACGTCAAAGGGC | AGAGTCCTAGCATATTTAGCC |
| 30x20 | Z'-4T-3[88] | AGAGTCCTAGCATATTTAGCCTTTTCCGGGTACTTTCCTGTGTGAAATTCCTGGGGT | AGAGTCCTAGCATATTTAGCC |
| 30x20 | Z'-4T-1[88] | AGAGTCCTAGCATATTTAGCCTTTTGCCTAATGCACTGCCCGCTTTCCA | AGAGTCCTAGCATATTTAGCC |
| 30x20 | Z'-4T-2[39] | AGAGTCCTAGCATATTTAGCCTTTTGCTGATTGCAAGCGGTCCACGCTGGTTTGAGCTT | AGAGTCCTAGCATATTTAGCC |
| 30x20 | Z'-4T-5[88] | AGAGTCCTAGCATATTTAGCCTTTTAGGGGGATGGGTTTTCCCAGTCACGAGGATCC | AGAGTCCTAGCATATTTAGCC |
| 30x20 | Z'-4T-4[39] | AGAGTCCTAGCATATTTAGCCTTTTAAATCCTGTTATAAATCAAAAGAATAGCCGTGCG | AGAGTCCTAGCATATTTAGCC |
| 30x20 | 1[72]-4T-V' | GTGTAAAGGTTATCCGCTCACAATCTGCAGGTTTTTGTGGAGTAGTGTCATGT | GTGGAGTAGTGTCATGT |
| 65x54 | 9[80]-4T-V' | CAAAAATAAAAGAGGACAGATGAAGAACTGACTTTTGTGGAGTAGTGTCATGT | GTGGAGTAGTGTCATGT |
| 65x54 | 16[95]-4T-V' | GTATCGGTGCTGTTTCCTGTGTGACGTAATCATTTTGTGGAGTAGTGTCATGT | GTGGAGTAGTGTCATGT |
| 65x54 | 5[80]-4T-V' | CGGAGACATTCATCAGTTGAGATTATTACAGGTTTTGTGGAGTAGTGTCATGT | GTGGAGTAGTGTCATGT |
| 65x54 | 15[112]-4T-V' | TTTCTTAACCGCTTTTGCGGGATCCGAGGGTATTTTGTGGAGTAGTGTCATGT | GTGGAGTAGTGTCATGT |
| 65x54 | 15[112]-4T-X' | TTTCTTAACCGCTTTTGCGGGATCCGAGGGTATTTTGCCAAGATAGACAGAAG | GCCAAGATAGACAGAAG |
| 65x54 | 8[95]-4T-V' | AGAACGAGCAATCATATGTACCCCCGTAAAACTTTTGTGGAGTAGTGTCATGT | GTGGAGTAGTGTCATGT |
| 65x54 | Z'-10[47] | AGAGTCCTAGCATATTTAGCCTTTTTAAATGTGTTGTTAAATCAGCTCAAGCCCCAA | AGAGTCCTAGCATATTTAGCC |
| 65x54 | Z'-11[128] | AGAGTCCTAGCATATTTAGCCTTTTCAACGGAGGCACCAACCTAAAACGTACAGAGG | AGAGTCCTAGCATATTTAGCC |
| 65x54 | Z'-14[47] | AGAGTCCTAGCATATTTAGCCTTTTACGCCAGGCTGTTGGGAAGGGCGATCGCACTC | AGAGTCCTAGCATATTTAGCC |
| 65x54 | Z'-15[128] | AGAGTCCTAGCATATTTAGCCTTTTATACCGATAAAATCTCCAAAAAAAAACAACTT | AGAGTCCTAGCATATTTAGCC |
| 65x54 | Z'-18[47] | AGAGTCCTAGCATATTTAGCCTTTTTGCGTATTGCGTTGCGCTCACTGCCAATTCCA | AGAGTCCTAGCATATTTAGCC |
| 65x54 | Z'-19[128] | AGAGTCCTAGCATATTTAGCCTTTTAACCCATGTAGTACCGCCACCCTCAGTACCAG | AGAGTCCTAGCATATTTAGCC |
| 65x54 | Z'-2[47] | AGAGTCCTAGCATATTTAGCCTTTTGCGAGCTGAATTCTGCGAACGAGTATGCTGTA | AGAGTCCTAGCATATTTAGCC |
| 65x54 | Z'-3[128] | AGAGTCCTAGCATATTTAGCCTTTTCAATACTGGACGATAAAAACCAAAAACTAATG | AGAGTCCTAGCATATTTAGCC |
| 65x54 | Z'-6[47] | AGAGTCCTAGCATATTTAGCCTTTTATAAATTAGAGTAATGTGTAGGTAATACTTTT | AGAGTCCTAGCATATTTAGCC |
| 65x54 | Z'-7[128] | AGAGTCCTAGCATATTTAGCCTTTTCATTGTGAGCTCATTCAGTGAATACGCATAGG | AGAGTCCTAGCATATTTAGCC |
| 100x70 | 5[160]-4T-V' | GCAAGGCCTCACCAGTAGCACCATGGGCTTGATTTTGTGGAGTAGTGTCATGT | GTGGAGTAGTGTCATGT |
| 100x70 | 9[160]-4T-V' | AGAGAGAAAAAAATGAAAATAGCAAGCAAACTTTTTGTGGAGTAGTGTCATGT | GTGGAGTAGTGTCATGT |
| 100x70 | 13[160]-4T-V' | GTAATAAGTTAGGCAGAGGCATTTATGATATTTTTTGTGGAGTAGTGTCATGT | GTGGAGTAGTGTCATGT |
| 100x70 | 17[160]-4T-V' | AGAAAACAAAGAAGATGATGAAACAGGCTGCGTTTTGTGGAGTAGTGTCATGT | GTGGAGTAGTGTCATGT |
| 100x70 | 12[111]-4T-V' | TAAATCATATAACCTGTTTAGCTAACCTTTAATTTTGTGGAGTAGTGTCATGT | GTGGAGTAGTGTCATGT |
| 100x70 | 12[175]-4T-V' | TTTTATTTAAGCAAATCAGATATTTTTTGTTTTTGTGGAGTAGTGTCATGT | GTGGAGTAGTGTCATGT |
| 100x70 | 18[63]-4T-Z' | ATTAAGTTTACCGAGCTCGAATTCGGGAAACCTGTCGTGCTTTTAGAGTCCTAGCATATTTAGCC | AGAGTCCTAGCATATTTAGCC |
| 100x70 | 4[63]-4T-Z' | ATAAGGGAACCGGATATTCATTACGTCAGGACGTTGGGAATTTTAGAGTCCTAGCATATTTAGCC | AGAGTCCTAGCATATTTAGCC |
| 100x70 | 18[127]-4T-Z' | GCGATCGGCAATTCCACACAACAGGTGCCTAATGAGTGTTTTAGAGTCCTAGCATATTTAGCC | AGAGTCCTAGCATATTTAGCC |
| 100x70 | 4[127]-4T-Z' | TTGTGTCGTGACGAGAAACACCAAATTTCAACTTTAATTTTTAGAGTCCTAGCATATTTAGCC | AGAGTCCTAGCATATTTAGCC |
| 100x70 | 18[191]-4T-Z' | ATTCATTTTTGTTTGGATTATACTAAGAAACCACCAGAAGTTTTAGAGTCCTAGCATATTTAGCC | AGAGTCCTAGCATATTTAGCC |
| 100x70 | 4[191]-4T-Z' | CACCCTCAGAAACCATCGATAGCATTGAGCCATTTGGGAATTTTAGAGTCCTAGCATATTTAGCC | AGAGTCCTAGCATATTTAGCC |
| 100x70 | 18[255]-4T-Z' | AACAATAACGTAAAACAGAAATAAAAATCCTTTGCCCGAATTTTAGAGTCCTAGCATATTTAGCC | AGAGTCCTAGCATATTTAGCC |
| 100x70 | 4[255]-4T-Z' | AGCCACCACTGTAGCGCGTTTTCAAGGGAGGGAAGGTAAATTTTAGAGTCCTAGCATATTTAGCC | AGAGTCCTAGCATATTTAGCC |

### Supplementary Table 7: List of DNA-PAINT staple strands

| DNA origami layout [nm] | Name | Sequence | DNA-PAINT Sequence |
|---|---|---|---|
| 65x54 | 14[63]-6T-P1 | GCGATTAAGTACCGAGCTCGAATTAATTGTTATTTTTT ATACATCTA | ATACATCTA |
| 65x54 | 19[112]-6T-P1 | TAGCAAGCCGATCTAAAGTTTTGTGTATGGGATTTTTT ATACATCTA | ATACATCTA |
| 65x54 | 2[63]-6T-P1 | TATTTTCATTAAGCAATAAAGCCTACATTATGTTTTTT ATACATCTA | ATACATCTA |
| 65x54 | 7[112]-6T-P1 | TAATTTCACTAACGGAACAACATTTAGGAATATTTTTT ATACATCTA | ATACATCTA |
| 100x70 | 23[32]-6T-P1 | CAAATCAAGTTTTTTGGGGTCGAAACGTGGATTTTTTATACATCTA | ATACATCTA |
| 100x70 | 4[47]-6T-P1 | GACCAACTAATGCCACTACGAAGGGGGTAGCATTTTTTATACATCTA | ATACATCTA |
| 100x70 | 23[224]-6T-P1 | GCACAGACAATATTTTTGAATGGGGTCAGTATTTTTTATACATCTA | ATACATCTA |
| 100x70 | 4[239]-6T-P1 | GCCTCCCTCAGAATGGAAAGCGCAGTAACAGTTTTTTTATACATCTA | ATACATCTA |

### Supplementary Table 8: List of modified oligonucleotides

| Designation | Sequence | Modification |
|---|---|---|
| Biotin-TEG-V | ACATGACACTACTCCAC | 5'-Biotin-TEG |
| Biotin-TEG-V-4T-AS635P | ACATGACACTACTCCACTTTT | 5'-Biotin-TEG; 3'- AS635P |
| Biotin-TEG-2T-P3*' | TTTCTTCATTA | 5'-Biotin-TEG |
| Biotin-TEG-V-2T-P3*' | ACATGACACTACTCCAC TTTCTTCATTA | 5'-Biotin-TEG |
| AF647-4T-X | TTTTCTTCTGTCTATCTTGGC | 5'-AF647 |
| DY548-4T-V | TTTTACATGACACTACTCCAC | 5'-DY548 |
| Z-TEG-Cholesterol | GGCTAAATATGCTAGGACTCT | 3'-Cholesterol |
| Z-TEG-Biotin | GGCTAAATATGCTAGGACTCT | 3'-TEG-Biotin |
| P1-Cy3b | CTAGATGTAT | 3'-Cy3b |
| P3-Cy3b | GTAATGAAGA | 3'-Cy3b |
| P3*'-2T-Z'-4T-TEG-Biotin | TCTTCATTATTAGAGTCCTAGCATATTTAGCCTTTT | 3'TEG-Biotin |

**Supplementary Table 9: DNA-PAINT imaging parameters for divalent DNA origami platforms**. In divalent DNA origami platforms ($M_{div}$, $L_{div}$), biotinylated TCR ligands were replaced by biotinylated DNA-PAINT sequences (Biotin-TEG-2T-P3') and imaged with P3-Cy3B. Additionally, four staple strands at the corners were extended with P1' DNA-PAINT docking strands for barcoding and imaged with P1-Cy3B by Exchange-PAINT. DNA origami were immobilized on neutravidin-coated glass slides and imaged under following conditions:

| Layout | Imager strand & Concentration (nM) | Frames (n) | Integration Time (ms) | Laser power density ($W/cm^2$) | Setup | Analyzed DNA origami (n) |
|---|---|---|---|---|---|---|
| **$M_{div}$ 10nm** | 8nm P1-Cy3B; 10nM P3-Cy3B | 20,000; 20,000 | 150, 200 | 1,038 | 1 | 49 |
| **$M_{div}$ 20nm** | 8nm P1-Cy3B; 10nM P3-Cy3B | 15,000; 15,000 | 150, 200 | 1,038 | 2 | 49 |
| **$M_{div}$ 30nm** | 8nm P1-Cy3B; 10nM P3-Cy3B | 15,000; 15,000 | 150, 150 | 1,038 | 2 | 49 |
| **$L_{div}$ 10nm** | 8nm P1-Cy3B; 10nM P3-Cy3B | 20,000; 20,000 | 150, 200 | 1,038 | 2 | 100 |
| **$L_{div}$ 20nm** | 8nm P1-Cy3B; 10nM P3-Cy3B | 15,000; 15,000 | 150, 200 | 1,038 | 2 | 100 |
| **$L_{div}$ 30nm** | 8nm P1-Cy3B; 10nM P3-Cy3B | 15,000; 15,000 | 150, 150 | 1,038 | 2 | 100 |

**Supplementary Table 10: DNA-PAINT imaging parameters for T-cell experiments.** For DNA-PAINT, biotinylated oligos were extended with P3' DNA-PAINT docking sites at their 3' end and imaged using Cy3B-labeled imager strands (P3-Cy3B). Additionally, four staple strands at the corners were extended with P1' DNA-PAINT docking strands for barcoding and imaged with P1-Cy3B by Exchange-PAINT to verify the integrity of DNA origami platforms upon T-cell interaction as well as to ensure that derived ligand positions could be allocated to the nanoplatforms. All experiments were performed using super-resolution microscope 1 under following conditions:

| Construct | Imager strand concentration (nM) | Frames (n) | Integration Time (ms) | Laser power density (W/cm$^2$) | Analyzed cells (n) | Number of ligands (n) |
|---|---|---|---|---|---|---|
| **DNA-H57** | 10 nM P3-Cy3B | 15,000 | 200 | 1,038 | n.a. | n.a. |
| **M-H57** | 8 nM P1-Cy3B<br>10 nM P3-Cy3B | 15,000<br>15,000 | 100<br>150 | 1,038 | 21 | 2,390 |
| **L-H57** | 8 nM P1-Cy3B<br>10 nM P3-Cy3B | 15,000<br>15,000 | 100<br>100 | 1,038 | 17 | 1,270 |
| **M-647** | 8 nM P1-Cy3B<br>10 nM P3-Cy3B | 15,000<br>15,000 | 100<br>150 | 1,038 | 22 | 627 |
